# p38γMAPK delays myelination and remyelination and is abundant in multiple sclerosis lesions

**DOI:** 10.1101/2023.01.04.522734

**Authors:** LN Marziali, Y Hwang, M Palmisano, A Cuenda, FJ Sim, C Volsko, R Dutta, B Trapp, L Wrabetz, ML Feltri

## Abstract

Multiple Sclerosis is a chronic inflammatory disease in which disability results from the disruption of myelin and axons. During the initial stages of the disease, injured myelin is replaced by mature myelinating oligodendrocytes that differentiate from oligodendrocyte precursor cells. However, myelin repair fails in secondary and chronic progressive stages of the disease and with aging, as the environment becomes progressively more hostile. This may be attributable to inhibitory molecules in the multiple sclerosis environment including activation of the p38MAPK family of kinases.

We explored oligodendrocyte precursor cell differentiation and myelin repair using animals with conditional ablation of p38MAPKγ from oligodendrocyte precursors.

We found that p38γMAPK ablation accelerated oligodendrocyte precursor cell differentiation and myelination. This resulted in an increase in both the total number of oligodendrocytes and the migration of progenitors *ex vivo* and faster remyelination in the cuprizone model of demyelination/remyelination. Consistent with its role as an inhibitor of myelination, p38γMAPK was significantly downregulated as oligodendrocyte precursor cells matured into oligodendrocytes. Notably, p38γMAPK was enriched in samples of leukocortical multiple sclerosis lesions from patients, which represent areas of failed remyelination.

Our data suggest that p38γ could be targeted to improve myelin repair in multiple sclerosis.

## Introduction

Myelin is the lipid-rich insulating layer that wraps axons, providing trophic support and ensuring rapid propagation of the electrical impulses that underlie nervous system function. In the CNS, myelin is produced by mature oligodendrocytes (OLs) that arise from oligodendrocyte progenitor cells (OPCs). In rodents, OPCs develop during late embryonic and early postnatal stages, proliferate, and migrate towards regions that will be myelinated later on.^1–3^ There is also a pool of OPCs that remain in an undifferentiated state that are responsible for adaptive myelination in the adult (e.g., during neuronal plasticity or motor learning) (reviewed by Bergles and Richardson^4^) or to replace injured myelin (remyelination) (reviewed by Cayre et al.^5^). In these instances, OPCs proliferate^6^ and migrate before differentiating into mature OLs to ultimately produce new myelin.^7^

Multiple sclerosis is a chronic inflammatory disease that affects myelin in the CNS, leading to demyelination (in plaques or lesions), axonal degeneration, and disability. In early stages of the disease, primary demyelination is followed by remyelination, but this fails in secondary and chronic progressive stages. The failure of OPCs to differentiate into mature myelinating OLs is one well-accepted cause of impaired remyelination^8^ (reviewed by Franklin and ffrench-Constant^9^ and Bebo et al.^10^). Factors that can favour remyelination in an inflammatory context are the target of much research (reviewed by Kotter^11^). Several myelination inhibitors including LINGO-1 and the muscarinic receptor M1 have been identified^12–14^ and their modulation led to partly successful clinical trials in early Multiple Sclerosis (reviewed by Wooliscroft^15^). While these outcomes are encouraging, it is becoming clear that efficient repair will require targeting multiple factors that must be active in the inflammatory environment of Multiple Sclerosis, within a specific window of opportunity that narrows during the course of the disease (reviewed by Kotter^11^).

Among the many extracellular and intracellular cues that regulate OPC differentiation (reviewed by Wheeler and Fuss^16^ and Galloway and Moore^17^) are mitogen-activated protein kinases (MAPKs), of which there are three canonical families in mammals: the extracellular signal-regulated kinases (ERKs), the c-Jun N-terminal kinases 1/2/3 (JNK1/2/3), and the p38 MAP kinases (p38MAPKs) (reviewed by Morrison^18^). Members of the p38 family are activated by growth factors during development ^19–22^ (reviewed by Huang et al.^23^) but also by stress signals, inflammatory cytokines (e.g., interleukins 1 and 17 and tumour necrosis factor alpha), oxidative stress, UV irradiation, and hypoxia–ischemia (reviewed by Cuenda and Rousseau^24^).

There are four known isoforms of p38 (α, β, γ, and δ), and all are expressed by different cell types in the brain, including neurons, glia, and endothelial cells.^25–28^ In the context of OLs, p38 is often equated with p38α and p38ß.^26,29^ Inhibition of p38α/ß limits OPC differentiation^26^ and developmental myelination; ^30,31^ however, p38α appears to inhibit remyelination after injury^32^ and mice null for p38ß mice have no obvious CNS phenotype.^25^. Less is known about the minor isoforms, p38γ and p38δ (reviewed by Cuenda and Sanz-Esquerro^33^), but their roles oppose those of the α and β isoforms in the proliferation of breast cancer cells^34^ and differentiation of skeletal muscle cells.^35,36^ We performed studies *in vitro* and *in vivo* demonstrating a key role of the γ isoform of p38 in the differentiation of OPCs, with implications in developmental myelination and remyelination. The data we present here suggest that p38γ could be a novel target for therapeutic strategies to enhance remyelination in Multiple Sclerosis.

## Materials and methods

### Animal models

The protocols for all experiments involving animals were approved by the Institutional Animal Care and Use Committees at the University at Buffalo and were performed in accordance with the National Institutes of Health’s Guide for the Care and Use of Laboratory Animals. Mice were backcrossed to the C57BL/6 strain to achieve ≥99% congenicity. p38γ null (p38γ^-/-^) and p38γ flox (p38γ^f/f^) mice have been described elsewhere^37^ and were a gift from A. Cuenda. NG2 CreER^T2^ (NG2 Cre) [B6.Cg-Tg(Cspg4-cre/Esr1*)BAkik/J; stock number 008538] and tdTomato [B6.Cg-Gt(ROSA)26Sortm9(CAG-tdTomato)Hze/J; stock number 007909] mice were purchased from The Jackson Laboratory. Transgenic mice were genotyped by PCR using tail genomic DNA according to the protocols of The Jackson Laboratory (tdTomato), Sabio et al.^37^ (p38γ^-/-^ and p38γ^f/f^) or an in-house protocol (NG2 Cre) (see Supplementary Table 1).

To ablate p38γ from OPCs during development, p38γ^f/f^-NG2 Cre animals (and p38γ^f/f^ control littermates) received daily 20-μl intraperitoneal injections of a solution containing 2.5-mg/ml tamoxifen (Tx) (Sigma-Aldrich; T5648) in corn oil (50 μg/pup).

Demyelination/remyelination was induced by using cuprizone (CPZ).^38,39^ Eight-week-old animals were fed a diet containing 0.2% (w/w) CPZ (Envigo; TD.140803) for 7 weeks to induce demyelination followed by normal diet for 2 weeks to allow remyelination. Female and male mice respond similarly to CPZ insult in the C57BL/6 background^40^ and thus both were used in these experiments. p38γ ablation was induced with daily intraperitoneal injections of 1 mg Tx (100 μl of a 10-mg/ml solution) for 10 days starting 5 days before the end of CPZ feeding.

### Reverse transcriptase quantitative PCR

Total RNA was isolated from cells with TRIzol reagent (Thermo Fisher), and 1 μg of RNA reverse transcribed using SuperScript III reverse transcriptase kit (Thermo Fisher; 18080093) with oligo(dT) primers. Quantitative PCR was performed with SYBR green PCR master mix (Thermo Fisher) and the primers listed in Supplementary Table 1. Transcription levels were analysed using the efficiency-corrected comparative threshold cycle method.

### Immunocytochemistry and immunohistochemistry

For immunocytochemistry, cells were fixed for 20 min at room temperature (RT) with 2% paraformaldehyde (PFA) in PBS. The cells were rinsed with PBS and blocked for 2 h at RT with 5% foetal bovine serum (FBS) in PBS with 0.2% Triton X-100. Cells were incubated overnight at 4°C with primary antibodies in PBS with 2% FBS and 0.1% Triton X-100 (antibodies listed in Supplementary Table 2). After three washes, the cells were incubated with 4’,6-diamidino-2-phenylindol (DAPI; 1 μg/ml) and the corresponding Alexa Fluor-conjugated secondary antibodies (1:500; Jackson ImmunoResearch Laboratories) for 2 h at RT. Cells were washed three times and mounted on slides with antifading agent.

For immunohistochemistry, animals were perfused with 4% PFA in PBS, and their brains postfixed with 4% PFA in PBS overnight at 4°C. The tissues were cryoprotected by immersion in 30% sucrose, and 30-μm-thick coronal slices were cut with a Leica CM1850 cryostat at ~0.62 mm from bregma according to the P56 mouse Allen Brain Atlas. Slices were incubated under free-floating conditions with antibodies as described above for immunocytochemistry. Sections were imaged with a Zeiss Apotome microscope and analysed using ImageJ software.

### Immunoblotting

Total proteins from cells or tissue were extracted with RIPA buffer supplemented with protease and phosphatase inhibitors. Proteins (1–2 μg) were loaded into 7.5–15% in-house-made gradient sodium dodecyl sulphate (SDS) polyacrylamide gels and electrotransferred onto polyvinylidene difluoride membranes. The membranes were blocked with 5% non-fat milk in PBS with 0.2% Tween 20 for 2 h at RT. Membranes were incubated for 2 h at RT with antibodies diluted in the blocking solution (Supplementary Table 2), washed, and incubated for another 2 h at RT with the corresponding peroxidase-conjugated secondary antibodies. The blots were washed and the labelled protein bands were visualized by using Pierce ECL reagent and a ChemiDoc XRS+ system (Bio-Rad).

### *In situ* hybridization

To produce the p38γ antisense probes, base pairs 1–258 of the mouse mRNA transcript (NM_013871.3; forward primer, 5’-AGTGGGCCGGGAG-3’; reverse, primer 5’-CTTGCGGGCGGGT-3’) or 98–158 of the human mRNA transcript (NM_001303252.2; forward primer, 5’-GGTGGTTCTAGAGGGCACCAACTCAGG-3’; reverse primer, 5’-GGTGGTGGGCCCGAGCTACAAAAGGGTCTATTTCCT-3’) were amplified and cloned into pBluescript SK(+/-) phagemid vectors. The plasmids were linearized with SapI, and *in vitro* transcription was performed with T7 RNA polymerase (NEB; M0251) and a digoxigenin RNA labelling kit (Roche; 11175025910) according to the manufacturer’s instructions.

To perform *in situ* hybridization (ISH), brain sections were treated with proteinase K (20 μg/ml) in Tris-EDTA buffer for 15 min at 37°C, washed with PBS, and fixed with 4% PFA for 15 min at RT. Samples were then incubated in 5× SSC (1× SSC is 0.15 M NaCl plus 0.015 M sodium citrate), 50% formamide, and 1% SDS for 10 min at 60°C followed by hybridization with 0.5 ng/ml of p38γ antisense probe in the same solution for 2 h at 60°C. The sections were washed twice for 10 min each at 60°C with 2× SSC, 50% formamide, and 0.1% SDS. After washing with maleic acid buffer 3 times for 5 min at RT, sections were incubated with 1× blocking reagent (Roche; 11096176001) for 2 h at RT. Sections were then incubated overnight at 4°C with anti-digoxigenin-alkaline phosphatase antibodies (1:5000; Roche; 11093274910; AB_514497) diluted in blocking solution. The sections were washed and incubated with NBT/BCIP (nitroblue tetrazolium/5-bromo-4-chloro-3-indolylphosphate) substrate for at least 6 h at RT.

In some cases, ISH was followed by immunohistochemistry. Endogenous peroxidases were blocked by incubating the sections in 1% hydrogen peroxidase for 10 min at RT. After blocking for 2 h at RT with 5% FBS in PBS with 0.2% Triton X-100, the sections were incubated overnight at 4°C with primary antibodies diluted in PBS with 2% FBS and 0.1% Triton X-100 (see Supplementary Table 2). After three washes, the sections were incubated with the corresponding peroxidase-conjugated secondary antibodies (1:1000) for 2 h at RT. After final washing, the labelled antibodies were developed with a Dako Liquid DAB+ substrate chromogen system (Dako; GV82511-2).

### Cortical OPC cultures

OPCs were isolated from the cortices of postnatal day 3-5 (P3–P5) mice by immunopanning as described previously by Emery and Dugas^41^ with modifications. The cortices were dissociated into a single cell suspension using a neural tissue dissociation kit (Miltenyi Biotec; 130-092-628). Microglia were eliminated by panning on plates coated with *Griffonia* (*Bandeiraea*) *simplicifolia* lectin 1 (5 μg/ml) (Vector Laboratories; L-1100). OPCs were purified on plates coated with an antibody to platelet-derived growth factor receptor alpha (PDGFrα) (1:4000) (Supplementary Table 2) and then seeded on coverslips coated with poly-D-lysine (20 μg/ml) (Sigma-Aldrich; P0899) and incubated at 37°C and 10% CO_2_ in Sato medium: Dulbecco’s modified Eagle medium (Thermo Fisher; 11965092) containing 0.1 mg/ml bovine serum albumin (Sigma-Aldrich; A4161), 0.1 mg/ml transferrin (Sigma-Aldrich; T1147), 16 μg/ml putrescine (Sigma-Aldrich; P5780), 60 ng/ml progesterone (Sigma-Aldrich; P8783), 40 ng/ml sodium selenite (Sigma-Aldrich; S5261), 10 ng/ml D-biotin (Sigma-Aldrich; B4639), 5 μg/ml insulin, 1 mM sodium pyruvate (Thermo Fisher; 11360-070), 5 μg/ml *N*-acetyl-L-cysteine (Sigma-Aldrich; A8199), 1× trace elements B (Cellgro; 99-175-CI), 4.2 ng/ml forskolin (Sigma-Aldrich; F6886), and 1× B27 (Thermo Fisher Scientific; 17504044). OPCs were kept proliferating for 48–72 h by adding 10 ng/ml ciliary neurotrophic factor (CNTF) (PeproTech; 450-13), 1 ng/ml Neurotrophin 3 (NT3) (PeproTech; 450-03), and 20 ng/ml PDGF-AA (PeproTech; 100-13A) to the basal medium. OPCs were then allowed to differentiate for 24, 48, or 72 h in the Sato medium described above also containing 10 ng/ml CNTF, 15 nM T3 (Sigma-Aldrich T6397), and 2.5 μg/ml L-ascorbic acid (Sigma-Aldrich; A4544).

### EdU experiments

Proliferating cells were labelled with the thymidine analogue 5-ethynyl-2’-deoxyuridine (EdU) (Sigma; 900584) at a dose of 50 mg/kg body weight *in vivo* or 1 μM *in vitro*. For the *in vivo* experiments, pups received one intraperitoneal injection to evaluate proliferation. For the *in vitro* experiments, EdU was added to the culture medium 1 h before fixation.

EdU incorporation was visualized by using click chemistry. Briefly, tissue sections or cells were incubated for 30 min at RT with a solution containing 8 μM sulfo-Cy3-azide (Lumiprobe; B1330) or 8 μM Alexa Fluor 488-azide (Cedarlane; CLK-1275-1), 2 mM CuSO_4_·5H_2_O, and 20 mg/ml ascorbic acid.

### Mouse p38γ cloning and KETAL-del and kinase domain mutagenesis

Wild type (WT) and mutant variants of mouse p38γ were cloned into a pULTRA plasmid,^42^ a gift from Malcolm Moore (Addgene plasmid no. 24129; http://n2t.net/addgene:24129; RRID: Addgene_24129). To obtain WT p38γ, 1 μg of mouse skeletal muscle RNA (extracted with TRIzol reagent) was reverse transcribed using oligo(dT) primers and the SuperScript III reverse transcriptase kit. The coding sequence of mouse p38γ mRNA (NM_013871.3) was amplified by PCR using Phusion high-fidelity DNA polymerase (Thermo Fisher; F530S) and the following primers: forward, 5’-GGTGG<u>TTCTAGA</u>ATGAGCTCCCCGCCACCCGCCCGCAAG-3’; reverse, 5’-GGTGGT<u>GAATTC</u>TCACAGAGCCGTCTCCTTTGGAACTCTGGCTCCTAGC-3’, which include restriction enzyme sites (underlined sequences; forward, XbaI; reverse, EcoRI) to ensure proper orientation upon cloning into the pULTRA plasmid. The PCR cycling conditions were 98°C for 2 min followed by 30 cycles of 98°C for 20 s and 72°C for 1 min.

To obtain a mouse p38γ mutant that lacks the KETAL binding motif (KETAL-del), the same strategy was used but with a reverse primer (with an EcoRI restriction site) that specifically binds upstream of the last 15 nucleotides of p38γ mRNA (coding the KETAL motif): 5’-GGTGGT<u>GAATTC</u>TCATGGAACTCTGGCTCCTAGCTGCCTAGGAGGCTTGA-3’.

The kinase activity of p38γ is ablated with a D171A mutation.^43^ Thus, targeted mutagenesis was performed on the cloned WT p38γ by using the Quick-Change Lightning site-directed mutagenesis kit (Agilent Technologies; 210518) to create the kinase dead mutant.

All constructs were verified by sequencing performed at the Sanger sequencing facility of Roswell Park Comprehensive Cancer Center (Buffalo, NY).

### Lentiviral particle production and OPC transduction

The ViraPower lentiviral packaging mix (Thermo Fischer; K497500) was used to package the p38γ constructs described above. HEK293T cells plated in 10-cm dishes were transfected with 3 μg of the construct plasmids along with 3 μg of each of the packaging plasmids using Lipofectamine 2000 transfection reagent (Thermo Fischer; 11668019). The medium was replaced after 16 h, and viral supernatants collected after 48 h. Lentiviral particles were concentrated by ultracentrifugation according to Kutner et al.^44^ The titres of the lentiviral concentrates were determined by using a qPCR lentivirus titre kit (abm; LV900).

To overexpress p38γ constructs, OPCs were infected with lentivirus at a multiplicity of infection of 10 in the presence of 1 μM Polybrene (EMD Millipore; TR-1003-G). The medium was replaced 24 h later, and OPCs were allowed to recover for another 48 h.

### p38α inhibition

p38α inhibition was induced via a previously characterised inhibitor, SB203580.^45^ For *in vitro* experiments, OPCs were treated with 2 μM SB203580 (or the vehicle, dimethyl sulfoxide) during differentiation.^26^

### RNA-sequencing

Total RNA was isolated with TRIzol from cortical OPCs isolated from P7 mouse pups and quantified with a RiboGreen assay (Thermo Fisher). The quality of the samples was checked with an Agilent Bioanalyzer 2100 RNA 6000 Nano chip (Agilent). cDNA libraries were prepared with the Illumina TruSeq RNA sample preparation kit (Illumina). Briefly, mRNA with poly(A) tails was isolated, cleaved, and first-strand reverse transcribed to cDNA using the SuperScript III reverse transcriptase kit and random primers; second-strand cDNA synthesis was performed using the second strand master mix supplied with the kit. After end repair, addition of a single A base, and ligation with adapters, the cDNA was enriched and purified by PCR to create the final cDNA library according to the manufacturer’s instructions. cDNA libraries were quantified with a PicoGreen assay (Thermo Fisher) and a library quantification kit (Kapa Biosystems). To confirm the quality and size of the cDNA library, an Agilent Bioanalyzer 2100 DNA 7500 chip was used. The cDNA libraries were then normalized, pooled, and paired-end sequenced (100 standard cycles) using the Illumina HiSeq 2500 at the UB Genomics and Bioinformatics Core facility (Buffalo, NY). Sequences were aligned to the UCSC Mm10 mouse genome using tophat (v2.0.13), and counts per gene were determined using htseq (v0.6.1). R/Bioconductor was used for subsequent analyses. After loading read counts by using DESeq2, edgeR was used for differential expression analysis of read counts; genes with less than one count across all samples were excluded. Pathway analysis was performed in edgeR using Gene Ontology (GO) enrichment analysis, KEGG pathway enrichment analysis, and gene set enrichment analysis of broad C2 gene sets (v5) using roast. Heat maps were plotted using the log2(counts per million).

### Human Multiple Sclerosis samples

Previously characterized brain sections from individuals with Multiple Sclerosis were provided by Dr. Dutta.^46,47^ Brain samples were obtained as part of the tissue procurement program approved by the Cleveland Clinic Institutional Review Board (IRB) with informed consent for all tissue donors. The de-identified samples were exempt from IRB approval at the University at Buffalo. As described by Dutta et al.,^47^ the patients’ brains were removed according to a rapid autopsy protocol, sliced (1□ cm thick), fixed in 4% PFA, embedded in paraffin, and sliced into 15-μm sections.

### *Ex vivo* acute slice culture and time-lapse acquisition

Brain acute slice cultures were performed according to De Simoni and Yu.^48^ Briefly, 150-μm coronal brain sections were sliced from P3 triple transgenic p38γ^f/f^ (or p38γ^f/+^)/tdTomato^+^/NG2 Cre^+^ mice with a Leica VT1000S vibratome. The sections were plated on Corning Transwell culture plate inserts (Sigma-Aldrich; CLS3492) and cultured in minimal essential medium with 1× GlutaMAX, Earle’s balanced salt solution, and 25% horse serum at 37°C with 5% CO_2_. After 24 h, 1 μM of 4 hydroxy-Tx (4-OH-Tx) was added, and slices were cultured for another 48 h. Culture medium was replaced and PDGF-AA (20 ng/ml) and basic fibroblast growth factor (bFGF) (10 ng/ml) were added 24 h before running the time-lapse experiments according to the procedure described by Cheli et al.^49^ The sections were placed on a spinning disc confocal inverted microscope (Olympus; IX83-DSU) equipped with an incubating chamber and a motorized stage. Images of red tdTomato^+^ cells were collected every 6 min for 24 h. Cell migration speed and distances were analysed by tracing at least 6 cells per slice using the motion tracking function in MetaMorph software (Molecular Devices).

### Electron microscopy analysis

Animals were perfused with PBS followed by 2.5% glutaraldehyde/4% PFA in 0.12 M phosphate buffer (pH 7.4). Tissue was collected and postfixed with 2% glutaraldehyde in phosphate buffer for at least 48 h at 4°C. The samples were stained with 1% osmium tetroxide for 2 h at RT, dehydrated, and embedded into epoxy resin (Sigma-Aldrich; 45359-1EA-F). Semithin (500 nm) and ultrathin (90 nm) sections were obtained with a Leica EM UC7 ultramicrotome. Ultrathin sections were stained with uranyl acetate and lead citrate and imaged with an FEI Tecnai G2 Spirit BioTwin transmission electron microscope. Images were analysed with ImageJ software. G-ratio analysis was conducted by drawing a circle around at least 100 imaged axons and fibres per animal. G-ratios were plotted against axon diameters according to log-linear fits.^50^

### Statistical analysis

Blinding to experimental conditions was performed by assigning numbers to the mice used in the study and revealing the genotypes only after performing all the quantifications. The tissue samples from patients were identified as control or multiple sclerosis samples after analysis. No estimations of sample number were performed, but at least four biological replicates were included per condition, and each experiment was repeated at least two times. GraphPad Prism 9.1.1 software was used to perform two-tailed Student’s *t* tests to compare two groups/time points and one-way ANOVAs followed by Newman–Keuls post-tests to compare more than two groups/time points. Statistical significance was set at a *P* value of <0.05.

### Data availability

All data are presented in the results or supplemental material.

## Results

### p38γ is downregulated during OPC differentiation and myelination

Myelination of the rodent CNS takes places during the first weeks of postnatal life, with peak expression of the genes for myelin basic protein (MBP) and proteolipid protein (PLP) in the brain at P20.^51^ The process is mostly complete by P60.^52,53^ We performed ISH to explore the expression of p38γ during myelination in WT mice by using brain sections taken at P5, P15, and P30. Unlike those from WT mice, sections from p38γ^-/-^ mice showed no ISH signal, thereby validating our probe (**Supplementary Fig. 1A, B**). p38γ was ubiquitously expressed at all three ages in different brain regions, such as cortex, corpus callosum, hippocampus, and caudate putamen, though the levels of expression in cortex and the corpus callosum were much lower at P30 (**Fig. 1A**). This indicates that p38γ is downregulated in regions that become heavily myelinated. We additionally performed immunohistochemistry to identify the cell types that expressed p38γ at P5 and P15. Our results show that p38γ was expressed by PDGFrα^+^ OPCs and Iba1^+^ microglia in the corpus callosum and NeuN neurons in the cortex, but was not detectable in MBP^+^ mature OLs and GFAP^+^ astrocytes in the corpus callosum (**Fig. 1B**). Western blot analysis of protein lysates from the corpus callosum at P5, P10, and P30 confirmed the decrease in p38γ expression during myelination, with concomitant increases in the levels of MBP and PLP at later time points (**Fig. 1C**).

**Figure 1.**
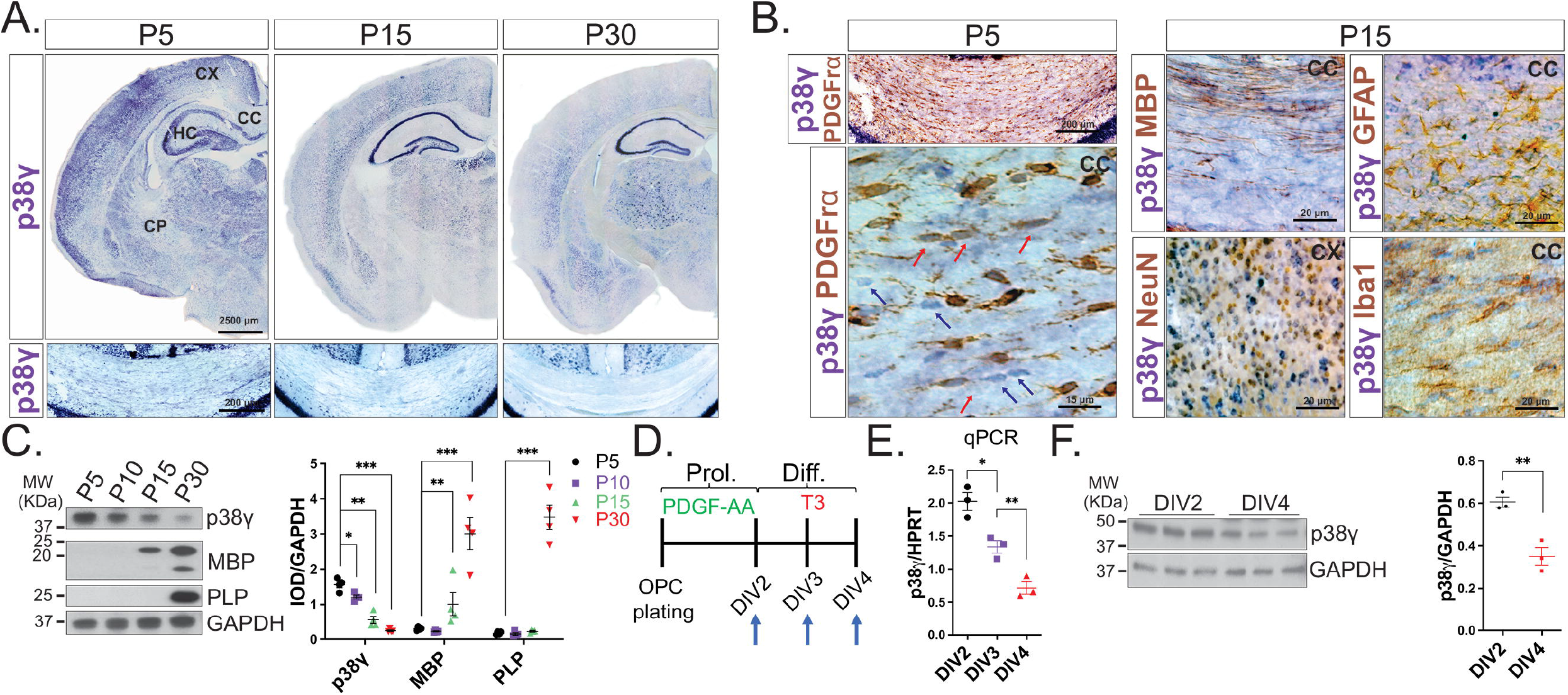
p38γ expression is downregulated during myelination and OPC differentiation. (**A**) ISH of coronal brain sections shows ubiquitous expression of p38γ during early postnatal development and a strong downregulation of p38γ during myelination in the cortex (CX), corpus callosum (CC), and caudate putamen (CP) of WT brains. p38γ expression in the hippocampus (HC) shows no change throughout development. (**B**) ISH for p38γ followed by immunohistochemistry for the OPC marker PDGFrα, mature OL marker MBP, astrocyte marker GFAP, microglial marker Iba1, and neuronal marker NeuN. At P5, p38γ is expressed by PDGFrα^+^ OPCs (red arrows) and other cell types (blue arrows). At P15, p38γ is expressed by NeuN^+^ neurons and Iba1^+^ microglia but not by MBP^+^ mature OLs or GFAP^+^ astrocytes. (**C**) Western blots for p38γ and myelin proteins (MBP and PLP) in lysates from the corpus callosa of WT animals showing that p38γ content decreases during myelination. IOD, integrated optical density. (**D**) Schematic representation of OPC culture procedures. After plating, OPCs were kept proliferating for 2 days (to DIV2) followed by 2 days of differentiation (DIV3–4). (**E**) Quantitative (q)PCR for p38γ in OPCs (DIV2) and mature OLs (DIV4) showing strong reduction in the expression of p38γ in mature OLs. HPRT, hypoxanthine phosphoribosyltransferase. (**F**) Western blots for p38γ in OPCs (DIV2) and mature OLs (DIV4) showing strong reduction in the content of p38γ in mature OLs. GAPDH, glyceraldehyde-3-phosphate dehydrogenase (reference for normalization). Values are expressed as the means ± SEMs. *\*P* < 0.05, *\*\*P* < 0.01, and \*\*\**P* < 0.001 by Student’s *t* test or one-way ANOVA followed by Newman–Keuls multiplecomparisons post-test.

These data suggest that OPCs downregulate p38γ expression as they mature into OLs. To test this, we measured p38γ mRNA and protein levels in OPCs differentiating *in vitro* (**Fig. 1D**). More than 90% of the cells in the culture were Olig2^+^, with 30–45% becoming PLP^+^ mature OLs by 3 and 4 days *in vitro* (DIV3–4) (**Supplementary Fig. 1C**). Quantitative PCR at DIV2, DIV3, and DIV4 showed that p38γ mRNA levels decreased as the OPCs differentiated (**Fig. 1E**). This was confirmed by Western blotting of lysates at DIV2 and DIV4 (**Fig. 1F**). Overall, these data show that p38γ is expressed by OPCs, microglia, and neurons and is downregulated during OPC differentiation and myelination.

### p38γ ablation accelerates OPC differentiation and myelination

p38γ^-/-^ mice do not show a pathological phenotype and can breed normally,^54^ but we detected an increased number of myelinated axons in the corpus callosum at P15 (**Supplementary Fig. 2A**). We examined mice null for the other minor form of p38 (p38δ) and found that p38δ^-/-^ mice do not have increased levels of myelin proteins (MBP, PLP, and myelin-associated glycoprotein) in whole brain lysates at P15, whereas the p38γ^-/-^ mice and double null mice (p38γ^-/-^/δ^-/-^) do; however, levels did not differ by genotype at 3 months (**Supplementary Fig. 2B**). This indicates that accelerated myelination is attributable to the deletion of p38γ. This acceleration occurred in the characteristic caudal-to-rostral pattern in the corpus callosum, with normal myelin thickness and compaction observed at P30 (data not shown). Overall, these data indicate that myelination occurs early, but proceeds normally in the absence of p38γ.

The above-described data showing that p38γ is expressed in OPCs and is inversely associated with their differentiation and with myelination suggests that p38γ functions autonomously in OPCs. To test this, we conditionally deleted p38γ in OPCs *in vivo* by using p38γ^f/f^-NG2 Cre mice (mice carrying the floxed allele for p38γ crossed with NG2 Cre mice). Tx was administered from P2 to P6 to induce Cre-mediated ablation of the gene encoding p38γ in OPCs of animals analysed at P15 and P30 (**Fig. 2A**). This resulted in robust p38γ recombination in genomic DNA (**Fig. 2B**). The tdTomato reporter line showed tdTomato expression in most Olig2^+^ oligodendrocytes in the cortices and corpus callosa of in the brains of mice at P15 (**Fig. 2C**).

**Figure 2.**
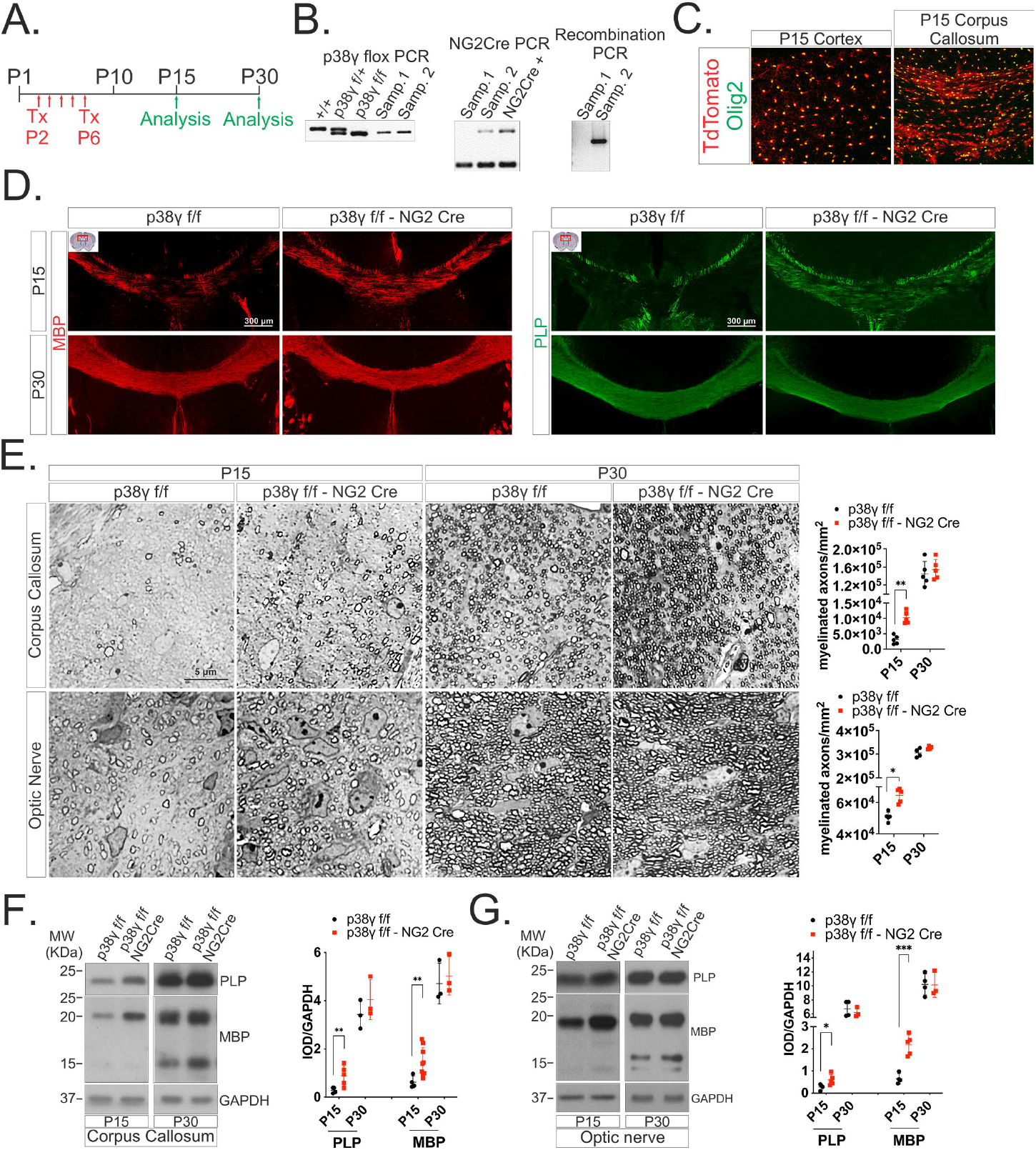
p38γ^f/f^-NG2 Cre mice show premature myelination. (**A**) Schematic representation of Tx injections and analysis time points. (**B**) Gel analysis of PCR products from tail DNA for the genotyping of mice that have 1 or 2 floxed p38γ alleles (p38γ^f/+^ or p38γ^f/f^, respectively) (left) and the NG2 Cre transgene (middle). (Right) Gel analysis from PCR using DNA from the corpus callosa of P15 mice showing that Tx-induced recombination only occurred in sample 2, which was from a mouse that had the floxed p38γ allele and the NG2 Cre transgene. (**C**) Immunostaining for Olig2 in coronal sections from a P15 tdTomato-NG2 Cre mouse treated with Tx showing that NG2 CreER^T2^ tags most Olig2^+^ OLs in the corpus callosum and cortex. (**D**) Immunostaining for MBP and PLP in coronal sections from P15 and P30 mice showing that myelination is prematurely advanced in p38γ^f/f^-NG2 Cre mice. (**E**) Semithin sections of P15 and P30 corpus callosum and optic nerves showing that p38γ^f/f^-NG2 Cre mice have a greater density of myelinated axons at P15 but no difference at P30. (**F**) Western blot analyses of P15 and P30 corpus callosum (**F**) and optic nerve (**G**) lysates showing that p38γ^f/f^-NG2 Cre mice have increased amounts of myelin proteins (MBP and PLP) at P15 but no differences at P30. Values are expressed as the means ± SEMs. \**P* < 0.05, \*\**P* < 0.01 and \*\*\**P* < 0.001 by Student’s *t* tests.

Immunohistochemical analyses showed greater expression of MBP and PLP in corpus callosa of p38γ^f/f^-NG2 Cre mice than in the same regions of control littermates at P15, but similar levels at P30 (**Fig. 2D**). These findings were confirmed by a morphological analysis showing that p38γ^f/f^-NG2 Cre mice have a higher density of normally myelinated axons than control littermates at P15 and similar numbers at P30 (**Fig. 2E**). Western blotting showed that the corpus callosa of p38γ^f/f^-NG2 Cre mice had more MBP and PLP than controls at P15 and no differences at P30 (**Fig. 2F**). Finally, accelerated myelination was also observed in the optic nerves of p38γ^f/f^-NG2 Cre mice (**Fig. 2E, G**).

To further explore the effect of p38γ ablation on accelerated myelination, we analysed OPC differentiation in mice subjected to the same Tx administration protocol and we included an analysis time point at P10 (**Fig. 3A**). We found that p38γ^f/f^-NG2 Cre mice had a greater percentage of cells of the OL lineage (Olig2^+^) than control littermates at P10 and P15 and similar numbers at P30 (**Fig. 3B**). The percentage of cells that were mature OLs (CC1^+^) was higher in p38γ^f/f^-NG2 Cre mice than in controls at P15 and no difference was observed at P30 (**Fig. 3C**), while the percentage of cells that were OPCs (PDGFrα^+^) was lower in p38γ^f/f^-NG2 Cre at P10 (**Fig. 3D**). Thus, early ablation of p38γ from OPCs results in a transient increase in cells of the OL lineage that can be attributed to earlier differentiation of OPCs into OLs. However, changes in the rate of proliferation could also influence the observed effects. We labelled proliferating cells with EdU (injected at P6) and found that the fraction of Olig2^+^ cells proliferating was higher in p38γ^f/f^-NG2 Cre mice than in the control littermates (**Supplementary Fig. 3A, B**), but it was lower in cells labelled *in vitro* (**Supplementary Fig. 3C**). To analyse the effect of ablation on OPC differentiation independently from the effect on proliferation, Tx was administered from P7 to P11 (rather than from P2 to P6) (**Fig. 3E**); recombination occurred in most mature CC1^+^ OLs at P15 (determined by tdTomato reporter expression in double transgenics expressing NG2 Cre) (**Fig. 3E**). When p38γ was ablated from this population of OLs (by administering Tx to p38γ^f/f^-NG2 Cre mice at P7–11), the percentage of Olig2^+^ cells at P15 was the same as that in the littermate controls (**Fig. 3F**) but the proportions of mature OLs (MBP^+^, **Fig. 3G**; CC1^+^ **Fig. 3H**) were higher. Thus, the enhanced differentiation was not the result of greater proliferation. These *in vivo* experiments indicate that p38γ ablation accelerates OPC differentiation in a cell-autonomous fashion, resulting in the early onset of myelination.

**Figure 3.**
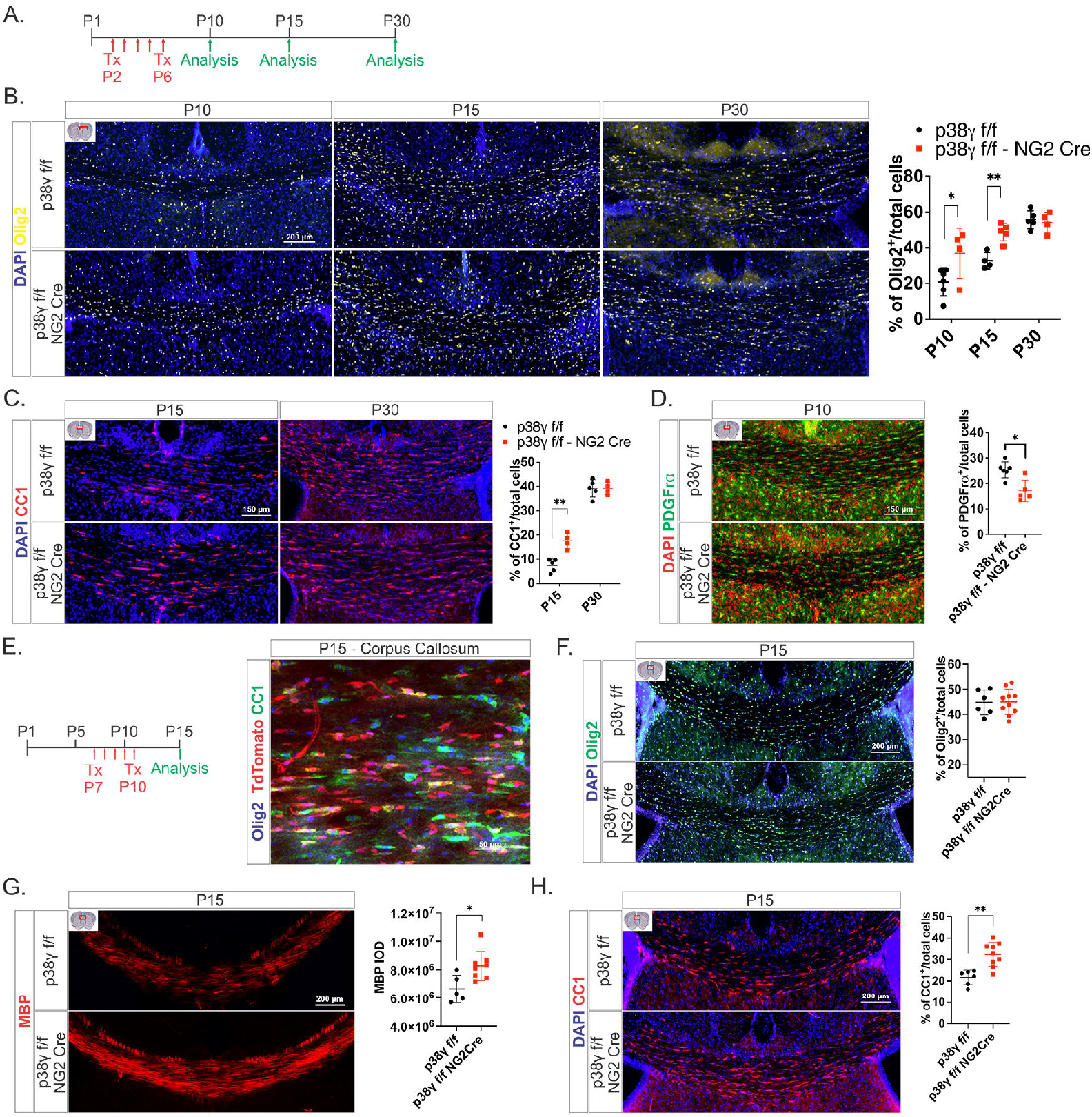
p38γ^f/f^-NG2 Cre mice show accelerated OPC differentiation. (**A**) Schematic representation of Tx injections and analysis time points. (**B**) Immunohistochemical analysis of coronal sections from mice at P10, P15, and P30 showing higher percentages of Olig2^+^ OLs in the corpus callosa of p38γ^f/f^-NG2 Cre mice than in those of p38γ^f/f^ controls at P10 and P15 but not at P30. (**C**) Immunostaining for CC1 in coronal sections of mice at P15 and P30 showing a higher percentage of mature OLs in the corpus callosa of p38γ^f/f^-NG2 Cre mice than in those of controls at P15 but not P30. (**D**) p38γ^f/f^-NG2 Cre mice had a lower percentage of PDGFrα^+^ OPCs than controls at P10. (**E**) Schematic representation of delayed Tx injections for p38γ ablation after a period of rapid proliferation; this paradigm results in Cre-mediated recombination in most Olig2^+^ OLs in the corpus callosum at P15. Immunostaining in coronal sections containing corpus callosa of mice at P15 showing that the percentages of Olig2^+^ cells in p38γ^f/f^-NG2 Cre mice were similar to those in control mice at P15 (**F**) but the percentages of MBP^+^ (**G**) and CC1^+^ (**H**) OLs were higher when Tx administration was delayed. Values are expressed as the means ± SEMs. \**P* < 0.05 and \*\**P* < 0.01 by Student’s *t* test.

### p38γ ablation accelerates OPC migration *ex vivo* and results in ERK1/2 activation

To determine whether p38γ ablation affects OPC migration, we tracked the migration of tdTomato-expressing OPCs in acute brain slices from P3 mice (p38γ^f/f^ [or p38γ^f/+^]-tdTomato-NG2 Cre mice). Coronal slices were cultured for 2 days in the presence of 1 μM Tx to induce p38γ ablation and tdTomato expression before PDGF-AA and basic fibroblast growth factor were added to promote migration. Time-lapse images were collected over 24 h to measure OPC speed and distance travelled. We found that OPCs in slices from p38γ^f/f^-NG2 Cre mice migrated faster and further than those from controls (**Fig. 4A, B** and **Supplementary Videos**).

**Figure 4.**
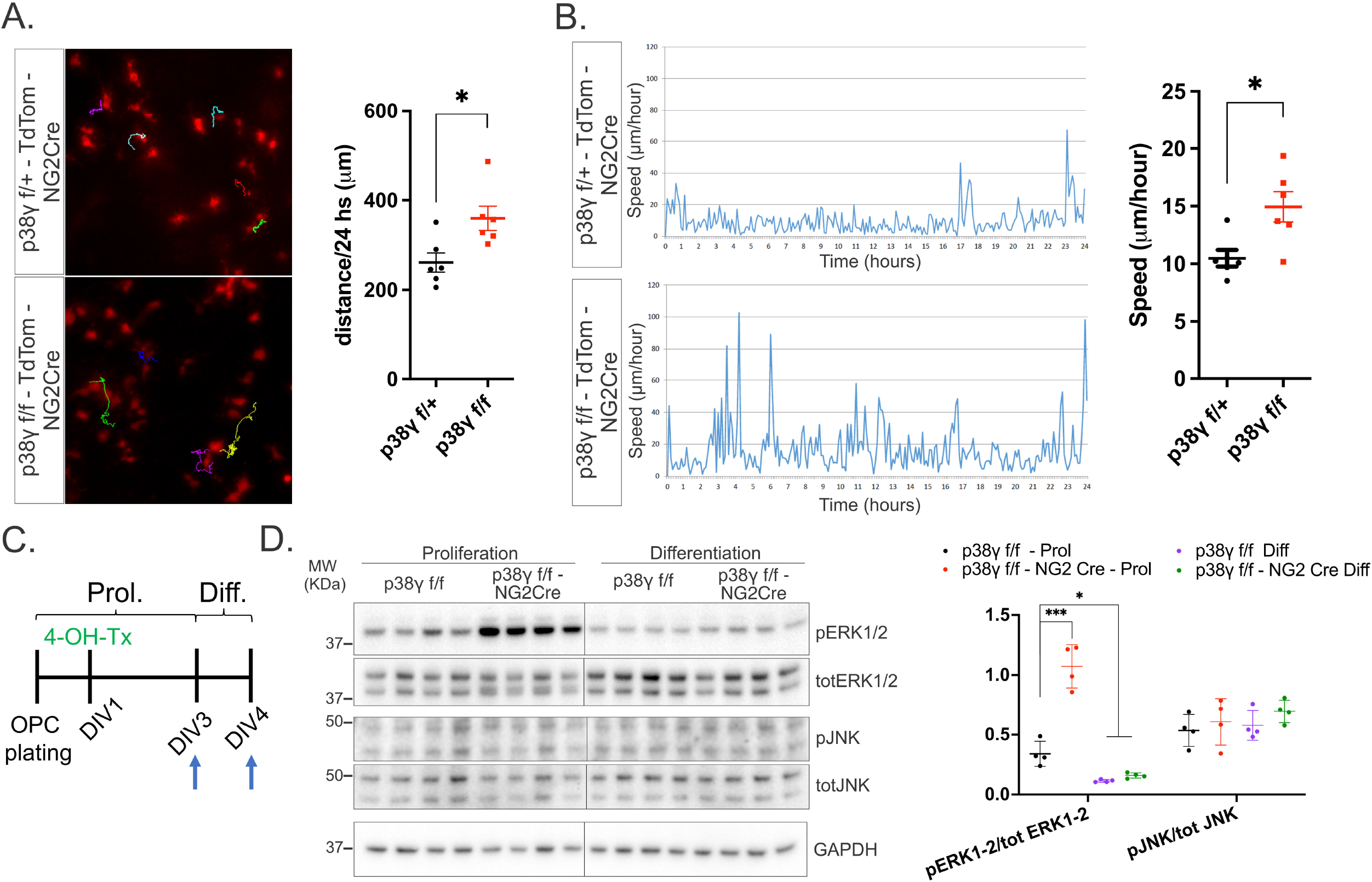
p38γ ablation accelerates OPC migration *ex vivo* and results in activation of the ERK1/2 pathway *in vitro*. OPC migration was tracked in acute coronal slice cultures from P3 tdTomato-NG2 Cre mice homozygous or heterozygous for the floxed p38γ allele. After 48 h in culture, slices were treated with Tx for 24 h to induce p38γ ablation and/or tdTomato expression. After PDGF-AA (10 ng/ml) and basic fibroblast growth factor (10 ng/ml) were added to the medium, time-lapse images were taken over 24 h to measure the speed and distance travelled of migrating OPCs. OPCs from p38γ^f/f^ mice travelled greater distances (**A**) and moved at higher speeds (**B**) than those from p38γ^f/+^ controls. (**C**) Schematic of OPC culture procedure for data in panel D. Cell lysates were obtained under conditions of proliferation (DIV3) and after 24 h of differentiation (DIV4). (**D**) Western blot analysis for total and phosphorylated ERK1/2 and JNK in OPC lysates at DIV3 and DIV4 showing that p38γ^f/f^-NG2 Cre OPCs have increased activation of ERK1/2 under conditions of proliferation and that JNK activation is not affected. Values are expressed as the means ± SEMs. *\*P* < 0.05 and \*\*\**P* < 0.001 by Student’s *t* test or one-way ANOVA followed by Newman–Keuls multiple-comparisons post-test.

Because multiple MAPKs are known to regulate the migration of different cell types, we sought to identify which of these might contribute to the migration phenotype in OPCs with p38γ ablation. Specifically, we assessed the activation of ERK1/2 and JNK in OPC cultures under conditions of proliferation (DIV3) and differentiation (DIV4) (**Fig. 4C**). Western blots of cell lysates showed increased phosphorylation of ERK/1/2 (relative to total ERK1/2 levels) in p38γ-ablated OPCs under conditions of proliferation, indicating increased activation, whereas no change in the phosphorylation of JNK was observed (**Fig. 4D**). Thus, the effect of p38γ ablation on migration may be mediated by ERK1/2 signalling.

### p38γ regulates OPC differentiation *in vitro*

To study the mechanisms by which p38γ influences OPC differentiation, we performed *in vitro* studies using OPCs isolated from p38γ^f/f^-NG2 Cre mice and p38γ^f/f^ controls. After plating, Tx was added to the culture medium for p38γ ablation, which was confirmed at DIV3 by Western blotting (**Fig. 5A**). After 48 h under differentiation conditions (DIV5), immunocytochemistry analyses showed that greater proportions of p38γ^f/f^-NG2 Cre OPCs were positive for MBP and PLP and a smaller proportion expressed PDGFrα (**Fig. 5B**), providing further evidence of the cell-autonomous effect of p38γ ablation on accelerated OPC differentiation. The increases in MBP and PLP in cells from p38γ^f/f^-NG2 Cre mice were confirmed by Western blotting (**Fig. 5C**).

**Figure 5.**
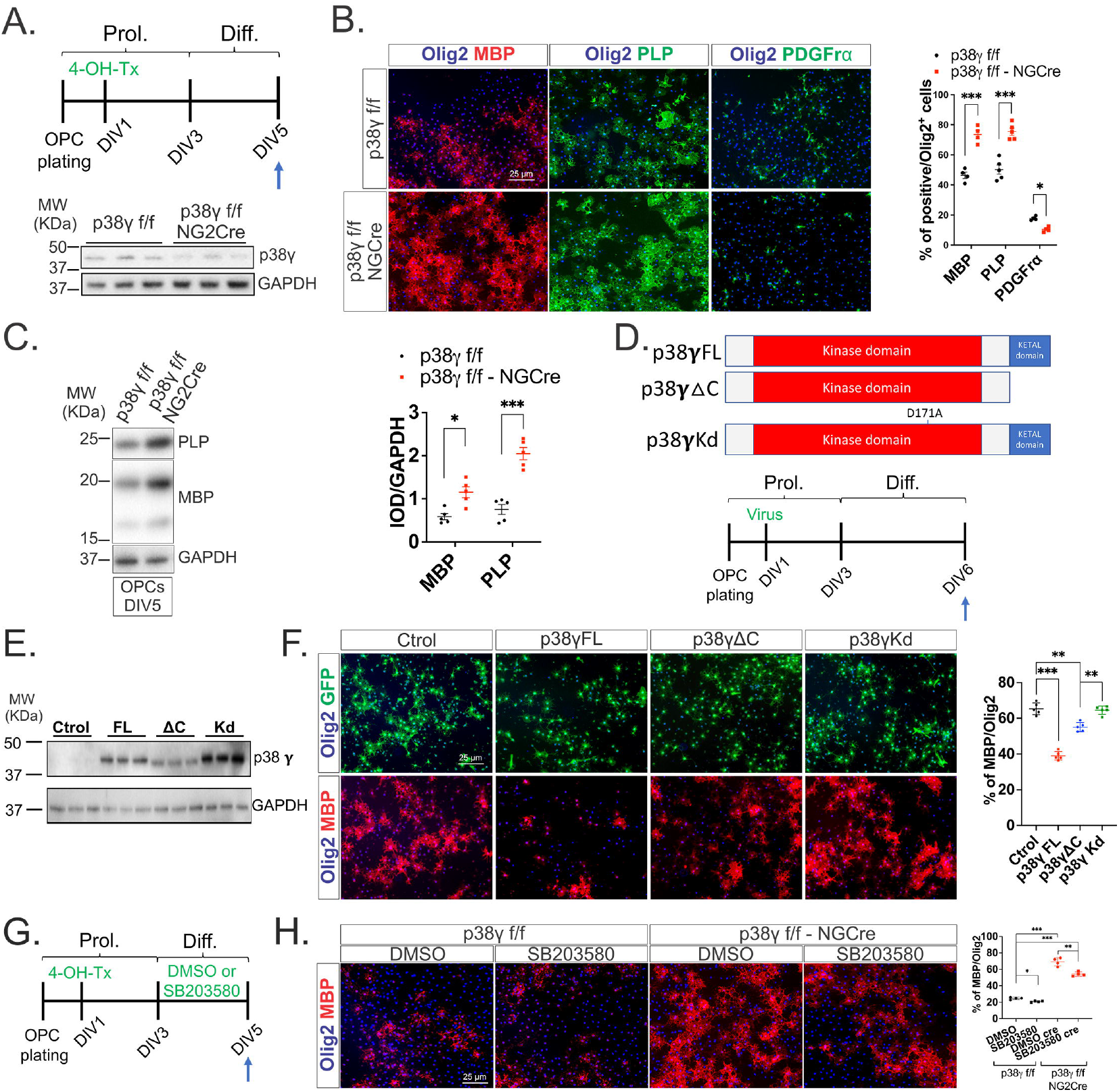
p38γ^f/f^-NG2 Cre OPCs differentiate earlier *in vitro* and p38α inhibition antagonizes this effect. (**A**) Schematic representation of OPC culture procedures for panels B and C. Tx was added to the culture medium for 48 h to induce p38γ ablation, which was confirmed by Western blotting of OPC lysates at DIV3. (**B**) Immunohistochemical analysis of OPC cultures after 48 h of differentiation (DIV5) showing that greater proportions of Olig2^+^ OPCs express PLP and MBP and a smaller proportion expresses PDGFrα when p38γ is ablated. (**C**) Western blot analysis of OPC lysates at DIV5 showing that p38γ^f/f^-NG2 Cre OPCs have increased amounts of MBP and PLP. (**D**) Schematic representations of the different p38γ constructs for overexpression of the full-length WT p38γ (p38γFL), the KETAL-del mutant lacking the C-terminal binding domain (p38γΔC), and the kinase-dead mutant (p38γKd). OPCs were infected with viruses harbouring these constructs at DIV1, indicated in the schematic at the bottom for data in panels E and F. (**E**) Western blots of OPC lysates at DIV3 confirming the overexpression of the different p38γ constructs. (**F**) Immunocytochemical analysis of OPCs after 72 h of differentiation (DIV6) showing that p38γFL overexpression drastically reduces the percentage Olig2^+^ cells that express MBP, whereas the effect was not as strong for p38γ missing the C-terminal domain and absent with the kinase-dead mutant. (**G**) Schematic representation of the treatment of OPC cultures with the p38α inhibitor SB203580 or dimethyl sulfoxide (DMSO) as a control for data in panel H. (**H**) Immunocytochemical analysis of OPCs after 48 h of differentiation showing that p38α inhibition decreases the percentage of Olig2^+^ cells that express MBP. Values are expressed as the means ± SEMs. \**P* < 0.05, \*\**P* < 0.01, and \*\*\**P* < 0.001 by Student’s *t* test or one-way ANOVA followed by Newman–Keuls multiple-comparisons posttest.

Using this *in vitro* system, we next explored if overexpression of p38γ can delay OPC differentiation. OPCs isolated from WT mice were infected with lentiviruses harbouring constructs for the full-length p38γ (p38γFL), p38γ missing the C-terminal KETAL domain that mediates interactions with the PDZ domains of substrate proteins (p38ΔC), a kinase-dead mutant p38γ (p38γKd), or green fluorescent protein (GFP) as a control (**Fig. 5D**). Overexpression of p38γ and the protein size was confirmed by Western blotting (**Fig. 5E**). Overexpression of p38γFL reduced the proportion of Olig2^+^ cells that expressed MBP (**Fig. 5F**). Overexpression of the truncated p38γ also reduced OPC differentiation, but to a lesser extent, which suggests that the effect of the full-length protein was partially attributable to PDZ-dependent binding of substrates. Notably, overexpression of the inactive mutant did not impact differentiation (**Fig. 5F**), indicating that the enzymatic activity of p38γ is required.

### p38α contributes to the effect of p38γ ablation on OPC differentiation

p38α accelerates OPC differentiation and developmental myelination.^26,31,32,55^ To determine if the effect of p38γ ablation on OPC differentiation is independent of p38α activity, we treated OPCs from p38γ^f/f^-NG2 Cre mice or p38γ^f/f^ controls with an inhibitor of p38α/β (SB203580) during differentiation (**Fig. 5G**). Our findings showed that p38α/β inhibition limited the differentiation of control and p38γ^f/f^-NG2 Cre OPCs (**Fig. 5H**), indicating that the effect of p38γ ablation may be partially dependent on p38α/β.

### p38γ influences the OPC transcriptome

To study the effects of p38γ on OPC gene expression we performed RNA-sequencing of WT and p38γ^-/-^ OPCs from P7 mice. Our initial analysis identified 490 genes that were dysregulated in p38γ^-/-^ OPCs. Of these, we focused on 82 genes that were shown to be enriched in OPCs according to previously published RNA-sequencing data^56^ for pathway analysis (**Fig. 6A**). In line with our previous findings, we found that several of the genes identified in the GOBERT_OLIGODENDROCYTE_DIFFERENTIATION_UP pathway were upregulated in p38γ^-/-^ OPCs (false discovery rate, 5.49 × 10^-29^) (**Fig. 6B**). Despite our contrasting results *in vivo* and *in vitro*, both GO and KEGG pathway enrichment analyses showed that cell cycle-related genes were upregulated in p38γ^-/-^ OPCs (false discovery rate, 6.77 × 10^-33^) (**Fig. 6C, D**). These data indicate that p38γ influences the size of the population of OPCs that mature into OLs.

**Figure 6.**
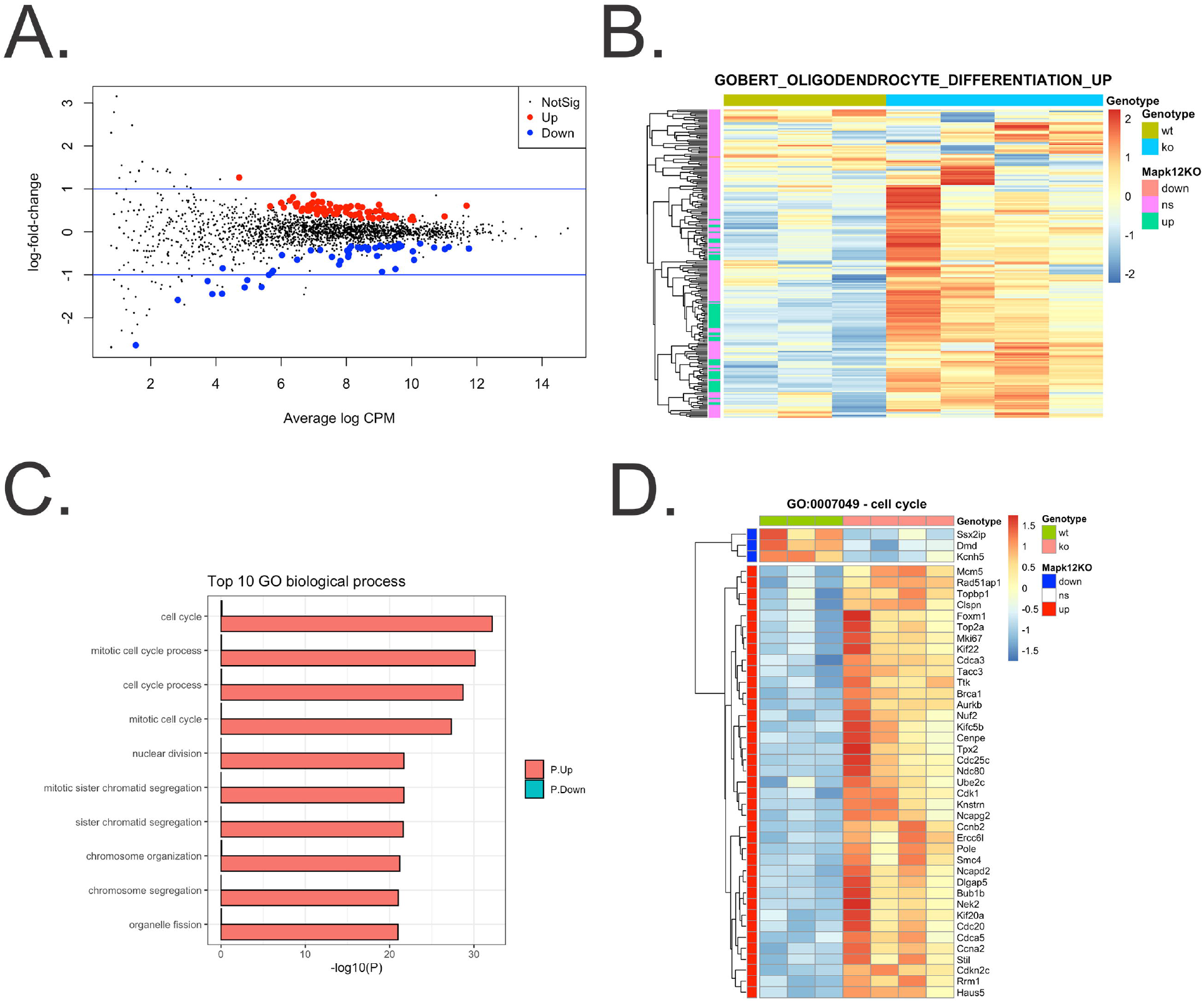
RNA-sequencing of p38γ^-/-^ OPCs. (**A**) Log_2_ fold changes of genes expressed in p38γ^-/-^ versus WT OPCs plotted against the average count size for every gene (CPM, counts per million). Red points represent genes that were significantly upregulated and blue points represent genes significantly downregulated at a <5% false discovery rate. (**B**) Heat map of the GOBERT_OLIGODENDROCYTE_DIFFERENTIATION_UP pathway^56^ showing the upregulated genes in p38γ^-/-^ OPCs in green. (**C**) GO analysis of biological processes significantly upregulated in p38γ^-/-^ OPCs. (**D**) Heat map of the GO cell cycle category showing downregulated genes in p38γ null OPCs in blue and upregulated genes in red.

### p38γ ablation accelerates OPC differentiation and remyelination after injury

To determine if p38γ ablation accelerates OPC differentiation and remyelination after injury, we used the cuprizone (CPZ) model of demyelination-remyelination, in which 7 weeks of cuprizone treatment (CPZ 7) results in demyelination of mouse brain and 2 weeks of recovery (CPZ 7+2) allows for remyelination.^38,39^ We found that the expression of p38γ is strongly upregulated upon demyelination (CPZ 7) in cortex, corpus callosum, and caudato putamen and that these high levels persist in the cortex and caudato putamen into the remyelination phase (CPZ 7+2) (**Supplementary Fig. 4A**). ISH for p38γ mRNA followed by immunohistochemical analysis revealed that p38γ was strongly upregulated in PDGFrα^+^ OPCs and Iba1^+^ microglia, but not in GFAP^+^ astrocytes during CPZ insult (**Supplementary Fig. 4B**).

To study the impact of p38γ ablation from OPCs during remyelination, we administered Tx to p38γ^f/f^-NG2 mice and control littermates starting at the last 5 days of the 7-week CPZ treatment and continued the Tx injections for 5 days into the 2-week recovery (**Fig. 7A**). CPZ induced similar levels of demyelination in mice with and without p38γ ablation as observed with immunohistochemistry for MBP in the corpus collosum, striatum (including caudato putamen), and cortex (**Fig. 7B**). However, after remyelination (CPZ 7+2), the levels of MBP were much higher in p38γ^f/f^-NG2 Cre animals (**Fig. 7B**), which was confirmed by Western blotting (**Fig. 7C**). The percentages of cells in the corpus collosum that were Olig2^+^ and CC1^+^ mature OLs during demyelination were similar in mice with or without p38γ ablation, but were higher in those with p38γ ablation after 2 weeks of recovery (CPZ 7+2) (**Fig. 7D, 7E**). Moreover, electron microscopy analyses revealed that these increases resulted in greater levels of myelination in the corpus callosa of p38γ^f/f^-NG2 mice at CPZ 7+2. Notably, this was observed not only as a greater density of myelinated axons (**Fig. 7F, G**), but also as thicker myelin sheaths (i.e., lower g-ratio) (**Fig. 7H**). Scatter plots showing g-ratio versus axon diameter with log-linear fits^50^ indicate that the smaller remyelinated axons had the thicker myelin in p38γ^f/f^-NG2 Cre mice (**Fig. 7H**). These *in vivo* experiments indicate that p38γ ablation accelerates OPC differentiation in the adult brain after injury, resulting in accelerated and more efficient remyelination.

**Figure 7.**
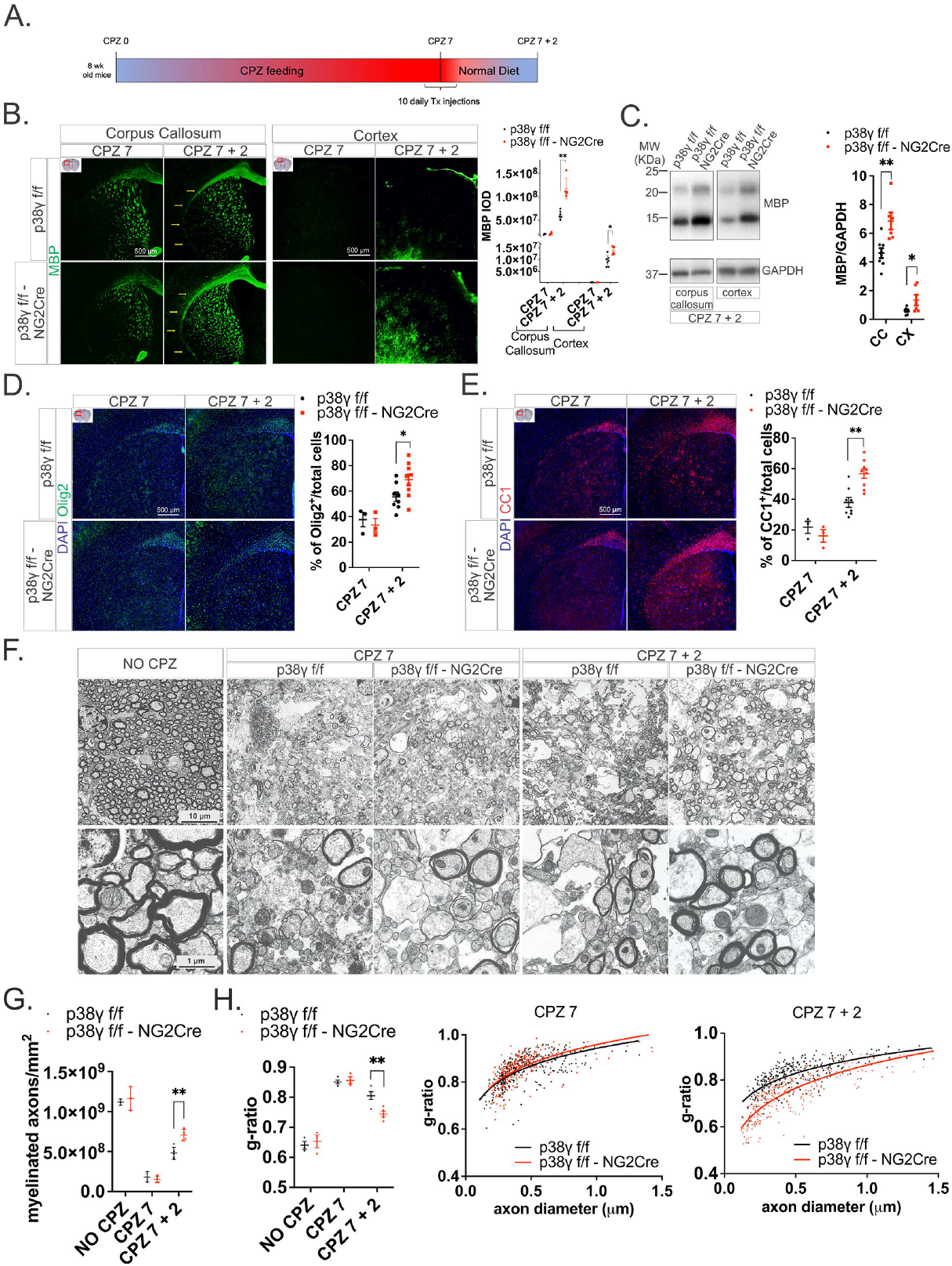
p38γ^f/f^-NG2 Cre mice show accelerated remyelination. (**A**) Schematic representation of CPZ feeding for demyelination and remyelination and Tx injections for p38γ ablation from OPCs. (**B**) Immunohistochemical analysis of coronal sections showing that p38γ^f/f^-NG2 Cre and control (p38γ^f/f^) mice have similar MBP expression in the corpus callosum and cortex after demyelination (CPZ 7) but p38γ^f/f^-NG2 Cre mice have greater expression after the 2-week remyelination/recovery period (CPZ 7+2). (**C**) Western blot analysis confirming that p38γ^f/f^-NG2 Cre mice have greater amounts of MBP in the corpus callosum and cortex at CPZ 7+2. Immunohistochemical analyses of coronal sections showing that p38γ^f/f^-NG2 Cre mice have higher percentages of Olig2^+^ OLs (**D**) and CC1^+^ mature OLs (**E**) in the corpus callosum than control mice at CPZ 7+2. (**F**) Electron microscopy analysis of cross sections of the axons that comprise the corpus callosum under normal conditions (left) and in p38γ^f/f^-NG2 Cre and control mice after demyelination (CPZ 7) (middle) and remyelination (CPZ 7+2) (right). (**G**) Quantifications of the density of myelinated axons in the corpus callosum. (**H**) Quantifications of myelin thickness (g-ratio) and g-ratio versus axon diameter scatter plots after demyelination (CPZ 7) and remyelination (CPZ 7 + 2). Values are expressed as the means ± SEMs. \**P* < 0.05 and \*\**P* < 0.01 by Student’s *t* tests. Statistical analysis of data in scatter plots was by log-linear fits with a 95% confidence interval, and slopes were compared with the extra sum of squares *F* test. CPZ 7: p38γ^f/f^, slope = 0.1186, *R*^2^ = 0.4756; absolute sum of squares = 0.6870; p38γ^f/f^-NG2 Cre, slope = 0.1259, *R*^2^ = 0.4439, absolute sum of squares = 0.5675 (*P* = 0.5067). CPZ 7 + 2: p38γ^f/f^, slope = 0.1155, *R*^2^ = 0.5393, absolute sum of squares = 0.5748; p38γ^f/f^-NG2 Cre, slope = 0.1751, *R*^2^ = 0.6759, absolute sum of squares = 0.8016 (*P* < 0.0001).

### p38γ is expressed by OPCs in white matter and cortical multiple sclerosis lesions

The data shown so far suggest that p38γ slows (re)myelination. To investigate this in the context of Multiple Sclerosis, in which remyelination becomes less efficient in later stages of the disease, we examined microarray data generated in a previous report by our collaborators, Dr. Dutta and Dr. Trapp.^46^ Those data showed that p38γ mRNA is upregulated in white matter lesions (WML) compared to levels in normal appearing white matter (NAWM) in multiple sclerosis tissue or in healthy white matter from a control brain (**Fig. 8A**). This increase was also apparent when comparing p38γ expression in NAWM and WML from the same individual (**Fig. 8B**). We performed ISH for p38γ followed by MBP staining in previously characterized white matter and cortical lesions.^46^ Our findings show that p38γ was strongly expressed in the cortex and WML where MBP content was low or absent compared to p38γ expression in NAWM (**Fig. 8C and 8D**). Detailed analysis of cortical areas that were demyelinated and shadow plaques (**Fig. 8E,** insets 1 and 2, enlarged in **8E**), confirmed our findings that p38γ expression and myelin content are inversely associated. Additional immunohistochemical analyses revealed that the cell types expressing p38γ (identified by ISH) included PDGFrα^+^ OPCs in the WML border facing NAWM and in the WML core (similar to region I) (**Fig. 8F**, red arrows). Few cells in these regions immunostained positive for MHCII (for antigen-presenting cells) or Iba1 (for microglia), suggesting that this lesion was immune inactive (**Fig. 8F**, right). These data support the hypothesis that p38γ is an inhibitor of remyelination in multiple sclerosis, which could thus be exploited as a potential therapeutic target to promote remyelination.

**Figure 8.**
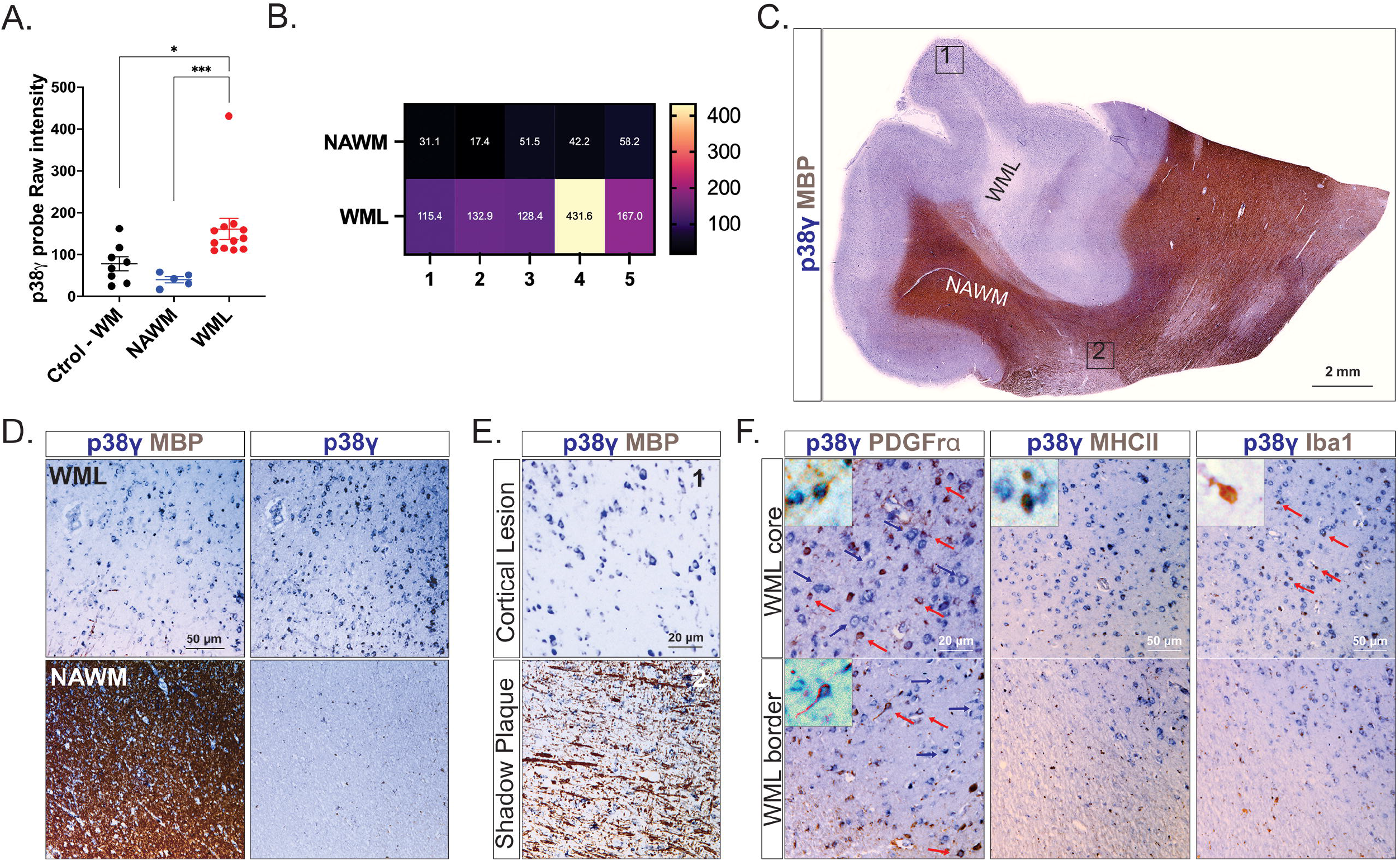
p38γ is expressed by OPCs in Multiple Sclerosis lesions. (**A**) Results from a microarray analysis^46^ of Multiple Sclerosis lesions showing p38γ expression in healthy control white matter (Ctrol-WM) and in NAWM and WML from patients with Multiple Sclerosis. (**B**) Heat map of p38γ expression in NAWM and WML from each of five individuals. Absolute values are shown inside each box. (**C** and **E**) ISH for p38γ followed by MBP immunostaining showing that p38γ is strongly expressed in WML when compared to NAWM (**D**), cortical lesion and shadow plaques (E). (**F**) ISH for p38γ followed by immunohistochemistry for PDGFrα for OPCs, MHCII for antigen-presenting cells, or Iba1 for microglia in the leukocortical lesion (area 1 in panel **C**). Both the border and core of the lesion show OPCs lesion express p38γ (red arrows) and other cell types that express p38γ (blue arrows). Few cells were positive for MHCII or Iba1. Values are expressed as the means ± SEMs. \**P* < 0.05 and \*\*\**P* < 0.001 by one-way ANOVA followed by Newman–Keuls multiple-comparisons post-test.

## Discussion

Here we describe the minor γ isoform of p38 MAPK as a novel inhibitor of myelination in the CNS during development and after injury. As OPC differentiation and myelination progress, the levels of p38γ decrease. This was observed both *in vitro* and *in vivo*. By conditionally ablating p38γ from OPCs in mice (p38-NG2 Cre mice), we showed that the decrease in p38γ accelerates OPC differentiation, developmental myelination, and remyelination after injury. We further showed that the deletion of p38γ from OPCs increases the speed and distance at which they migrate. Proliferating OPCs from p38^f/f^-NG2 Cre mice had greater activation of ERK1/2, which is known to influence OPC migration during development.^57,58^ Further studies will determine if there is cross talk between p38 and ERK1/2 pathways under these conditions.

Our data indicate that p38γ delays OPC differentiation and myelination during development and after myelin injury. The finding that p38γ deletion increased not only the number of myelinated fibers, but also their thickness is of particular importance, because remyelination is almost never efficient and usually results in the formation of a thinner myelin sheath (reviewed by Franklin and Ffrench-Constant^9^). The effect of p38’ on the thickness of repaired myelin is, to our knowledge, a unique property not reported for other myelin inhibitors. p38γ was barely detectable in the brains of adult mice but was strongly upregulated during CPZ-induced demyelination, consistent with previous data showing that CPZ treatment activates the p38 pathway in corpus callosa lysates.^59^ We also found that as the animals recovered and remyelination progressed, p38γ levels quickly downregulated. Although the CPZ model lacks the autoimmune nature of multiple sclerosis, it elicits microgliosis, astrogliosis, and the production of inflammatory molecules^38,60,61 62^ and is a useful tool for exploring remyelination potential.

With the progression of the disease state in individuals with multiple sclerosis, the lesion environment becomes restrictive to OPC differentiation and subsequent remyelination.^8,9^ We found that p38γ expression was much higher in WMLs than in NAWM in patient samples. Our ISH findings confirm that PDGFrα^+^ OPCs within these lesions expressed p38γ, suggesting that p38γ is an inhibitor of OPC differentiation in Multiple Sclerosis. Furthermore, the p38γ expression was in WMLs largely devoid of microglia and antigen-presenting cells. Multiple Sclerosis lesions are classified according to their inflammatory and demyelinating activity^63,64^: lesions with a large number of microglia/macrophages/antigen-presenting cells are considered active lesions and have the potential to remyelinate, whereas lesions with low numbers of these cells are inactive and fail to remyelinate. Our findings support the idea that p38γ is an inhibitor of remyelination in the context of Multiple Sclerosis.

Inhibition of p38γ is a more promising therapy to promote myelin repair than p38α/β inhibition, because it impacts both developmental myelination and remyelination and will likely have no or limited side effects. This is because the expression of p38γ (and the other minor isoform, p38δ) is more restricted than that of p38α/β and knockout mice have no obvious phenotype.^54^ Several preclinically validated p38α inhibitors underwent clinical trials for various inflammatory conditions, but only one candidate (pirfenidone, a weak inhibitor) was approved for the treatment of idiopathic pulmonary fibrosis (reviewed by Burton^65^; ClinicalTrials.gov number N022535).

Currently, no approved treatment exists to repair existing lesions in patients with multiple sclerosis, though many approaches have been investigated (reviewed by Bove^66^ and Hauser^67^) and some have undergone clinical trials. Among these are clemastine (a histamine/muscarinic receptor antagonist), GSK239512 (a histamine H3 receptor antagonist), and opicinumab (a LINGO-1 blocking antibody). Clemastine promotes OPC differentiation^12^ and remyelination in CPZ-^68^ and EAE-induced demyelination^13^ and was thus investigated in patients with relapsing remitting multiple sclerosis under immunomodulatory therapy (NCT02040298;ReBUILD). Although there was improvement in visual evoked potentials, no significant effects on other clinical or MRI outcome measures were observed.^69^ GSK239512 was tested as a potential remyelinating agent because activation of the H3 receptor inhibits OPC differentiation.^70^ Similarly to clemastine, GSK239512 was tested in patients with relapsing remitting multiple sclerosis (NCT01772199) and a small positive effect on remyelination was observed by MRI. Inhibition of LINGO-1 promotes OPC differentiation and myelination^71^ and remyelination after lysolecithin/CPZ- or EAE-induced demyelination^14^ with no immunomodulatory activity, but trials in patients with unilateral acute optic neuritis (NCT01721161; RENEW)^74^ and relapsing remitting multiple sclerosis (NCT01864148; SYNERGY)^75^ showed no real clinical benefit. Immunomodulatory therapy, such as interferon-beta treatment, negatively impacts OPC differentiation^76^ and remyelination,^77^ thereby restricting the window of opportunity for remyelinating therapies. Alternatively, a novel target with immunomodulatory activity, which is also capable of promoting OPC differentiation and remyelination could be greatly beneficial.

Recent work showed that the p38γ/δ subfamily mediates aspects of immune system activation, including cytokine production in macrophages and dendritic cells,^78^ neutrophil migration and recruitment to inflammation sites,^79^ and alveolar mucus production.^80^ Furthermore, p38γ and p38δ have been shown to mediate autoimmunity.^81^ This, combined with the roles demonstrated herein on OPC differentiation and remyelination indicate that p38γ inhibition, possibly coupled with p38δ inhibition, represents a potentially game-changing therapeutic target for multiple sclerosis.

## Abbreviations

Akt: protein kinase B
bFGF: basic fibroblast growth factor
CNTF: ciliary neurotrophic factor
ERK1/2: extracellular signal-regulated kinase 1/2
FBS: foetal bovine serum
GFP: green fluorescent protein
ISH: *in situ* hybridization
JNK: c-Jun N-terminal kinase
MAG: myelin-associated glycoprotein
MAPK: mitogen-activated protein kinase
MBP: myelin basic protein
NT3: Neurotrophin 3
OL: oligodendrocyte
OPC: oligodendrocyte precursor cell
PDGFrα: platelet-derived growth factor receptor alpha
PFA: paraformaldehyde
PLP: proteolipid protein
RT: room temperature
T3: 3,3, 5-Triiodo-L-thyronine
Tx: tamoxifen
4-OH-Tx: 4 hydroxy-Tx
WT: wild type

## Acknowledgements

We thank Pablo Paez for sharing equipment and expertise for the migration experiments, Karen Dietz for critical reading of the manuscript, and Bianca Weisnstock Guttman and members of our laboratory for constructive discussions.

## Supplementary material

Supplementary material is available at *Brain* online.

## Supplementary Material

**Supplementary Fig 1.**
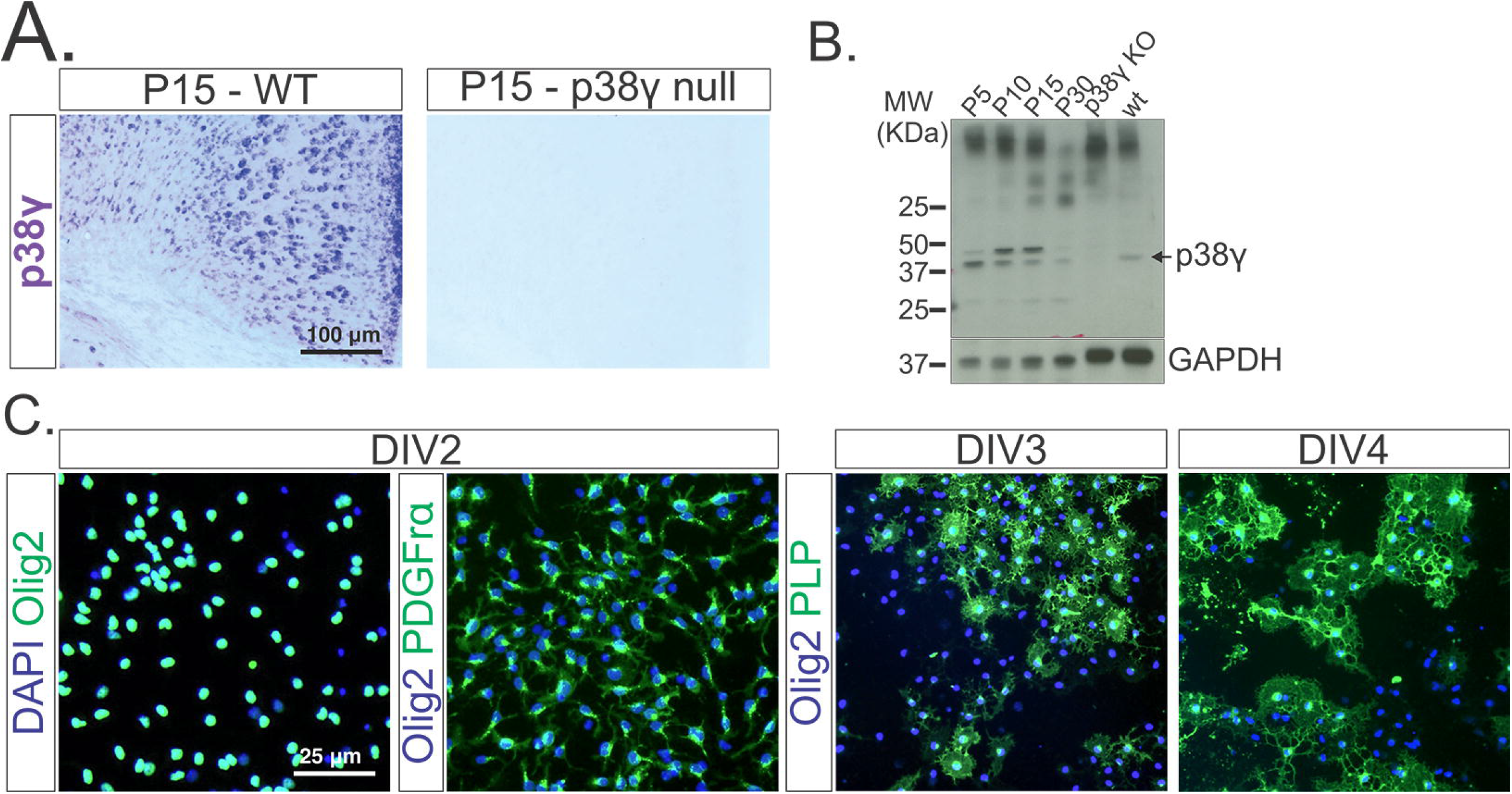
(**A**) Validation of the p38γ antisense probe showing lack of signal in p38γ^-/-^ (null) coronal brain sections. (**B**) Validation of the p38γ antibody showing lack of signal in p38γ^-/-^ (KO) whole brain lysates. (**C**) OPC cultures were of high purity, shown by Olig2 immunohistochemistry. At the end of proliferation (DIV2), most OLs were PDGFrα^+^, and after 24 h (DIV3) or 48 h (DIV4) of differentiation, the number of PLP^+^ mature OLs greatly increased.

**Supplementary Fig 2.**
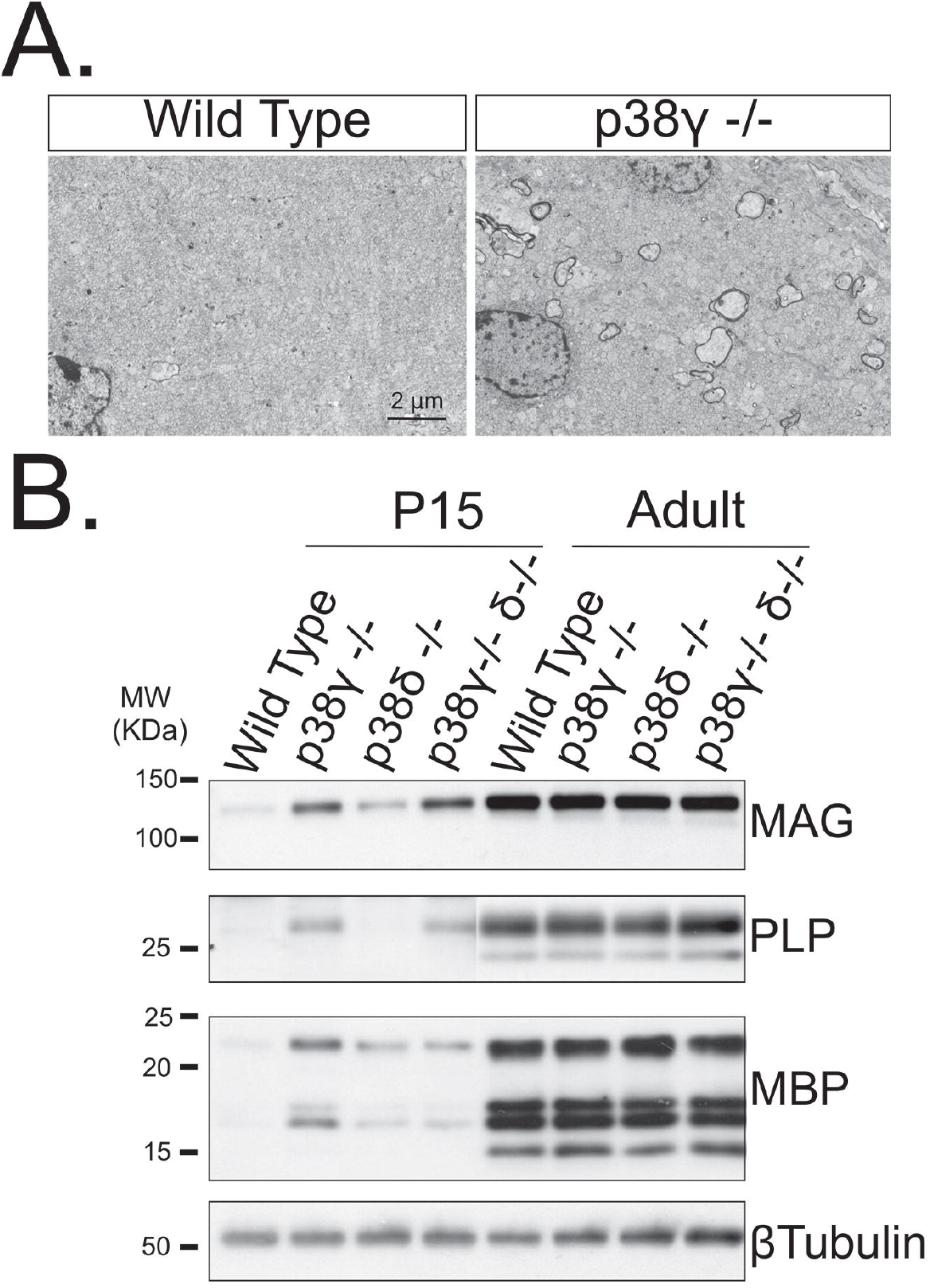
p38γ^-/-^ null mice show accelerated myelination and p38δ^-/-^ mice have no phenotype. (**A**) Electron microscopy of P15 corpus callosum showing more myelinated axons in the p38γ^-/-^ mouse. (**B**) Western blot analysis of corpus callosum lysates from mice at P15 and 3 months (adult) showing that p38γ^-/-^ mice have increased amounts of myelin proteins MAG, MBP, and PLP) and that p38δ^-/-^ mice have amounts similar to those in the controls.

**Supplementary Fig 3.**
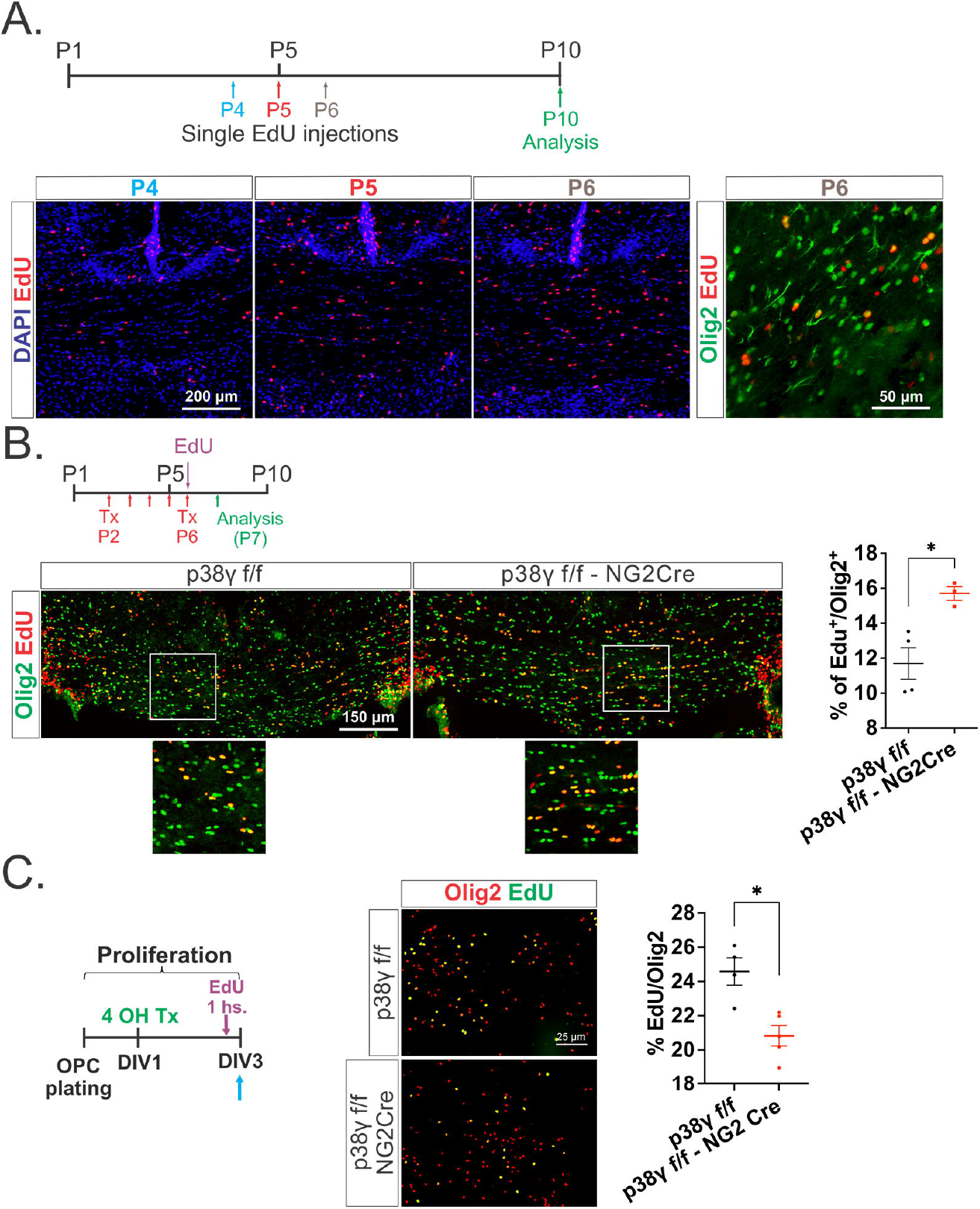
p38γ^-/-^ OPCs show increased proliferation *in vivo* and decreased proliferation *in vitro*. (**A**) (Top) Schematic representation of EdU injection in WT mice. A single intraperitoneal injection of EdU was given to WT mice at P4, P5, or P6, and brains were collected at P10. (Bottom) Sections analysed at P10 show the increase in the number of proliferating cells between P4 and P6. A proportion of Olig2^+^ cells incorporated EdU injected at P6. (**B**) (Top) Schematic representation of Tx and EdU injections. (Bottom) Immunohistochemical analysis of coronal brain sections of p38γ^f/f^-NG2 Cre and p38γ^f/f^ mice showing increased percentage of Edu^+^ Olig2^+^ OLs in the corpus callosa of the p38γ^f/f^-NG2 Cre mice at P7. (**C**) (Left) Schematic representation of OPC culture procedures. EdU was added during proliferation 1 h before the cells were fixed. (Right) Immunohistochemical analysis showing reduced percentage of Edu^+^ Olig2^+^ OPCs in p38γ^f/f^-NG2 Cre cultures. Values are expressed as the means ± SEMs. \**P* < 0.05 by Student’s *t* test.

**Supplementary Fig 4.**
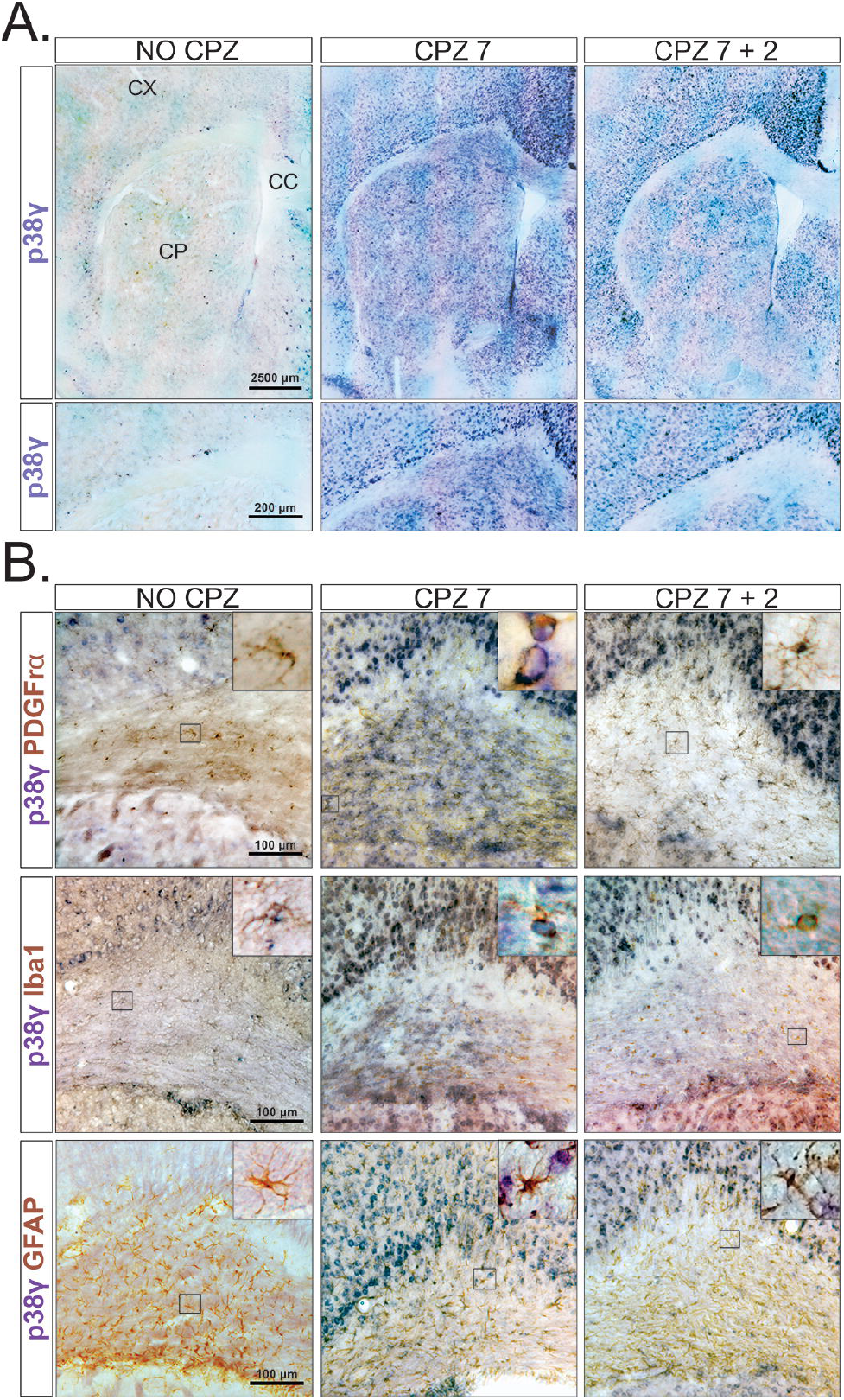
p38γ is upregulated during CPZ-induced demyelination in WT animals. (**A**) ISH for p38γ in coronal brain sections from animals fed a normal diet (NO CPZ), CPZ for 7 weeks (CPZ 7; demyelination), or CPZ for 7 weeks followed by a normal diet for 2 weeks (CPZ 7+2; remyelination). Results show that p38γ is ubiquitously upregulated after CPZ-induced demyelination (CPZ 7). p38γ upregulation persists in the cortex (CX) and the caudate putamen (CP) and decreases in corpus callosum (CC) after remyelination (CPZ 7+2). (**B**) ISH for p38γ followed by immunohistochemistry of coronal sections for PDGFrα (OPCs), Iba1 (microglia), and GFAP (astrocytes). p38γ is strongly upregulated in OPCs and microglia after demyelination (CPZ 7) and undetectable in astrocytes.

**Supplementary Fig 5.**
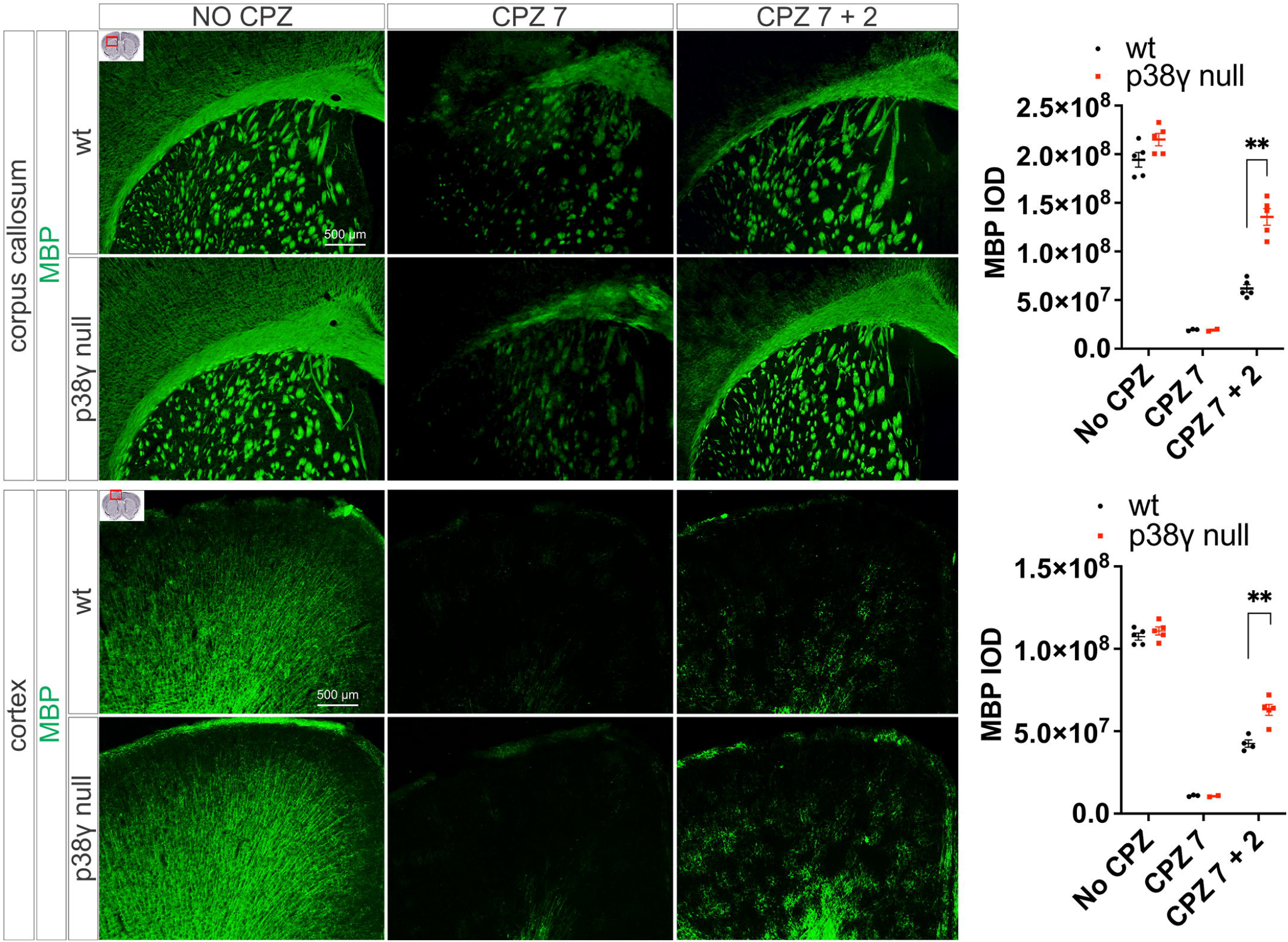
p38γ null mice show accelerated remyelination. Immunohistochemical analysis of coronal brain sections showing that p38γ^-/-^ (null) and control (WT) mice have similar expression of MBP in the corpus callosum and cortex after demyelination (CPZ 7). After remyelination (CPZ 7+2), p38γ^-/-^ mice show higher amounts of MBP than control mice in the cortex and corpus callosum. \*\**P* < 0.01.

