## Supplementary material for "p38γMAPK delays myelination and remyelination and is abundant in multiple sclerosis lesions": Sup Table 1

**Supp. Table 1. Primers**

| Gene | Forward | Forwar/Reverse | Reverse | Annealing Temperature (°C) |
|---|---|---|---|---|
| MAPK12 null | CCTGAGGTTTAGATAGGCTGTATGTCTCACTCACAC | CACTTCGCCCAATAGCAGCCAGTCCCTTCC | GAATTCCCAGTAGGTCATTCCTGGGACCATCC | 65 |
| MAPK12 flox | CCAGGAGGTGACCAAAACGGC | TGGGCTGCGAAGGTAGAGGTG | CTGAGGCGGAAAGAACCAGCT | 65 |
| MAPK12 recombination | TGGGCTGCGAAGGTAGAGGTG | | GTGTCACGTGCTCAGGGCCTG | 55 |
| Cspg4 (NG2Cre) | CAATCCTAACCAGGAGCAAGAG | GCGCGCCTGAAGATATAGAA | CAAGTCGCAAGTCGCAATTC | 61 |
| MAPK12 | CCATACTTTGAGTCCCTTCGG | - | AGTCACACGCTTCCATTCC | 60 |
| HPRT | CCTCATGGACTGATTATGGACAG | - | TCAGCAAAGAACTTATAGCCCC | 60 |
