## Supplementary material for "p38γMAPK delays myelination and remyelination and is abundant in multiple sclerosis lesions": Sup Table 2

**Sup. Table 2. Antibodies**

| Antibody | Source | Identifier (RRID) |
|---|---|---|
| β Tubulin | Sigma-Aldrich T4026 | AB_477577 |
| CC1 | EMD Millipore OP80 | AB_2057371 |
| Phopho ERK1/2 | Cell Signaling Technologies 9101 | AB_331646 |
| Total ERK1/2 | Cell Signaling Technologies 9102 | AB_330744 |
| GFAP | Sigma-Aldrich G9269 | AB_477035 |
| GAPDH | Sigma-Aldrich G9545 | AB_796208 |
| Iba1 | Wako 019-19741 | AB_839504 |
| Phosphor JNK | Cell Signaling Technology 9255 | AB_2307321 |
| Total JNK | Cell Signaling Technology 9252 | AB_2250373 |
| MAG | EMD Millipore MAB1567 | AB_2137847 |
| MBP | Covance SMI99P | AB_10120129 |
| MHCII | BD Biosciences 555557 | AB_395939 |
| NeuN | EMD Millipore MAB377 | AB_2298772 |
| Olig2 | Proteintech 13999-1-AP | AB_2157541 |
| P38γ | Proteintech 20184-1-AP | AB_10665949 |
| PDGFrα | Neuromics gt15150 | AB_2737233 |
| PDGFrα - Panning | BD Pharmingen 558774 | AB_397117 |
| PLP | Generous gift of Dr. Paez | AB_2341144 |
