## Supplementary material for "p38γMAPK delays myelination and remyelination and is abundant in multiple sclerosis lesions": Sup Table 3

| | NGenes | Direc | PValue | FDR |
|---|---|---|---|---|
| ROSTY_CERVICAL_CANCER_PROLIFERATION_CLUSTER | 85 | Up | 1.71E-38 | 8.12E-35 |
| SOTIRIOU_BREAST_CANCER_GRADE_1_VS_3_UP | 74 | Up | 2.39E-37 | 5.67E-34 |
| FISCHER_DREAM_TARGETS | 255 | Up | 1.15E-32 | 1.75E-29 |
| KOBAYASHI_EGFR_SIGNALING_24HR_DN | 111 | Up | 1.48E-32 | 1.75E-29 |
| MARSON_BOUND_BY_E2F4_UNSTIMULATED | 173 | Up | 5.98E-32 | 5.49E-29 |
| GOBERT_OLIGODENDROCYTE_DIFFERENTIATION_UP | 217 | Up | 6.95E-32 | 5.49E-29 |
| FLORIO_NEOCORTEX_BASAL_RADIAL_GLIA_DN | 107 | Up | 1.27E-30 | 8.60E-28 |
| FISCHER_G2_M_CELL_CYCLE | 94 | Up | 3.71E-29 | 2.20E-26 |
| HOFFMANN_LARGE_TO_SMALL_PRE_BII_LYMPHOCYTE_UP | 66 | Up | 5.96E-29 | 3.14E-26 |
| LEE_EARLY_T_LYMPHOCYTE_UP | 52 | Up | 5.02E-27 | 2.38E-24 |
| HORIUCHI_WTAP_TARGETS_DN | 94 | Up | 1.01E-26 | 4.37E-24 |
| DUTERTRE_ESTRADIOL_RESPONSE_24HR_UP | 148 | Up | 1.57E-26 | 6.20E-24 |
| KONG_E2F3_TARGETS | 56 | Up | 2.97E-26 | 1.08E-23 |
| MORI_IMMATURE_B_LYMPHOCYTE_DN | 44 | Up | 1.60E-25 | 5.43E-23 |
| GAVIN_FOXP3_TARGETS_CLUSTER_P6 | 44 | Up | 5.78E-25 | 1.78E-22 |
| BENPORATH_CYCLING_GENES | 168 | Up | 6.13E-25 | 1.78E-22 |
| REACTOME_CELL_CYCLE_MITOTIC | 122 | Up | 6.38E-25 | 1.78E-22 |
| CROONQUIST_NRAS_SIGNALING_DN | 40 | Up | 1.01E-24 | 2.65E-22 |
| CROONQUIST_IL6_DEPRIVATION_DN | 57 | Up | 1.64E-24 | 4.09E-22 |
| CHIANG_LIVER_CANCER_SUBCLASS_PROLIFERATION_UP | 76 | Up | 2.51E-24 | 5.94E-22 |
| REACTOME_CELL_CYCLE | 148 | Up | 2.69E-24 | 6.07E-22 |
| SHEDDEN_LUNG_CANCER_POOR_SURVIVAL_A6 | 139 | Up | 4.68E-24 | 1.01E-21 |
| MORI_LARGE_PRE_BII_LYMPHOCYTE_UP | 37 | Up | 5.73E-24 | 1.18E-21 |
| VECCHI_GASTRIC_CANCER_EARLY_UP | 108 | Up | 7.45E-24 | 1.46E-21 |
| TANG_SENESCENCE_TP53_TARGETS_DN | 32 | Up | 7.68E-24 | 1.46E-21 |
| GRAHAM_CML_DIVIDING_VS_NORMAL_QUIESCENT_UP | 72 | Up | 1.15E-22 | 2.10E-20 |
| ZHOU_CELL_CYCLE_GENES_IN_IR_RESPONSE_24HR | 70 | Up | 1.31E-22 | 2.29E-20 |
| GOLDRATH_ANTIGEN_RESPONSE | 93 | Up | 1.92E-22 | 3.25E-20 |
| GRAHAM_NORMAL_QUIESCENT_VS_NORMAL_DIVIDING_DN | 45 | Up | 4.39E-22 | 7.18E-20 |
| MARKEY_RB1_ACUTE_LOF_UP | 69 | Up | 1.02E-21 | 1.60E-19 |
| BASAKI_YBX1_TARGETS_UP | 88 | Up | 1.55E-21 | 2.37E-19 |
| CAIRO_HEPATOBLASTOMA_CLASSES_UP | 118 | Up | 1.68E-21 | 2.49E-19 |
| BURTON_ADIPOGENESIS_3 | 53 | Up | 2.32E-21 | 3.33E-19 |
| ODONNELL_TFRC_TARGETS_DN | 59 | Up | 3.68E-21 | 5.13E-19 |
| ZHAN_MULTIPLE_MYELOMA_PR_UP | 32 | Up | 3.92E-21 | 5.31E-19 |
| BLUM_RESPONSE_TO_SALIRASIB_DN | 104 | Up | 7.69E-21 | 1.01E-18 |
| WHITEFORD_PEDIATRIC_CANCER_MARKERS | 55 | Up | 1.69E-20 | 2.17E-18 |
| WU_APOPTOSIS_BY_CDKN1A_VIA_TP53 | 31 | Up | 2.58E-20 | 3.22E-18 |
| REACTOME_CELL_CYCLE_CHECKPOINTS | 71 | Up | 6.09E-20 | 7.39E-18 |
| ISHIDA_E2F_TARGETS | 29 | Up | 8.74E-20 | 1.04E-17 |
| PUJANA_BRCA2_PCC_NETWORK | 119 | Up | 1.27E-19 | 1.47E-17 |
| WONG_EMBRYONIC_STEM_CELL_CORE | 84 | Up | 1.73E-19 | 1.96E-17 |
| BENPORATH_PROLIFERATION | 52 | Up | 2.01E-19 | 2.21E-17 |
| MORI_PRE_BI_LYMPHOCYTE_UP | 33 | Up | 2.91E-19 | 3.14E-17 |

| | | | | |
|---|---|---|---|---|
| KANG_DOXORUBICIN_RESISTANCE_UP | 38 | Up | 3.05E-19 | 3.21E-17 |
| ZHOU_CELL_CYCLE_GENES_IN_IR_RESPONSE_6HR | 49 | Up | 3.42E-19 | 3.52E-17 |
| PUJANA_CHEK2_PCC_NETWORK | 128 | Up | 4.23E-19 | 4.26E-17 |
| WHITFIELD_CELL_CYCLE_G2 | 53 | Up | 8.31E-19 | 8.20E-17 |
| REACTOME_M_PHASE | 72 | Up | 1.41E-18 | 1.36E-16 |
| BURTON_ADIPOGENESIS_PEAK_AT_24HR | 22 | Up | 1.96E-18 | 1.86E-16 |
| REACTOME_MITOTIC_PROMETAPHASE | 56 | Up | 4.05E-18 | 3.76E-16 |
| WHITFIELD_CELL_CYCLE_LITERATURE | 26 | Up | 4.82E-18 | 4.39E-16 |
| FURUKAWA_DUSP6_TARGETS_PCI35_DN | 34 | Up | 6.25E-18 | 5.59E-16 |
| FERREIRA_EWINGS_SARCOMA_UNSTABLE_VS_STABLE_UP | 49 | Up | 7.62E-18 | 6.69E-16 |
| BERENJENO_TRANSFORMED_BY_RHOA_UP | 127 | Up | 8.19E-18 | 7.05E-16 |
| WINNEPENNINCKX_MELANOMA_METASTASIS_UP | 64 | Up | 1.39E-17 | 1.18E-15 |
| REACTOME_RESOLUTION_OF_SISTER_CHROMATID_COHESION | 36 | Up | 2.54E-17 | 2.11E-15 |
| SARRIO_EPITHELIAL_MESENCHYMAL_TRANSITION_UP | 71 | Up | 2.78E-17 | 2.27E-15 |
| KAMMINGA_EZH2_TARGETS | 25 | Up | 4.15E-17 | 3.33E-15 |
| PUJANA_XPRSS_INT_NETWORK | 62 | Up | 5.20E-17 | 4.11E-15 |
| VILLANUEVA_LIVER_CANCER_KRT19_UP | 54 | Up | 7.82E-17 | 6.07E-15 |
| CHEMNITZ_RESPONSE_TO_PROSTAGLANDIN_E2_UP | 41 | Up | 1.00E-16 | 7.65E-15 |
| REACTOME_MITOTIC_METAPHASE_AND_ANAPHASE | 45 | Up | 1.27E-16 | 9.55E-15 |
| NUYTTEN_EZH2_TARGETS_DN | 239 | Up | 1.35E-16 | 9.96E-15 |
| LE_EGR2_TARGETS_UP | 46 | Up | 1.64E-16 | 1.20E-14 |
| SHEPARD_CRUSH_AND_BURN_MUTANT_DN | 67 | Up | 3.37E-16 | 2.42E-14 |
| NAKAYAMA_SOFT_TISSUE_TUMORS_PCA2_UP | 38 | Up | 3.83E-16 | 2.71E-14 |
| WHITFIELD_CELL_CYCLE_G2_M | 61 | Up | 5.53E-16 | 3.86E-14 |
| LINDGREN_BLADDER_CANCER_CLUSTER_3_UP | 91 | Up | 7.07E-16 | 4.85E-14 |
| EGUCHI_CELL_CYCLE_RB1_TARGETS | 18 | Up | 7.84E-16 | 5.31E-14 |
| LI_WILMS_TUMOR_VS_FETAL_KIDNEY_1_DN | 50 | Up | 1.32E-15 | 8.71E-14 |
| MISSIAGLIA_REGULATED_BY_METHYLATION_DN | 50 | Up | 1.32E-15 | 8.71E-14 |
| GREENBAUM_E2A_TARGETS_UP | 17 | Up | 1.48E-15 | 9.60E-14 |
| WANG_RESPONSE_TO_GSK3_INHIBITOR_SB216763_DN | 88 | Up | 3.04E-15 | 1.94E-13 |
| FARMER_BREAST_CANCER_CLUSTER_2 | 23 | Up | 3.06E-15 | 1.94E-13 |
| ZHENG_GLIOBLASTOMA_PLASTICITY_UP | 101 | Up | 6.93E-15 | 4.32E-13 |
| YU_MYC_TARGETS_UP | 22 | Up | 7.37E-15 | 4.54E-13 |
| PID_PLK1_PATHWAY | 22 | Up | 8.95E-15 | 5.43E-13 |
| FEVR_CTNNB1_TARGETS_DN | 123 | Up | 9.13E-15 | 5.47E-13 |
| RHODES_UNDIFFERENTIATED_CANCER | 26 | Up | 2.05E-14 | 1.22E-12 |
| CASORELLI_ACUTE_PROMYELOCYTIC_LEUKEMIA_DN | 111 | Up | 2.18E-14 | 1.28E-12 |
| REACTOME_SEPARATION_OF_SISTER_CHROMATIDS | 37 | Up | 3.41E-14 | 1.97E-12 |
| BIDUS_METASTASIS_UP | 44 | Up | 5.54E-14 | 3.16E-12 |
| KINSEY_TARGETS_OF_EWSR1_FLII_FUSION_UP | 265 | Up | 6.79E-14 | 3.83E-12 |
| PUJANA_BRCA1_PCC_NETWORK | 190 | Up | 8.35E-14 | 4.65E-12 |
| SHEPARD_BMYB_TARGETS | 38 | Up | 9.15E-14 | 5.04E-12 |
| TOYOTA_TARGETS_OF_MIR34B_AND_MIR34C | 94 | Up | 9.83E-14 | 5.35E-12 |
| PUJANA_BREAST_CANCER_WITH_BRCA1_MUTATED_UP | 29 | Up | 1.11E-13 | 5.96E-12 |
| REACTOME_MITOTIC_SPINDLE_CHECKPOINT | 33 | Up | 1.42E-13 | 7.56E-12 |

| | | | | |
|---|---|---|---|---|
| PUJANA_BRCA_CENTERED_NETWORK | 48 | Up | 2.89E-13 | 1.52E-11 |
| FRASOR_RESPONSE_TO_SERM_OR_FULVESTRANT_DN | 22 | Up | 3.66E-13 | 1.91E-11 |
| MITSIADES_RESPONSE_TO_APLIDIN_DN | 63 | Up | 4.53E-13 | 2.33E-11 |
| TARTE_PLASMA_CELL_VS_PLASMABLAST_DN | 60 | Up | 7.05E-13 | 3.59E-11 |
| REACTOME_RHO_GTPASES_ACTIVATE_FORMINS | 31 | Up | 9.44E-13 | 4.76E-11 |
| PETROVA_ENDOTHELIUM_LYMPHATIC_VS_BLOOD_UP | 38 | Up | 3.79E-12 | 1.89E-10 |
| REICHERT_MITOSIS_LIN9_TARGETS | 20 | Up | 4.70E-12 | 2.32E-10 |
| GARGALOVIC_RESPONSE_TO_OXIDIZED_PHOSPHOLIPIDS_TURQUOISE_DN | 19 | Up | 7.16E-12 | 3.50E-10 |
| LINDGREN_BLADDER_CANCER_CLUSTER_1_DN | 90 | Up | 8.04E-12 | 3.89E-10 |
| PID_AURORA_B_PATHWAY | 17 | Up | 9.57E-12 | 4.58E-10 |
| CROONQUIST_NRAS_VS_STROMAL_STIMULATION_DN | 34 | Up | 1.41E-11 | 6.70E-10 |
| ODONNELL_TARGETS_OF_MYC_AND_TFRC_DN | 28 | Up | 1.48E-11 | 6.93E-10 |
| REACTOME_MITOTIC_G2_G2_M_PHASES | 34 | Up | 1.88E-11 | 8.73E-10 |
| PID_AURORA_A_PATHWAY | 9 | Up | 2.04E-11 | 9.41E-10 |
| GAL_LEUKEMIC_STEM_CELL_DN | 51 | Up | 2.48E-11 | 1.13E-09 |
| DODD_NASOPHARYNGEAL_CARCINOMA_DN | 262 | Up | 2.79E-11 | 1.26E-09 |
| ALCALAY_AML_BY_NPM1_LOCALIZATION_DN | 55 | Up | 2.83E-11 | 1.26E-09 |
| MANALO_HYPOXIA_DN | 68 | Up | 3.26E-11 | 1.44E-09 |
| FUJII_YBX1_TARGETS_DN | 76 | Up | 3.35E-11 | 1.47E-09 |
| MOLENAAR_TARGETS_OF_CCND1_AND_CDK4_DN | 29 | Up | 7.48E-11 | 3.25E-09 |
| DELPUECH_FOXO3_TARGETS_DN | 10 | Up | 1.02E-10 | 4.41E-09 |
| REACTOME_G2_M_CHECKPOINTS | 36 | Up | 1.14E-10 | 4.87E-09 |
| POOLA_INVASIVE_BREAST_CANCER_UP | 55 | Up | 1.17E-10 | 4.94E-09 |
| AFFAR_YY1_TARGETS_DN | 88 | Up | 1.20E-10 | 5.05E-09 |
| RIZ_ERYTHROID_DIFFERENTIATION | 18 | Up | 1.51E-10 | 6.29E-09 |
| YANG_BCL3_TARGETS_UP | 88 | Up | 2.08E-10 | 8.56E-09 |
| PETROVA_PROX1_TARGETS_UP | 11 | Up | 2.10E-10 | 8.56E-09 |
| REACTOME_KINESINS | 15 | Up | 2.12E-10 | 8.59E-09 |
| RUIZ_TNC_TARGETS_DN | 61 | Up | 2.17E-10 | 8.71E-09 |
| FINETTI_BREAST_CANCER_KINOME_RED | 14 | Up | 2.42E-10 | 9.65E-09 |
| SCIAN_CELL_CYCLE_TARGETS_OF_TP53_AND_TP73_DN | 13 | Up | 2.66E-10 | 1.05E-08 |
| MUELLER_PLURINET | 54 | Up | 2.69E-10 | 1.05E-08 |
| REACTOME_CYCLIN_A_B1_B2_ASSOCIATED_EVENTS_DURING_G2_M_TRANSITION | 10 | Up | 3.12E-10 | 1.21E-08 |
| NADERI_BREAST_CANCER_PROGNOSIS_UP | 19 | Up | 3.95E-10 | 1.52E-08 |
| REN_BOUND_BY_E2F | 26 | Up | 6.35E-10 | 2.43E-08 |
| REACTOME_MITOTIC_G1_PHASE_AND_G1_S_TRANSITION | 37 | Up | 9.83E-10 | 3.73E-08 |
| KEGG_CELL_CYCLE | 39 | Up | 9.94E-10 | 3.74E-08 |
| BOYAULT_LIVER_CANCER_SUBCLASS_G3_UP | 30 | Up | 1.15E-09 | 4.28E-08 |
| REACTOME_APC_C_MEDIATED_DEGRADATION_OF_CELL_CYCLE_PROTEINS | 17 | Up | 1.16E-09 | 4.28E-08 |
| KAUFFMANN_MELANOMA_RELAPSE_UP | 30 | Up | 1.26E-09 | 4.61E-08 |
| SMID_BREAST_CANCER_LUMINAL_A_DN | 10 | Up | 1.29E-09 | 4.71E-08 |
| SONG_TARGETS_OF_IE86_CMV_PROTEIN | 20 | Up | 1.38E-09 | 5.00E-08 |

| | | | | |
|---|---|---|---|---|
| PID_FOXM1_PATHWAY | 13 | Up | 1.49E-09 | 5.34E-08 |
| PATIL_LIVER_CANCER | 126 | Up | 1.62E-09 | 5.77E-08 |
| WEST_ADRENOCORTICAL_TUMOR_UP | 60 | Up | 2.31E-09 | 8.16E-08 |
| LY_AGING_MIDDLE_DN | 10 | Up | 2.55E-09 | 8.96E-08 |
| MORI_MATURE_B_LYMPHOCYTE_DN | 17 | Up | 2.75E-09 | 9.59E-08 |
| BENPORATH_ES_1 | 84 | Up | 2.78E-09 | 9.62E-08 |
| FOURNIER_ACINAR_DEVELOPMENT_LATE_2 | 69 | Up | 3.83E-09 | 1.31E-07 |
| JOHNSTONE_PARVB_TARGETS_3_DN | 185 | Up | 4.18E-09 | 1.42E-07 |
| REACTOME_CONDENSATION_OF_PROMETAPHASE_CHROMOSOMES | 7 | Up | 5.01E-09 | 1.69E-07 |
| MONTERO_THYROID_CANCER_POOR_SURVIVAL_UP | 9 | Up | 5.18E-09 | 1.74E-07 |
| KAUFFMANN_DNA_REPAIR_GENES | 54 | Up | 5.32E-09 | 1.77E-07 |
| REACTOME_DNA_REPLICATION | 32 | Up | 5.45E-09 | 1.81E-07 |
| SHEPARD_BMYB_MORPHOLINO_DN | 72 | Up | 6.00E-09 | 1.98E-07 |
| REACTOME_RHO_GTPASE_EFFECTORS | 48 | Up | 6.83E-09 | 2.23E-07 |
| SMIRNOV_RESPONSE_TO_IR_6HR_DN | 26 | Up | 7.99E-09 | 2.58E-07 |
| REACTOME_COPI_DEPENDENT_GOLGI_TO_ER_RETROGRADE_TRAFFIC | 18 | Up | 8.01E-09 | 2.58E-07 |
| VANTVEER_BREAST_CANCER_METASTASIS_DN | 36 | Up | 9.48E-09 | 3.04E-07 |
| BOYAULT_LIVER_CANCER_SUBCLASS_G23_UP | 17 | Up | 1.01E-08 | 3.20E-07 |
| KAUFFMANN_DNA_REPLICATION_GENES | 36 | Up | 1.16E-08 | 3.66E-07 |
| SU_TESTIS | 18 | Up | 1.29E-08 | 4.06E-07 |
| REACTOME_GOLGI_TO_ER_RETROGRADE_TRANSPORT | 19 | Up | 1.55E-08 | 4.83E-07 |
| BOHN_PRIMARY_IMMUNODEFICIENCY_SYNDROM_UP | 12 | Up | 1.84E-08 | 5.68E-07 |
| PID_ATR_PATHWAY | 14 | Up | 2.14E-08 | 6.59E-07 |
| LIANG_SILENCED_BY_METHYLATION_DN | 6 | Up | 2.32E-08 | 7.10E-07 |
| STEIN_ESRRA_TARGETS_RESPONSIVE_TO_ESTROGEN_DN | 19 | Up | 2.64E-08 | 8.02E-07 |
| REACTOME_S_PHASE | 36 | Up | 3.11E-08 | 9.39E-07 |
| LY_AGING_OLD_DN | 20 | Up | 3.25E-08 | 9.74E-07 |
| LI_WILMS_TUMOR_ANAPLASTIC_UP | 9 | Up | 3.27E-08 | 9.76E-07 |
| GRAHAM_CML_QUIESCENT_VS_NORMAL_QUIESCENT_UP | 26 | Up | 4.56E-08 | 1.35E-06 |
| HU_GENOTOXIC_DAMAGE_4HR | 13 | Up | 5.17E-08 | 1.52E-06 |
| PUJANA_ATM_PCC_NETWORK | 129 | Up | 5.95E-08 | 1.74E-06 |
| XU_HGF_TARGETS_INDUCED_BY_AKT1_48HR_DN | 8 | Up | 6.47E-08 | 1.88E-06 |
| KIM_WT1_TARGETS_DN | 93 | Up | 7.76E-08 | 2.24E-06 |
| PAL_PRMT5_TARGETS_UP | 44 | Up | 8.79E-08 | 2.52E-06 |
| SCIBETTA_KDM5B_TARGETS_DN | 29 | Up | 8.91E-08 | 2.54E-06 |
| SASAKI_ADULT_T_CELL_LEUKEMIA | 32 | Up | 1.02E-07 | 2.89E-06 |
| REACTOME_DNA_REPAIR | 56 | Up | 1.12E-07 | 3.16E-06 |
| TIEN_INTESTINE_PROBIOTICS_24HR_UP | 102 | Up | 1.15E-07 | 3.23E-06 |
| REACTOME_REGULATION_OF_TP53_ACTIVITY | 24 | Up | 1.20E-07 | 3.35E-06 |
| RODRIGUES_THYROID_CARCINOMA_POORLY_DIFFERENTIATED_UP | 145 | Up | 1.30E-07 | 3.60E-06 |
| REACTOME_SIGNALING_BY_RHO_GTPASES | 70 | Up | 1.35E-07 | 3.72E-06 |
| KEGG_PROGESTERONE_MEDIATED_OOCYTE_MATURATION | 12 | Up | 1.39E-07 | 3.80E-06 |

| | | | | |
|---|---|---|---|---|
| LE_NEURONAL_DIFFERENTIATION_DN | 7 | Up | 1.56E-07 | 4.24E-06 |
| COLINA_TARGETS_OF_4EBP1_AND_4EBP2 | 87 | Up | 1.64E-07 | 4.45E-06 |
| NAKAMURA_CANCER_MICROENVIRONMENT_DN | 19 | Up | 1.74E-07 | 4.68E-06 |
| WILCOX_RESPONSE_TO_PROGESTERONE_UP | 41 | Up | 1.90E-07 | 5.07E-06 |
| REACTOME_POLO_LIKE_KINASE_MEDIATED_EVENTS | 8 | Up | 2.26E-07 | 6.02E-06 |
| FERRANDO_T_ALL_WITH_MLL_ENL_FUSION_DN | 16 | Up | 3.22E-07 | 8.52E-06 |
| REACTOME_MITOTIC_PROPHASE | 14 | Up | 3.50E-07 | 9.22E-06 |
| REACTOME_HOMOLOGY_DIRECTED_REPAIR | 33 | Up | 4.36E-07 | 1.14E-05 |
| VANTVEER_BREAST_CANCER_ESR1_DN | 60 | Up | 4.46E-07 | 1.16E-05 |
| MORI_EMU_MYC_LYMPHOMA_BY_ONSET_TIME_UP | 25 | Up | 4.92E-07 | 1.27E-05 |
| KEGG_DNA_REPLICATION | 15 | Up | 5.02E-07 | 1.29E-05 |
| WHITFIELD_CELL_CYCLE_S | 41 | Up | 5.68E-07 | 1.45E-05 |
| REACTOME_TRANSCRIPTIONAL_REGULATION_BY_TP53 | 44 | Up | 5.83E-07 | 1.48E-05 |
| REACTOME_NEURONAL_SYSTEM | 65 | Down | 5.98E-07 | 1.52E-05 |
| ZERBINI_RESPONSE_TO_SULINDAC_DN | 4 | Up | 6.74E-07 | 1.70E-05 |
| RODRIGUES_THYROID_CARCINOMA_ANAPLASTIC_UP | 124 | Up | 7.22E-07 | 1.81E-05 |
| REACTOME_CONDENSATION_OF_PROPHASE_CHROMOSOMES | 7 | Up | 8.00E-07 | 2.00E-05 |
| PID_E2F_PATHWAY | 17 | Up | 8.36E-07 | 2.07E-05 |
| REACTOME_DNA_STRAND_ELONGATION | 16 | Up | 8.42E-07 | 2.08E-05 |
| PUJANA_BREAST_CANCER_LIT_INT_NETWORK | 27 | Up | 8.68E-07 | 2.13E-05 |
| KEGG_OOCYTE_MEIOSIS | 17 | Up | 9.90E-07 | 2.42E-05 |
| WEST_ADRENOCORTICAL_TUMOR_MARKERS_UP | 13 | Up | 1.27E-06 | 3.08E-05 |
| REACTOME_MHC_CLASS_II_ANTIGEN_PRESENTATION | 14 | Up | 1.48E-06 | 3.59E-05 |
| REACTOME_REGULATION_OF_PLK1_ACTIVITY_AT_G2_M_TRANSITION | 22 | Up | 1.53E-06 | 3.67E-05 |
| REACTOME_DNA_DOUBLE_STRAND_BREAK_REPAIR | 39 | Up | 1.60E-06 | 3.82E-05 |
| REACTOME_AURKA_ACTIVATION_BY_TPX2 | 21 | Up | 1.62E-06 | 3.86E-05 |
| REACTOME_REGULATION_OF_TP53_ACTIVITY_THROUGH_PHOSPHORYLATION | 22 | Up | 1.71E-06 | 4.03E-05 |
| VERNELL_RETINOBLASTOMA_PATHWAY_UP | 33 | Up | 1.71E-06 | 4.03E-05 |
| SENGUPTA_NASOPHARYNGEAL_CARCINOMA_UP | 95 | Up | 1.82E-06 | 4.27E-05 |
| REACTOME_G2_M_DNA_REPLICATION_CHECKPOINT | 4 | Up | 1.94E-06 | 4.52E-05 |
| GRADE_COLON_AND_RECTAL_CANCER_UP | 43 | Up | 2.16E-06 | 5.00E-05 |
| REACTOME_ACTIVATION_OF_THE_PRE_REPLICATIVE_COMPLEX | 20 | Up | 2.17E-06 | 5.00E-05 |
| REACTOME_DNA_REPLICATION_PRE_INITIATION | 20 | Up | 2.17E-06 | 5.00E-05 |
| KUMAMOTO_RESPONSE_TO_NUTLIN_3A_DN | 6 | Up | 2.24E-06 | 5.12E-05 |
| REACTOME_SUMOYLATION_OF_DNA_REPLICATION_PROTEINS | 7 | Up | 2.68E-06 | 6.10E-05 |
| RHEIN_ALL_GLUCOCORTICOID_THERAPY_DN | 79 | Up | 2.95E-06 | 6.68E-05 |
| REACTOME_ACTIVATION_OF_ATR_IN_RESPONSE_TO_REPLICATION_STRESS | 20 | Up | 2.97E-06 | 6.69E-05 |
| MODY_HIPPOCAMPUS_PRENATAL | 4 | Up | 3.15E-06 | 7.07E-05 |
| REACTOME_INTRA_GOLGI_AND_RETROGRADE_GOLGI_TO_ER_TRAFFIC | 24 | Up | 3.21E-06 | 7.17E-05 |
| SERVITJA_LIVER_HNF1A_TARGETS_UP | 42 | Up | 3.90E-06 | 8.68E-05 |

| | | | | |
|---|---|---|---|---|
| REACTOME_GOLGI_CISTERNAE_PERICENTRIOLAR_STACK_REORGANIZATION | 4 | Up | 4.79E-06 | 0.000105 |
| REACTOME_ACTIVATION_OF_NIMA_KINASES_NEK9_NEK6_NEK7 | 4 | Up | 4.79E-06 | 0.000105 |
| REACTOME_THE_ROLE_OF_GTSE1_IN_G2_M_PROGRESSION_AFTER_G2_CHECKPOINT | 4 | Up | 4.79E-06 | 0.000105 |
| GRAHAM_CML_QUIESCENT_VS_CML_DIVIDING_DN | 4 | Up | 5.54E-06 | 0.0001209 |
| REACTOME_FACTORS_INVOLVED_IN_MEGAKARYOCYTE_DEVELOPMENT_AND_PLATELET_PRODUCTION | 25 | Up | 7.21E-06 | 0.0001567 |
| ZHU_SKIL_TARGETS_DN | 4 | Up | 7.35E-06 | 0.000159 |
| WHITFIELD_CELL_CYCLE_G1_S | 34 | Up | 9.76E-06 | 0.0002103 |
| FISCHER_G1_S_CELL_CYCLE | 53 | Up | 1.02E-05 | 0.0002176 |
| WIELAND_UP_BY_HBV_INFECTION | 10 | Up | 1.04E-05 | 0.0002219 |
| SMID_BREAST_CANCER_BASAL_UP | 152 | Up | 1.07E-05 | 0.0002278 |
| REACTOME_INITIATION_OF_NUCLEAR_ENVELOPE_NE_REFORMATION | 6 | Up | 1.12E-05 | 0.0002361 |
| PYEON_CANCER_HEAD_AND_NECK_VS_CERVICAL_UP | 41 | Up | 1.13E-05 | 0.0002361 |
| FOURNIER_ACINAR_DEVELOPMENT_LATE_DN | 10 | Up | 1.13E-05 | 0.0002361 |
| REACTOME_UNWINDING_OF_DNA | 9 | Up | 1.32E-05 | 0.000275 |
| LEE_LIVER_CANCER_SURVIVAL_DN | 33 | Up | 1.38E-05 | 0.0002865 |
| REACTOME_HDR_THROUGH_HOMOLOGOUS_RECOMBINATION_HRR | 25 | Up | 1.49E-05 | 0.0003085 |
| REACTOME_SWITCHING_OF_ORIGINS_TO_A_POST_REPLICATIVE_STATE | 16 | Up | 1.53E-05 | 0.0003156 |
| BHATI_G2M_ARREST_BY_2METHOXYESTRADIOL_UP | 28 | Up | 1.66E-05 | 0.0003404 |
| BIOCARTA_EFP_PATHWAY | 3 | Up | 1.76E-05 | 0.0003604 |
| BIOCARTA_CELLCYCLE_PATHWAY | 6 | Up | 1.83E-05 | 0.0003706 |
| GEORGES_TARGETS_OF_MIR192_AND_MIR215 | 189 | Up | 1.83E-05 | 0.0003706 |
| GROSS_HYPOXIA_VIA_ELK3_UP | 41 | Up | 1.87E-05 | 0.0003774 |
| REACTOME_ORC1_REMOVAL_FROM_CHROMATIN | 13 | Up | 1.96E-05 | 0.0003934 |
| STEIN_ESR1_TARGETS | 25 | Up | 2.04E-05 | 0.0004077 |
| SUNG_METASTASIS_STROMA_DN | 17 | Up | 2.08E-05 | 0.0004136 |
| SPIELMAN_LYMPHOBLAST_EUROPEAN_VS_ASIAN_UP | 39 | Up | 2.13E-05 | 0.0004214 |
| REACTOME_PROCESSING_OF_DNA_DOUBLE_STRAND_BREAK_ENDS | 20 | Up | 2.39E-05 | 0.0004723 |
| REACTOME_G2_M_DNA_DAMAGE_CHECKPOINT | 19 | Up | 2.49E-05 | 0.000489 |
| SHIPP_DLBCL_VS_FOLLICULAR_LYMPHOMA_UP | 6 | Up | 2.63E-05 | 0.0005142 |
| JOHANSSON_GLIOMAGENESIS_BY_PDGFB_UP | 21 | Up | 2.85E-05 | 0.0005562 |
| REACTOME_TP53_REGULATES_TRANSCRIPTION_OF_CELL_CYCLE_GENES | 12 | Up | 3.05E-05 | 0.0005914 |
| ANDERSEN_CHOLANGIOCARCINOMA_CLASS2 | 38 | Up | 3.32E-05 | 0.0006424 |
| REACTOME_PHOSPHORYLATION_OF_EMI1 | 5 | Up | 3.44E-05 | 0.0006633 |
| GARCIA_TARGETS_OF_FLI1_AND_DAX1_DN | 33 | Up | 5.04E-05 | 0.0009671 |
| BIOCARTA_RANMS_PATHWAY | 4 | Up | 5.21E-05 | 0.000995 |
| REACTOME_POTASSIUM_CHANNELS | 17 | Down | 5.39E-05 | 0.0010259 |
| REACTOME_MITOTIC_TELOPHASE_CYTOKINESIS | 4 | Up | 5.50E-05 | 0.0010427 |

| | | | | |
|---|---|---|---|---|
| INAMURA_LUNG_CANCER_SCC_UP | 3 | Up | 5.92E-05 | 0.0011172 |
| SIMBULAN_PARP1_TARGETS_DN | 7 | Up | 5.96E-05 | 0.0011207 |
| BURTON_ADIPOGENESIS_PEAK_AT_16HR | 15 | Up | 6.01E-05 | 0.0011261 |
| BIOCARTA_MPR_PATHWAY | 4 | Up | 6.24E-05 | 0.0011594 |
| REACTOME_APC_CDC20_MEDIATED_DEGRADATION_OF_NEK2A | 6 | Up | 6.24E-05 | 0.0011594 |
| BENPORATH_MYC_MAX_TARGETS | 65 | Up | 6.33E-05 | 0.0011721 |
| REACTOME_PROTEIN_PROTEIN_INTERACTIONS_AT_SYNAPSES | 25 | Down | 6.66E-05 | 0.0012286 |
| MIKKELSEN_ES_LCP_WITH_H3K27ME3 | 2 | Down | 6.93E-05 | 0.0012731 |
| DANG_BOUND_BY_MYC | 88 | Up | 7.02E-05 | 0.0012845 |
| COWLING_MYCN_TARGETS | 12 | Down | 7.62E-05 | 0.0013846 |
| REACTOME_ASSEMBLY_OF_THE_PRE_REPLICATIVE_COMPLEX | 11 | Up | 7.64E-05 | 0.0013846 |
| GEORGES_CELL_CYCLE_MIR192_TARGETS | 20 | Up | 7.66E-05 | 0.0013846 |
| RHODES_CANCER_META_SIGNATURE | 11 | Up | 8.45E-05 | 0.0015222 |
| WANG_METASTASIS_OF_BREAST_CANCER_ESR1_UP | 6 | Up | 8.81E-05 | 0.0015817 |
| MCBRYAN_PUBERTAL_BREAST_6_7WK_DN | 27 | Up | 9.06E-05 | 0.0016196 |
| REACTOME_NUCLEAR_ENVELOPE_NE_REASSEMBLY | 8 | Up | 9.26E-05 | 0.001649 |
| SLEBOS_HEAD_AND_NECK_CANCER_WITH_HPV_UP | 19 | Up | 9.36E-05 | 0.0016616 |
| REACTOME_APC_C_CDH1_MEDIATED_DEGRADATION_OF_CDC20_AND_OTHER_APC_C_CDH1_TARGETED_PROTEINS_IN_LATE_MITOSIS_EARLY_G1 | 9 | Up | 0.000102 | 0.0017959 |
| THILLAINADESAN_ZNF217_TARGETS_UP | 12 | Up | 0.000102 | 0.0018023 |
| PEART_HDAC_PROLIFERATION_CLUSTER_DN | 16 | Up | 0.000107 | 0.0018655 |
| LY_AGING_PREMATURE_DN | 10 | Up | 0.000107 | 0.0018655 |
| GENTLES_LEUKEMIC_STEM_CELL_DN | 4 | Up | 0.000108 | 0.0018819 |
| REACTOME_CHROMOSOME_MAINTENANCE | 22 | Up | 0.000111 | 0.0019309 |
| LEE_BMP2_TARGETS_DN | 154 | Up | 0.000117 | 0.002018 |
| BOYAULT_LIVER_CANCER_SUBCLASS_G123_UP | 10 | Up | 0.000117 | 0.002018 |
| OHASHI_AURKB_TARGETS | 4 | Up | 0.000119 | 0.0020417 |
| CROSBY_E2F4_TARGETS | 6 | Up | 0.000152 | 0.0025965 |
| REACTOME_RECRUITMENT_OF_MITOTIC_CENTROSOME_PROTEINS_AND_COMPLEXES | 19 | Up | 0.000155 | 0.0026334 |
| REACTOME_RECRUITMENT_OF_NUMA_TO_MITOTIC_CENTROSOMES | 19 | Up | 0.000155 | 0.0026334 |
| ZHAN_MULTIPLE_MYELOMA_MF_UP | 4 | Down | 0.000173 | 0.0029317 |
| REACTOME_NUCLEAR_PORE_COMPLEX_NPC_DISASSEMBLY | 5 | Up | 0.000175 | 0.0029449 |
| WANG_CISPLATIN_RESPONSE_AND_XPC_UP | 42 | Up | 0.000185 | 0.0031146 |
| REACTOME_ANCHORING_OF_THE_BASAL_BODY_TO_THE_PLASMA_MEMBRANE | 21 | Up | 0.000196 | 0.0032812 |
| CAFFAREL_RESPONSE_TO_THC_24HR_5_DN | 7 | Up | 0.000199 | 0.003327 |
| REACTOME_G0_AND_EARLY_G1 | 11 | Up | 0.000211 | 0.0035118 |
| LOPEZ_MESOTELIOMA_SURVIVAL_TIME_UP | 7 | Up | 0.000214 | 0.0035427 |
| BIOCARTA_AKAP95_PATHWAY | 3 | Up | 0.000216 | 0.0035644 |
| WEI_MYCN_TARGETS_WITH_E_BOX | 114 | Up | 0.000217 | 0.0035756 |
| MEINHOLD_OVARIAN_CANCER_LOW_GRADE_DN | 5 | Up | 0.000219 | 0.0035843 |
| HAMAI_APOPTOSIS_VIA_TRAIL_UP | 126 | Up | 0.000219 | 0.0035843 |

| | | | | |
|---|---|---|---|---|
| WILLIAMS_ESR1_TARGETS_UP | 8 | Up | 0.000222 | 0.0036223 |
| MARTENS_TRETINOIN_RESPONSE_DN | 59 | Up | 0.000238 | 0.0038572 |
| RIGGI_EWING_SARCOMA_PROGENITOR_UP | 74 | Down | 0.000252 | 0.0040783 |
| REACTOME_DEUBIQUITINATION | 16 | Up | 0.000266 | 0.0042908 |
| TURASHVILI_BREAST_LOBULAR_CARCINOMA_VS_LOBULAR_NORMAL_UP | 21 | Down | 0.000283 | 0.004553 |
| BIOCARTA_MCM_PATHWAY | 9 | Up | 0.000332 | 0.0053151 |
| PYEON_HPV_POSITIVE_TUMORS_UP | 27 | Up | 0.000348 | 0.0055534 |
| PEART_HDAC_PROLIFERATION_CLUSTER_UP | 10 | Up | 0.000351 | 0.0055834 |
| REACTOME_RESOLUTION_OF_D_LOOP_STRUCTURES | 16 | Up | 0.000353 | 0.0055978 |
| NAKAMURA_LUNG_CANCER | 4 | Up | 0.000363 | 0.005728 |
| REACTOME_HOMOLOGOUS_DNA_PAIRING_AND_STRAND_EXCHANGE | 20 | Up | 0.000373 | 0.0058765 |
| REACTOME_NUCLEAR_ENVELOPE_BREAKDOWN | 9 | Up | 0.000377 | 0.0059216 |
| CHICAS_RB1_TARGETS_GROWING | 94 | Up | 0.000382 | 0.0059717 |
| CONCANNON_APOPTOSIS_BY_EPOXOMICIN_DN | 48 | Up | 0.000389 | 0.0060664 |
| GOZGIT_ESR1_TARGETS_DN | 120 | Down | 0.000393 | 0.0061067 |
| RAY_TUMORIGENESIS_BY_ERBB2_CDC25A_UP | 31 | Up | 0.000396 | 0.0061309 |
| HONRADO_BREAST_CANCER_BRCA1_VS_BRCA2 | 6 | Up | 0.000418 | 0.006449 |
| CHIARETTI_T_ALL_RELAPSE_PROGNOSIS | 6 | Up | 0.00044 | 0.0067698 |
| YAO_TEMPORAL_RESPONSE_TO_PROGESTERONE_CLUSTER_15 | 10 | Up | 0.000446 | 0.0068409 |
| TARTE_PLASMA_CELL_VS_B_LYMPHOCYTE_DN | 2 | Down | 0.000448 | 0.0068517 |
| REACTOME_TP53_REGULATES_TRANSCRIPTION_OF_GENES_INVOLVED_IN_G2_CELL_CYCLE_ARREST | 7 | Up | 0.000459 | 0.0069824 |
| REACTOME_NEGATIVE_REGULATION_OF_NOTCH4_SIGNALING | 1 | Up | 0.00046 | 0.0069824 |
| PID_P73PATHWAY | 12 | Up | 0.000465 | 0.0070318 |
| GLINSKY_CANCER_DEATH_UP | 3 | Up | 0.000482 | 0.0072665 |
| REACTOME_HDR_THROUGH_SINGLE_STRAND_ANNEALING_SSA | 16 | Up | 0.000555 | 0.0083442 |
| JAEGER_METASTASIS_UP | 16 | Up | 0.000576 | 0.0086399 |
| KOKKINAKIS_METHIONINE_DEPRIVATION_96HR_DN | 14 | Up | 0.000579 | 0.0086549 |
| CAFFAREL_RESPONSE_TO_THC_DN | 7 | Up | 0.000594 | 0.0088442 |
| PID_ATM_PATHWAY | 9 | Up | 0.000601 | 0.0089216 |
| REACTOME_APC_C_CDC20_MEDIATED_DEGRADATION_OF_CYCLIN_B | 6 | Up | 0.000634 | 0.009382 |
| FERRANDO_HOX11_NEIGHBORS | 5 | Up | 0.000671 | 0.0099071 |
| SCHMIDT_POR_TARGETS_IN_LIMB_BUD_UP | 3 | Down | 0.000704 | 0.0103631 |
| ZAMORA_NOS2_TARGETS_UP | 12 | Up | 0.000714 | 0.0104376 |
| STANHILL_HRAS_TRANSFROMATION_UP | 1 | Down | 0.000716 | 0.0104376 |
| KYNG_RESPONSE_TO_H2O2_VIA_ERCC6 | 1 | Down | 0.000716 | 0.0104376 |
| ZHANG_BREAST_CANCER_PROGENITORS_UP | 118 | Up | 0.000745 | 0.0108313 |
| REACTOME_ADHERENS_JUNCTIONS_INTERACTIONS | 5 | Down | 0.000772 | 0.0111858 |
| QI_HYPOXIA | 21 | Down | 0.000776 | 0.0111992 |
| BIOCARTA_G2_PATHWAY | 9 | Up | 0.000778 | 0.0111992 |
| VANTVEER_BREAST_CANCER_POOR_PROGNOSIS | 12 | Up | 0.00081 | 0.0116295 |
| DUTERTRE_ESTRADIOL_RESPONSE_6HR_UP | 50 | Up | 0.000821 | 0.0117532 |

| | | | | |
|---|---|---|---|---|
| ABRAMSON_INTERACT_WITH_AIRE | 10 | Up | 0.000827 | 0.0117973 |
| REACTOME_INTERACTION_BETWEEN_L1_AND_ANKYRINS | 3 | Down | 0.000832 | 0.0118332 |
| REACTOME_REGULATION_OF_TP53_EXPRESSION_AND_DEGRADATION | 4 | Up | 0.000834 | 0.0118372 |
| BROWNE_HCMV_INFECTION_2HR_DN | 9 | Up | 0.000837 | 0.0118384 |
| REACTOME_RESOLUTION_OF_D_LOOP_STRUCTURES_THROUGH_SYNTHESIS_DEPENDENT_STRAND_ANNEALING_SDSA | 15 | Up | 0.00088 | 0.0124077 |
| LEE_TARGETS_OF_PTCH1_AND_SUFU_UP | 7 | Up | 0.000888 | 0.0124848 |
| PID_FANCONI_PATHWAY | 13 | Up | 0.000896 | 0.0125569 |
| KEGG_BASE_EXCISION_REPAIR | 8 | Up | 0.000991 | 0.0138523 |
| MARKS_HDAC_TARGETS_DN | 2 | Up | 0.001011 | 0.0140517 |
| BAKER_HEMATOPOIESIS_STAT3_TARGETS | 2 | Up | 0.001011 | 0.0140517 |
| CUI_TCF21_TARGETS_2_UP | 106 | Up | 0.001031 | 0.0142869 |
| REACTOME_CHK1_CHK2_CDS1_MEDIATED_INACTIVATION_OF_CYCLIN_B_CDK1_COMPLEX | 4 | Up | 0.001043 | 0.014409 |
| LEE_LIVER_CANCER_MYC_E2F1_UP | 10 | Up | 0.001048 | 0.0144291 |
| REACTOME_INHIBITION_OF_THE_PROTEOLYTIC_ACTIVITY_OF_APC_C_REQUIRED_FOR_THE_ONSET_OF_ANAPHASE_BY_MITOTIC_SPINDLE_CHECKPOINT_COMPONENTS | 5 | Up | 0.001076 | 0.0147754 |
| REACTOME_HSF1_DEPENDENT_TRANSACTIVATION | 5 | Down | 0.001113 | 0.0152456 |
| PUIFFE_INVASION_INHIBITED_BY_ASCITES_UP | 8 | Up | 0.001122 | 0.0153225 |
| KRIEG_HYPOXIA_NOT_VIA_KDM3A | 108 | Up | 0.001137 | 0.0154817 |
| KINSEY_TARGETS_OF_EWSR1_FLII_FUSION_DN | 60 | Down | 0.001166 | 0.0158333 |
| TURASHVILI_BREAST_DUCTAL_CARCINOMA_VS_LOBULAR_NORMAL_DN | 13 | Down | 0.001185 | 0.0160126 |
| WONG_PROTEASOME_GENE_MODULE | 4 | Up | 0.001186 | 0.0160126 |
| BHATTACHARYA_EMBRYONIC_STEM_CELL | 19 | Up | 0.001216 | 0.0163657 |
| RIZ_ERYTHROID_DIFFERENTIATION_CCNE1 | 8 | Up | 0.001244 | 0.0166905 |
| REACTOME_NUCLEAR_RECEPTOR_TRANSCRIPTION_PATHWAY | 9 | Down | 0.001264 | 0.0169183 |
| MASRI_RESISTANCE_TO_TAMOXIFEN_AND_AROMATASE_INHIBITORS_UP | 5 | Down | 0.001278 | 0.0170209 |
| PID_ATF2_PATHWAY | 3 | Up | 0.001279 | 0.0170209 |
| REACTOME_PHOSPHORYLATION_OF_THE_APC_C | 6 | Up | 0.001295 | 0.0171899 |
| YU_BAP1_TARGETS | 14 | Up | 0.001343 | 0.0177806 |
| REACTOME_MASTL_FACILITATES_MITOTIC_PROGRESSION | 3 | Up | 0.001378 | 0.0181881 |
| BORCZUK_MALIGNANT_MESOTHELIOMA_UP | 38 | Up | 0.001413 | 0.018601 |
| KALMA_E2F1_TARGETS | 4 | Up | 0.001473 | 0.0193339 |
| SMID_BREAST_CANCER_RELAPSE_IN_BRAIN_UP | 10 | Up | 0.001538 | 0.0201322 |
| REACTOME_CELL_CELL_COMMUNICATION | 14 | Down | 0.001582 | 0.0206532 |
| WONG_ADULT_TISSUE_STEM_MODULE | 122 | Down | 0.001594 | 0.0207509 |
| ZHENG_GLIOBLASTOMA_PLASTICITY_DN | 18 | Down | 0.001615 | 0.0209622 |
| REACTOME_CLASS_A_1_RHODOPSIN_LIKE_RECEPTORS | 30 | Down | 0.00164 | 0.0212355 |
| CHEN_ETV5_TARGETS_TESTIS | 8 | Up | 0.001657 | 0.0213941 |
| KEGG_NEUROACTIVE_LIGAND_RECEPTOR_INTERACTION | 39 | Down | 0.001688 | 0.021728 |
| BERTUCCI_MEDULLARY_VS_DUCTAL_BREAST_CANCER_UP | 15 | Up | 0.001698 | 0.0218086 |

| | | | | |
|---|---|---|---|---|
| REACTOME_SENESCENCE_ASSOCIATED_SECRETORY_PHENOTYPE_SASP | 7 | Up | 0.001778 | 0.0227632 |
| REACTOME_G1_S_SPECIFIC_TRANSCRIPTION | 13 | Up | 0.001797 | 0.0229492 |
| BIOCARTA_BARD1_PATHWAY | 4 | Up | 0.001809 | 0.0230079 |
| REACTOME_ESTABLISHMENT_OF_SISTER_CHROMATID_COHESION | 3 | Up | 0.001811 | 0.0230079 |
| REACTOME_CELL_CELL_JUNCTION_ORGANIZATION | 8 | Down | 0.001845 | 0.0233679 |
| REACTOME_TELOMERE_C_STRAND_LAGGING_STRAND_SYNTHESIS | 9 | Up | 0.001908 | 0.0241074 |
| VANHARANTA_UTERINE_FIBROID_UP | 17 | Down | 0.001954 | 0.0246217 |
| REICHERT_G1S_REGULATORS_AS_PI3K_TARGETS | 3 | Up | 0.002029 | 0.0252834 |
| BIOCARTA_AKAPCENTROSOME_PATHWAY | 1 | Up | 0.002036 | 0.0252834 |
| BIOCARTA_SAM68_PATHWAY | 1 | Up | 0.002036 | 0.0252834 |
| REACTOME_MAPK3_ERK1_ACTIVATION | 1 | Up | 0.002036 | 0.0252834 |
| REACTOME_TELOMERE_MAINTENANCE | 11 | Up | 0.002038 | 0.0252834 |
| REACTOME_EXTENSION_OF_TELOMERES | 11 | Up | 0.002038 | 0.0252834 |
| WANG_IMMORTALIZED_BY_HOXA9_AND_MEIS1_DN | 4 | Down | 0.002085 | 0.0257959 |
| SMID_BREAST_CANCER_NORMAL_LIKE_UP | 68 | Down | 0.002091 | 0.0257995 |
| JEON_SMAD6_TARGETS_DN | 7 | Up | 0.002141 | 0.0263488 |
| NUNODA_RESPONSE_TO_DASATINIB_IMATINIB_UP | 5 | Up | 0.002224 | 0.0273018 |
| WALLACE_JAK2_TARGETS_UP | 3 | Up | 0.002351 | 0.028789 |
| BIOCARTA_RB_PATHWAY | 5 | Up | 0.002566 | 0.0312585 |
| BIOCARTA_CDC25_PATHWAY | 5 | Up | 0.002566 | 0.0312585 |
| BENPORATH_ES_2 | 13 | Up | 0.002647 | 0.0321585 |
| MCCLUNG_DELTA_FOSB_TARGETS_2WK | 4 | Down | 0.002661 | 0.0322413 |
| SEMBA_FHIT_TARGETS_DN | 6 | Up | 0.002755 | 0.0333048 |
| BIOCARTA_ATRBRCA_PATHWAY | 9 | Up | 0.002773 | 0.0334342 |
| DAIRKEE_CANCER_PRONE_RESPONSE_BPA_E2 | 14 | Up | 0.002839 | 0.0341318 |
| PID_BARD1_PATHWAY | 13 | Up | 0.002846 | 0.0341318 |
| KIM_ALL_DISORDERS_CALB1_CORR_UP | 25 | Down | 0.002892 | 0.034607 |
| WANG_SMARCE1_TARGETS_UP | 84 | Down | 0.00295 | 0.0352086 |
| VECCHI_GASTRIC_CANCER_ADVANCED_VS_EARLY_UP | 53 | Down | 0.002967 | 0.0353151 |
| ODONNELL_TARGETS_OF_MYC_AND_TFRC_UP | 6 | Down | 0.002998 | 0.0355952 |
| NADLER_OBESITY_UP | 5 | Down | 0.003014 | 0.0356992 |
| CUI_GLUCOSE_DEPRIVATION | 9 | Up | 0.003021 | 0.0356992 |
| JOHANSSON_BRAIN_CANCER_EARLY_VS_LATE_DN | 8 | Down | 0.003029 | 0.035702 |
| REACTOME_ADAPTIVE_IMMUNE_SYSTEM | 48 | Up | 0.003093 | 0.0363668 |
| REACTOME_RAF_INDEPENDENT_MAPK1_3_ACTIVATION | 2 | Up | 0.003128 | 0.0366881 |
| BIOCARTA_DNAFRAGMENT_PATHWAY | 2 | Up | 0.003187 | 0.0372816 |
| REACTOME_VOLTAGE_GATED_POTASSIUM_CHANNELS | 8 | Down | 0.003224 | 0.0376206 |
| REACTOME_CELL_JUNCTION_ORGANIZATION | 10 | Down | 0.003441 | 0.040018 |
| PARENT_MTOR_SIGNALING_DN | 8 | Down | 0.003446 | 0.040018 |
| RIZ_ERYTHROID_DIFFERENTIATION_HBZ | 7 | Up | 0.003471 | 0.0402113 |
| BRUNO_HEMATOPOIESIS | 10 | Down | 0.003558 | 0.0411169 |
| AUNG_GASTRIC_CANCER | 6 | Up | 0.003635 | 0.0419089 |

| | | | | |
|---|---|---|---|---|
| DURAND_STROMA_S_UP | 60 | Down | 0.003645 | 0.041921 |
| REACTOME_DISEASES_OF_PROGRAMMED_CELL_DEATH | 2 | Up | 0.003694 | 0.0423758 |
| GERY_CEBP_TARGETS | 17 | Down | 0.003779 | 0.043251 |
| AMIT_EGF_RESPONSE_60_HELA | 4 | Down | 0.003792 | 0.0432953 |
| DELYS_THYROID_CANCER_DN | 63 | Down | 0.003947 | 0.0449589 |
| OHASHI_AURKA_TARGETS | 1 | Up | 0.003981 | 0.0452358 |
| GROSS_HYPOXIA_VIA_ELK3_ONLY_DN | 5 | Up | 0.004019 | 0.045552 |
| BIOCARTA_STATHMIN_PATHWAY | 2 | Up | 0.004185 | 0.0473188 |
| CAFFAREL_RESPONSE_TO_THC_24HR_5_UP | 4 | Up | 0.004223 | 0.0476385 |
| SUNG_METASTASIS_STROMA_UP | 21 | Down | 0.004259 | 0.0478903 |
| REACTOME_DNA_DAMAGE_BYPASS | 8 | Up | 0.004265 | 0.0478903 |
| KEGG_CALCIUM_SIGNALING_PATHWAY | 20 | Down | 0.004294 | 0.0480923 |
| CHEMELLO_SOLEUS_VS_EDL_MYOFIBERS_UP | 2 | Down | 0.004311 | 0.0481719 |
| SMID_BREAST_CANCER_BASAL_DN | 97 | Down | 0.004344 | 0.048431 |
| MARTINEZ_RESPONSE_TO_TRABECTEDIN_DN | 41 | Up | 0.004388 | 0.0488029 |
| LUI_THYROID_CANCER_CLUSTER_5 | 2 | Down | 0.004409 | 0.0489187 |
| MYLLYKANGAS_AMPLIFICATION_HOT_SPOT_12 | 2 | Up | 0.004419 | 0.0489195 |
| KORKOLA_TERATOMA | 15 | Up | 0.004522 | 0.0499466 |
| XU_RESPONSE_TO_TRETINOIN_UP | 2 | Up | 0.004731 | 0.0521248 |
| WU_APOPTOSIS_BY_CDKN1A_NOT_VIA_TP53 | 4 | Up | 0.004744 | 0.0521518 |
| LEE_LIVER_CANCER_MYC_UP | 11 | Up | 0.004811 | 0.0527603 |
| SCHAEFFER_PROSTATE_DEVELOPMENT_48HR_DN | 76 | Down | 0.004869 | 0.0532733 |
| REACTOME_RUNX1_REGULATES_TRANSCRIPTION_OF_GENES_INVOLVED_IN_DIFFERENTIATION_OF_MYELOID_CELLS | 2 | Down | 0.004958 | 0.0540687 |
| IWANAGA_E2F1_TARGETS_INDUCED_BY_SERUM | 9 | Up | 0.004964 | 0.0540687 |
| REACTOME_TRANSCRIPTION_OF_E2F_TARGETS_UNDER_NEGATIVE_CONTROL_BY_P107_RBL1_AND_P130_RBL2_IN_COMPLEX_WITH_HDAC1 | 8 | Up | 0.005159 | 0.0560605 |
| HASLINGER_B_CLL_WITH_MUTATED_VH_GENES | 4 | Down | 0.005196 | 0.0562364 |
| RICKMAN_TUMOR_DIFFERENTIATED_WELL_VS_MODERATELY_UP | 14 | Down | 0.005202 | 0.0562364 |
| REACTOME_AMINE_LIGAND_BINDING_RECEPTORS | 4 | Down | 0.005222 | 0.0562364 |
| MATTIOLI_MGUS_VS_PCL | 18 | Up | 0.005222 | 0.0562364 |
| SA_REG_CASCADE_OF_CYCLIN_EXPR | 2 | Up | 0.005249 | 0.0562614 |
| REACTOME_G2_PHASE | 2 | Up | 0.005249 | 0.0562614 |
| KIM_ALL_DISORDERS_OLIGODENDROCYTE_NUMBER_CORR_UP | 45 | Down | 0.005355 | 0.0572718 |
| MIKKELSEN_DEDIFFERENTIATED_STATE_UP | 1 | Down | 0.005399 | 0.0574877 |
| LOPEZ_EPITHELIOID_MESOTHELIOMA | 1 | Down | 0.005399 | 0.0574877 |
| YAMAZAKI_TCEB3_TARGETS_DN | 39 | Up | 0.005414 | 0.0575106 |
| BURTON_ADIPOGENESIS_12 | 3 | Up | 0.005468 | 0.0579629 |
| REACTOME_TP53_REGULATES_TRANSCRIPTION_OF_GENES_INVOLVED_IN_G1_CELL_CYCLE_ARREST | 3 | Up | 0.00555 | 0.0586982 |
| ELVIDGE_HYPOXIA_BY_DMOG_UP | 28 | Down | 0.005594 | 0.0590302 |
| REACTOME_PROCESSIVE_SYNTHESIS_ON_THE_LAGGING_STRAND | 6 | Up | 0.005697 | 0.0599834 |

| | | | | |
|---|---|---|---|---|
| VECCHI_GASTRIC_CANCER_EARLY_DN | 75 | Down | 0.005712 | 0.0600115 |
| REACTOME_OVARIAN_TUMOR_DOMAIN_PROTEASES | 2 | Up | 0.005745 | 0.0602168 |
| LI_CISPLATIN_RESISTANCE_DN | 12 | Down | 0.005771 | 0.0603636 |
| REACTOME_RECOGNITION_OF_DNA_DAMAGE_BY_PCNA_CONTAINING_REPLICATION_COMPLEX | 7 | Up | 0.005832 | 0.0608597 |
| REACTOME_TRANSCRIPTIONAL_REGULATION_BY_MECP2 | 5 | Up | 0.006077 | 0.0632826 |
| REACTOME_LAGGING_STRAND_SYNTHESIS | 7 | Up | 0.006152 | 0.0638038 |
| RODRIGUES_THYROID_CARCINOMA_DN | 13 | Down | 0.006154 | 0.0638038 |
| BIOCARTA_AGR_PATHWAY | 6 | Down | 0.006208 | 0.0640999 |
| PRAMOONJAGO_SOX4_TARGETS_DN | 12 | Up | 0.00621 | 0.0640999 |
| CHUANG_OXIDATIVE_STRESS_RESPONSE_DN | 5 | Up | 0.006308 | 0.0649766 |
| REACTOME_BASE_EXCISION_REPAIR | 11 | Up | 0.006366 | 0.0654287 |
| REACTOME_G_ALPHA_Q_SIGNALLING_EVENTS | 22 | Down | 0.006465 | 0.0662966 |
| REACTOME_E2F_MEDIATED_REGULATION_OF_DNA_REPLICATION | 9 | Up | 0.006506 | 0.0665748 |
| REACTOME_SUMOYLATION | 16 | Up | 0.006559 | 0.066978 |
| PICCALUGA_ANGIOIMMUNOBLASTIC_LYMPHOMA_UP | 64 | Down | 0.006612 | 0.067374 |
| FLECHNER_PBL_KIDNEY_TRANSPLANT_OK_VS_DONOR_UP | 8 | Down | 0.00663 | 0.067379 |
| BENPORATH_ES_CORE_NINE_CORRELATED | 11 | Up | 0.006641 | 0.067379 |
| REACTOME_NEUREXINS_AND_NEUROLIGINS | 12 | Down | 0.00671 | 0.0678249 |
| GOUYER_TUMOR_INVASIVENESS | 1 | Down | 0.006742 | 0.0678249 |
| CHEN_ETV5_TARGETS_SERTOLI | 1 | Down | 0.006742 | 0.0678249 |
| MILICIC_FAMILIAL_ADENOMATOUS_POLYPOSIS_UP | 1 | Down | 0.006742 | 0.0678249 |
| LI_WILMS_TUMOR_VS_FETAL_KIDNEY_2_DN | 14 | Down | 0.006961 | 0.0698775 |
| REACTOME_OXIDATIVE_STRESS_INDUCED_SENESCENCE | 2 | Up | 0.007029 | 0.0702627 |
| REACTOME_ONCOGENE_INDUCED_SENESCENCE | 2 | Up | 0.007029 | 0.0702627 |
| ROSS_AML_OF_FAB_M7_TYPE | 13 | Down | 0.007206 | 0.0718768 |
| PID_RXR_VDR_PATHWAY | 4 | Down | 0.007259 | 0.0722383 |
| KARAKAS_TGFB1_SIGNALING | 7 | Up | 0.007273 | 0.0722383 |
| IZADPANAH_STEM_CELL_ADIPOSE_VS_BONE_DN | 26 | Down | 0.007387 | 0.0731176 |
| RAY_TARGETS_OF_P210_BCR_ABL_FUSION_DN | 4 | Down | 0.007392 | 0.0731176 |
| GRAHAM_CML_QUIESCENT_VS_CML_DIVIDING_UP | 1 | Down | 0.007541 | 0.0739763 |
| DACOSTA_UV_RESPONSE_VIA_ERCC3_XPCS_UP | 1 | Down | 0.007541 | 0.0739763 |
| ALONSO_METASTASIS_EMT_DN | 1 | Down | 0.007541 | 0.0739763 |
| HOLLEMAN_DAUNORUBICIN_B_ALL_UP | 1 | Down | 0.007541 | 0.0739763 |
| REACTOME_RESOLUTION_OF_AP_SITES_VIA_THE_MULTIPLE_NUCLEOTIDE_PATCH_REPLACEMENT_PATHWAY | 7 | Up | 0.00765 | 0.0748857 |
| CAIRO_HEPATOBLASTOMA_POOR_SURVIVAL | 5 | Up | 0.007682 | 0.0750434 |
| KRASNOSELSKAYA_ILF3_TARGETS_DN | 17 | Up | 0.007728 | 0.0753361 |
| CHICAS_RB1_TARGETS_LOW_SERUM | 14 | Up | 0.007913 | 0.0769851 |
| REACTOME_CELLULAR_SENESCENCE | 14 | Up | 0.007953 | 0.077026 |
| HORTON_SREBF_TARGETS | 5 | Down | 0.007966 | 0.077026 |
| CUI_TCF21_TARGETS_2_DN | 133 | Down | 0.007966 | 0.077026 |
| REACTOME_PCNA_DEPENDENT_LONG_PATCH_BASE_EXCISION_REPAIR | 6 | Up | 0.007993 | 0.0771313 |

| | | | | |
|---|---|---|---|---|
| REACTOME_CARDIAC_CONDUCTION | 15 | Down | 0.008057 | 0.0775873 |
| SCIAN_INVERSED_TARGETS_OF_TP53_AND_TP73_UP | 1 | Down | 0.008084 | 0.0776886 |
| RAMPON_ENRICHED_LEARNING_ENVIRONMENT_LATE_UP | 1 | Down | 0.008316 | 0.0794399 |
| REACTOME_VLDLR_INTERNALISATION_AND_DEGRADATION | 1 | Down | 0.008316 | 0.0794399 |
| REACTOME_VLDL_CLEARANCE | 1 | Down | 0.008316 | 0.0794399 |
| GENTILE_RESPONSE_CLUSTER_D3 | 6 | Up | 0.008483 | 0.0805817 |
| REACTOME_TANDEM_PORE_DOMAIN_POTASSIUM_CHANNELS | 2 | Down | 0.008487 | 0.0805817 |
| REACTOME_PHASE_4_RESTING_MEMBRANE_POTENTIAL | 2 | Down | 0.008487 | 0.0805817 |
| SHETH_LIVER_CANCER_VS_TXNIP_LOSS_PAM3 | 20 | Up | 0.00853 | 0.0808269 |
| PID_PRL_SIGNALING_EVENTS_PATHWAY | 1 | Up | 0.008709 | 0.0820363 |
| PID_IL2_STAT5_PATHWAY | 1 | Up | 0.008709 | 0.0820363 |
| IWANAGA_E2F1_TARGETS_NOT_INDUCED_BY_SERUM | 1 | Up | 0.008709 | 0.0820363 |
| REACTOME_DEPOSITION_OF_NEW_CENPA_CONTAINING_NUCLEOSOMES_AT_THE_CENTROMERE | 11 | Up | 0.008752 | 0.0822747 |
| KEGG_NUCLEOTIDE_EXCISION_REPAIR | 11 | Up | 0.008949 | 0.0839658 |
| REACTOME_GPCR_LIGAND_BINDING | 49 | Down | 0.009044 | 0.0846815 |
| YAO_HOXA10_TARGETS_VIA_PROGESTERONE_UP | 16 | Down | 0.009119 | 0.0851126 |
| REACTOME_REGULATION_OF_MECP2_EXPRESSION_AND_ACTIVITY | 3 | Up | 0.009126 | 0.0851126 |
| MATZUK_SPERMATOCYTE | 11 | Up | 0.009243 | 0.0860398 |
| IVANOVA_HEMATOPOIESIS_STEM_CELL | 29 | Down | 0.009268 | 0.0861053 |
| PIONTEK_PKD1_TARGETS_DN | 5 | Up | 0.009436 | 0.0874625 |
| ELVIDGE_HIF1A_AND_HIF2A_TARGETS_DN | 21 | Down | 0.009451 | 0.0874625 |
| REACTOME_RECEPTOR_TYPE_TYROSINE_PROTEIN_PHOSPHATASES | 10 | Down | 0.009525 | 0.087973 |
| REACTOME_CHL1_INTERACTIONS | 1 | Down | 0.009591 | 0.0884131 |
| MARSON_BOUND_BY_FOXP3_UNSTIMULATED | 108 | Up | 0.009667 | 0.0889359 |
| SWEET_LUNG_CANCER_KRAS_DN | 91 | Down | 0.009687 | 0.0889496 |
| KANG_IMMORTALIZED_BY_TERT_UP | 17 | Down | 0.009715 | 0.0890368 |
| HELLEBREKERS_SILENCED_DURING_TUMOR_ANGIOGENESIS | 14 | Down | 0.009884 | 0.0904083 |
| REACTOME_SUMOYLATION_OF_DNA_DAMAGE_RESPONSE_AND_REPAIR_PROTEINS | 5 | Up | 0.010172 | 0.0928607 |
| MIKKELSEN_MEF_ICP_WITH_H3K27ME3 | 16 | Down | 0.010233 | 0.0932397 |
| IRITANI_MAD1_TARGETS_DN | 4 | Up | 0.010571 | 0.0960875 |
| KATSANOU_ELAVL1_TARGETS_DN | 36 | Up | 0.0106 | 0.0960875 |
| REACTOME_FANCONI_ANEMIA_PATHWAY | 8 | Up | 0.010607 | 0.0960875 |
| MARIADASON_RESPONSE_TO_BUTYRATE_SULINDAC_6 | 5 | Up | 0.010724 | 0.0969662 |
| YANG_BREAST_CANCER_ESR1_BULK_DN | 3 | Up | 0.010757 | 0.0970581 |
| OLSSON_E2F3_TARGETS_DN | 10 | Up | 0.010775 | 0.0970581 |
| BIOCARTA_NO1_PATHWAY | 2 | Down | 0.010847 | 0.0975175 |
| BENPORATH_SUZ12_TARGETS | 185 | Down | 0.010901 | 0.0978205 |
| GINESTIER_BREAST_CANCER_20Q13_AMPLIFICATION_DN | 16 | Up | 0.011041 | 0.0988877 |
| ABBUD_LIF_SIGNALING_2_DN | 1 | Down | 0.011117 | 0.0993682 |
| GENTILE_UV_RESPONSE_CLUSTER_D4 | 8 | Up | 0.011136 | 0.0993682 |
| YUAN_ZNF143_PARTNERS | 4 | Up | 0.011166 | 0.0994436 |

| | | | | |
|---|---|---|---|---|
| REACTOME_NUCLEOTIDE_EXCISION_REPAIR | 15 | Up | 0.011257 | 0.1000636 |
| REACTOME_GLOBAL_GENOME_NUCLEOTIDE_EXCISION_REPAIR_GG_NER | 12 | Up | 0.011417 | 0.1012968 |
| REACTOME_MEMBRANE_TRAFFICKING | 56 | Up | 0.011642 | 0.1031041 |
| HOLLEMAN_VINCRISTINE_RESISTANCE_B_ALL_UP | 5 | Down | 0.011712 | 0.1035286 |
| REACTOME_CA2_ACTIVATED_K_CHANNELS | 2 | Down | 0.011779 | 0.103931 |
| REACTOME_FATTY_ACYL_COA_BIOSYNTHESIS | 10 | Down | 0.011926 | 0.1050319 |
| HOOI_ST7_TARGETS_DN | 22 | Down | 0.012055 | 0.1059673 |
| MIKKELSEN_NPC_HCP_WITH_H3K4ME3_AND_H3K27ME3 | 37 | Down | 0.012342 | 0.1079864 |
| BORLAK_LIVER_CANCER_EGF_UP | 10 | Up | 0.012353 | 0.1079864 |
| BIOCARTA_CARM_ER_PATHWAY | 1 | Up | 0.012401 | 0.1079864 |
| XU_AKT1_TARGETS_48HR | 1 | Up | 0.012401 | 0.1079864 |
| OHM_EMBRYONIC_CARCINOMA_UP | 1 | Up | 0.012401 | 0.1079864 |
| ZHAN_MULTIPLE_MYELOMA_MF_DN | 9 | Down | 0.012421 | 0.1079864 |
| SANSOM_APC_TARGETS_REQUIRE_MYC | 23 | Up | 0.012528 | 0.1087132 |
| BROWNE_HCMV_INFECTION_20HR_UP | 24 | Down | 0.012588 | 0.1090304 |
| REACTOME_VEGFR2_MEDIATED_CELL_PROLIFERATION | 2 | Down | 0.012646 | 0.1093404 |
| REACTOME_DISEASES_OF_BASE_EXCISION_REPAIR | 1 | Up | 0.012743 | 0.1096464 |
| IVANOVA_HEMATOPOIESIS_LATE_PROGENITOR | 55 | Up | 0.012749 | 0.1096464 |
| NUYTTEN_EZH2_TARGETS_UP | 122 | Down | 0.012751 | 0.1096464 |
| BUYTAERT_PHOTODYNAMIC_THERAPY_STRESS_DN | 120 | Up | 0.012778 | 0.1096785 |
| REACTOME_UB_SPECIFIC_PROCESSING_PROTEASES | 13 | Up | 0.013067 | 0.1118487 |
| REACTOME_TRANSCRIPTION_COUPLED_NUCLEOTIDE_EXCISION_REPAIR_TC_NER | 14 | Up | 0.013078 | 0.1118487 |
| REACTOME_RESOLUTION_OF_ABASIC_SITES_AP_SITES | 9 | Up | 0.013121 | 0.1120173 |
| GHO_ATF5_TARGETS_DN | 2 | Up | 0.013405 | 0.1141535 |
| PARK_HSC_AND_MULTIPOTENT_PROGENITORS | 5 | Up | 0.01342 | 0.1141535 |
| REACTOME_PROCESSIVE_SYNTHESIS_ON_THE_C_STRAND_OF_THE_TELOMERE | 4 | Up | 0.013569 | 0.1152172 |
| DURAND_STROMA_NS_UP | 30 | Down | 0.013752 | 0.1165587 |
| ELLWOOD_MYC_TARGETS_DN | 6 | Down | 0.014006 | 0.1184993 |
| TARTE_PLASMA_CELL_VS_PLASMABLAST_UP | 49 | Down | 0.014073 | 0.1188518 |
| YAO_TEMPORAL_RESPONSE_TO_PROGESTERONE_CLUSTER_16 | 20 | Down | 0.014391 | 0.1213234 |
| REACTOME_DNA_REPLICATION_INITIATION | 4 | Up | 0.014577 | 0.1225299 |
| HEIDENBLAD_AMPLICON_8Q24_UP | 6 | Up | 0.014586 | 0.1225299 |
| ELVIDGE_HYPOXIA_UP | 32 | Down | 0.014677 | 0.122943 |
| ANASTASSIOU_MULTICANCER_INVASIVENESS_SIGNATURE | 23 | Down | 0.014687 | 0.122943 |
| GOLDRATH_IMMUNE_MEMORY | 8 | Down | 0.014751 | 0.1230487 |
| ACEVEDO_LIVER_CANCER_WITH_H3K9ME3_DN | 17 | Up | 0.014751 | 0.1230487 |
| GARY_CD5_TARGETS_DN | 58 | Up | 0.014807 | 0.1232351 |
| ZHAN_MULTIPLE_MYELOMA_PR_DN | 9 | Down | 0.014826 | 0.1232351 |
| JAZAERI_BREAST_CANCER_BRCA1_VS_BRCA2_UP | 3 | Up | 0.014883 | 0.1234937 |
| KEGG_HOMOLOGOUS_RECOMBINATION | 11 | Up | 0.015053 | 0.1246881 |
| MCMURRAY_TP53_HRAS_COOPERATION_RESPONSE_UP | 4 | Down | 0.015306 | 0.1264861 |

| | | | | |
|---|---|---|---|---|
| REACTOME_E2F_ENABLED_INHIBITION_OF_PRE_REPLICATION_COMPLEX_FORMATION | 6 | Up | 0.015324 | 0.1264861 |
| REACTOME_EPHB_MEDIATED_FORWARD_SIGNALING | 5 | Down | 0.015476 | 0.1275012 |
| ZHANG_RESPONSE_TO_CANTHARIDIN_DN | 8 | Up | 0.0155 | 0.1275012 |
| GROSS_HYPOXIA_VIA_HIF1A_ONLY | 3 | Down | 0.015634 | 0.1283756 |
| GROSS_ELK3_TARGETS_UP | 7 | Down | 0.015664 | 0.1284028 |
| REACTOME_HYALURONAN_BIOSYNTHESIS_AND_EXPORT | 2 | Down | 0.015715 | 0.1284763 |
| CHIANG_LIVER_CANCER_SUBCLASS_UNANNOTATED_DN | 33 | Up | 0.015727 | 0.1284763 |
| SESTO_RESPONSE_TO_UV_C7 | 9 | Up | 0.015755 | 0.1284824 |
| PID_VEGFR1_2_PATHWAY | 6 | Down | 0.015883 | 0.1293027 |
| KAMMINGA_SENESCENCE | 3 | Up | 0.015985 | 0.1299082 |
| BENPORATH_ES_WITH_H3K27ME3 | 190 | Down | 0.016071 | 0.1301927 |
| REACTOME_DNA_DOUBLE_STRAND_BREAK_RESPONSE | 8 | Up | 0.016075 | 0.1301927 |
| REACTOME_TRANSMISSION_ACROSS_CHEMICAL_SYNAPSES | 31 | Down | 0.016109 | 0.1302463 |
| LAU_APOPTOSIS_CDKN2A_UP | 6 | Up | 0.016302 | 0.1315822 |
| BIOCARTA_FLUMAZENIL_PATHWAY | 1 | Down | 0.016516 | 0.1330824 |
| REACTOME_L1CAM_INTERACTIONS | 13 | Down | 0.016642 | 0.133867 |
| KIM_MYCN_AMPLIFICATION_TARGETS_DN | 19 | Down | 0.016773 | 0.1346969 |
| SCHAEFFER_PROSTATE_DEVELOPMENT_48HR_UP | 82 | Down | 0.016812 | 0.1347819 |
| CAFFAREL_RESPONSE_TO_THC_24HR_3_DN | 2 | Up | 0.016928 | 0.1353206 |
| REACTOME_SCF_SKP2_MEDIATED_DEGRADATION_OF_P27_P21 | 3 | Up | 0.016936 | 0.1353206 |
| LIN_NPAS4_TARGETS_UP | 15 | Down | 0.01712 | 0.1365594 |
| HADDAD_T_LYMPHOCYTE_AND_NK_PROGENITOR_DN | 13 | Up | 0.017214 | 0.1370756 |
| REACTOME_MAPK6_MAPK4_SIGNALING | 5 | Up | 0.017289 | 0.1374416 |
| LOPEZ_MESOTHELIOMA_SURVIVAL_WORST_VS_BEST_UP | 5 | Up | 0.01735 | 0.1376918 |
| SANSOM_APC_TARGETS_UP | 21 | Up | 0.017624 | 0.1396394 |
| BENPORATH_EED_TARGETS | 177 | Down | 0.017732 | 0.1402566 |
| RIZ_ERYTHROID_DIFFERENTIATION_12HR | 4 | Down | 0.017842 | 0.14089 |
| MOREAUX_B_LYMPHOCYTE_MATURATION_BY_TACI_DN | 14 | Up | 0.01791 | 0.1411951 |
| URS_ADIPOCYTE_DIFFERENTIATION_UP | 15 | Down | 0.017947 | 0.1412533 |
| BIOCARTA_PTC1_PATHWAY | 5 | Up | 0.018052 | 0.1418406 |
| MURAKAMI_UV_RESPONSE_1HR_UP | 2 | Down | 0.018082 | 0.1418416 |
| LUCAS_HNF4A_TARGETS_UP | 12 | Down | 0.018169 | 0.1422919 |
| LY_AGING_OLD_UP | 3 | Down | 0.018477 | 0.14446 |
| MINGUEZ_LIVER_CANCER_VASCULAR_INVASION_UP | 1 | Up | 0.01862 | 0.1448637 |
| REACTOME_SYNTHESIS_OF_ACTIVE_UBIQUITIN_ROLES_OF_E1_AND_E2_ENZYMES | 1 | Up | 0.01862 | 0.1448637 |
| IVANOVA_HEMATOPOIESIS_STEM_CELL_LONG_TERM | 34 | Down | 0.01862 | 0.1448637 |
| PID_EPHB_FWD_PATHWAY | 2 | Down | 0.018793 | 0.1459716 |
| REACTOME_CILIUM_ASSEMBLY | 33 | Up | 0.018875 | 0.1461709 |
| REACTOME_DUAL_INCISION_IN_GG_NER | 9 | Up | 0.018881 | 0.1461709 |
| UROSEVIC_RESPONSE_TO_IMIQUIMOD | 3 | Up | 0.018914 | 0.1461895 |
| POMEROY_MEDULLOBLASTOMA_DESMOPLASIC_VS_CLASSIC_DN | 13 | Down | 0.018995 | 0.1465764 |
| CHIANG_LIVER_CANCER_SUBCLASS_POLYSOMY7_UP | 14 | Down | 0.019272 | 0.1484696 |

| | | | | |
|---|---|---|---|---|
| REACTOME_TRAFFICKING_OF_GLUR2_CONTAINING_AMPA_RECEPTORS | 4 | Down | 0.019311 | 0.1485353 |
| REACTOME_GAP_FILLING_DNA_REPAIR_SYNTHESIS_AND_LIGATION_IN_GG_NER | 7 | Up | 0.019382 | 0.1488369 |
| MANALO_HYPOXIA_UP | 31 | Down | 0.019448 | 0.1491024 |
| REACTOME_NEUROFASCIN_INTERACTIONS | 2 | Down | 0.019527 | 0.1494237 |
| REACTOME_POLYMERASE_SWITCHING_ON_THE_C_STRAND_OF_THE_TELOMERE | 6 | Up | 0.019553 | 0.1494237 |
| PARK_HSC_MARKERS | 3 | Up | 0.019599 | 0.1495313 |
| BIOCARTA_TUBBY_PATHWAY | 1 | Down | 0.019744 | 0.149912 |
| BIOCARTA_BOTULIN_PATHWAY | 1 | Down | 0.019744 | 0.149912 |
| REACTOME_MUSCARINIC_ACETYLCHOLINE_RECEPTORS | 1 | Down | 0.019744 | 0.149912 |
| PID_VEGFR1_PATHWAY | 2 | Down | 0.019813 | 0.1501969 |
| CHUNG_BLISTER_CYTOTOXICITY_DN | 3 | Down | 0.020027 | 0.1515807 |
| KEGG_GLYCEROPHOSPHOLIPID_METABOLISM | 10 | Down | 0.020143 | 0.1519952 |
| REACTOME_PLATELET_ACTIVATION_SIGNALING_AND_AGGREGATION | 29 | Down | 0.020146 | 0.1519952 |
| ZHAN_EARLY_DIFFERENTIATION_GENES_DN | 8 | Up | 0.020257 | 0.1525884 |
| REACTOME_INACTIVATION_OF_CDC42_AND_RAC1 | 1 | Down | 0.020676 | 0.1554992 |
| JOSEPH_RESPONSE_TO_SODIUM_BUTYRATE_DN | 9 | Down | 0.020847 | 0.1565347 |
| YAMAZAKI_TCEB3_TARGETS_UP | 36 | Down | 0.020936 | 0.156955 |
| CROONQUIST_NRAS_VS_STROMAL_STIMULATION_UP | 4 | Down | 0.021159 | 0.1583762 |
| KEGG_SMALL_CELL_LUNG_CANCER | 15 | Down | 0.021208 | 0.1584895 |
| REACTOME_METABOLISM_OF_LIPIDS | 86 | Down | 0.021479 | 0.1601487 |
| HAN_SATB1_TARGETS_UP | 68 | Down | 0.021497 | 0.1601487 |
| KIM_MYCL1_AMPLIFICATION_TARGETS_DN | 3 | Up | 0.021583 | 0.1604611 |
| REACTOME_SIGNALING_BY_RETINOIC_ACID | 6 | Down | 0.021607 | 0.1604611 |
| REACTOME_SEROTONIN_RECEPTORS | 2 | Down | 0.021648 | 0.1605145 |
| REACTOME_DUAL_INCISION_IN_TC_NER | 12 | Up | 0.022374 | 0.1656359 |
| WANG_TUMOR_INVASIVENESS_UP | 37 | Up | 0.022566 | 0.1668017 |
| GUO_HEX_TARGETS_UP | 9 | Down | 0.022706 | 0.1675736 |
| REACTOME_ION_HOMEOSTASIS | 4 | Down | 0.022774 | 0.1678128 |
| KEGG_CELL_ADHESION_MOLECULES_CAMS | 19 | Down | 0.022833 | 0.1679889 |
| BIOCARTA_SRCRPTP_PATHWAY | 4 | Up | 0.023167 | 0.1700064 |
| MCDOWELL_ACUTE_LUNG_INJURY_DN | 11 | Down | 0.023179 | 0.1700064 |
| REACTOME_TRANSLESION_SYNTHESIS_BY_Y_FAMILY_DNA_POLYMERASES_BYPASSES_LESIONS_ON_DNA_TEMPLATE | 6 | Up | 0.023408 | 0.1711531 |
| REACTOME_TERMINATION_OF_TRANSLESION_DNA_SYNTHESIS | 6 | Up | 0.023408 | 0.1711531 |
| KEGG_FOCAL_ADHESION | 40 | Down | 0.023547 | 0.1719012 |
| BERTUCCI_MEDULLARY_VS_DUCTAL_BREAST_CANCER_DN | 34 | Down | 0.023907 | 0.1740239 |
| GENTILE_UV_RESPONSE_CLUSTER_D1 | 1 | Up | 0.023911 | 0.1740239 |
| XU_HGF_SIGNALING_NOT_VIA_AKT1_48HR_DN | 7 | Up | 0.024153 | 0.1755191 |
| REACTOME_ACTIVATION_OF_GENE_EXPRESSION_BY_SREBF_SREBP | 6 | Down | 0.024228 | 0.175795 |

| | | | | |
|---|---|---|---|---|
| REACTOME_NOTCH3_ACTIVATION_AND_TRANSMISSION_OF_SIGNAL_TO_THE_NUCLEUS | 4 | Up | 0.024298 | 0.1760006 |
| WEINMANN_ADAPTATION_TO_HYPOXIA_DN | 13 | Down | 0.024332 | 0.1760006 |
| BIOCARTA_GABA_PATHWAY | 2 | Down | 0.024368 | 0.1760006 |
| XU_GH1_AUTOCRINE_TARGETS_DN | 18 | Down | 0.02446 | 0.1763959 |
| LASTOWSKA_NEUROBLASTOMA_COPY_NUMBER_UP | 21 | Up | 0.024749 | 0.1779564 |
| BURTON_ADIPOGENESIS_10 | 4 | Down | 0.024752 | 0.1779564 |
| REACTOME_ACTIVATED_NTRK2_SIGNALS_THROUGH_RAS | 1 | Down | 0.025327 | 0.1809952 |
| REACTOME_ACTIVATED_NTRK2_SIGNALS_THROUGH_PI3K | 1 | Down | 0.025327 | 0.1809952 |
| REACTOME_ACTIVATED_NTRK2_SIGNALS_THROUGH_FRS2_AND_FRS3 | 1 | Down | 0.025327 | 0.1809952 |
| REACTOME_ACTIVATED_NTRK2_SIGNALS_THROUGH_CDK5 | 1 | Down | 0.025327 | 0.1809952 |
| REACTOME_LAMININ_INTERACTIONS | 15 | Down | 0.02543 | 0.1814574 |
| MOHANKUMAR_HOXA1_TARGETS_DN | 24 | Down | 0.025596 | 0.1823696 |
| MCBRYAN_TERMINAL_END_BUD_UP | 3 | Up | 0.025664 | 0.1825771 |
| KYNG_RESPONSE_TO_H2O2_VIA_ERCC6_DN | 3 | Down | 0.026112 | 0.1854821 |
| CHUNG_BLISTER_CYTOTOXICITY_UP | 13 | Up | 0.026606 | 0.1881173 |
| SANSOM_APC_TARGETS | 33 | Up | 0.026624 | 0.1881173 |
| RASHI_RESPONSE_TO_IONIZING_RADIATION_4 | 7 | Up | 0.026636 | 0.1881173 |
| PID_INTEGRIN3_PATHWAY | 12 | Down | 0.026641 | 0.1881173 |
| PEREZ_TP53_TARGETS | 163 | Down | 0.026765 | 0.1887063 |
| CHIBA_RESPONSE_TO_TSA_DN | 6 | Down | 0.027026 | 0.1900549 |
| REACTOME_TRAFFICKING_OF_AMPA_RECEPTORS | 6 | Down | 0.027036 | 0.1900549 |
| GAVIN_FOXP3_TARGETS_CLUSTER_T7 | 6 | Up | 0.027088 | 0.1901387 |
| SHETH_LIVER_CANCER_VS_TXNIP_LOSS_PAM1 | 38 | Up | 0.027169 | 0.1902883 |
| WAMUNYOKOLI_OVARIAN_CANCER_LMP_DN | 33 | Down | 0.027193 | 0.1902883 |
| BOYLAN_MULTIPLE_MYELOMA_D_UP | 8 | Down | 0.027242 | 0.1902883 |
| KEGG_LONG_TERM_DEPRESSION | 6 | Down | 0.027359 | 0.1902883 |
| REACTOME_GLI_PROTEINS_BIND_PROMOTERS_OF_HH_RESPONSIVE_GENES_TO_PROMOTE_TRANSCRIPTION | 1 | Down | 0.027382 | 0.1902883 |
| POMEROY_MEDULLOBLASTOMA_PROGNOSIS_UP | 7 | Down | 0.027393 | 0.1902883 |
| IKEDA_MIR30_TARGETS_DN | 7 | Down | 0.027402 | 0.1902883 |
| REACTOME_TP53_REGULATES_METABOLIC_GENES | 2 | Down | 0.027431 | 0.1902883 |
| REACTOME_REGULATION_OF_CHOLESTEROL_BIOSYNTHESIS_BY_SREBP_SREBF | 7 | Down | 0.027838 | 0.1925658 |
| NAKAMURA_METASTASIS | 6 | Down | 0.02784 | 0.1925658 |
| NIKOLSKY_BREAST_CANCER_17Q11_Q21_AMPLICON | 11 | Up | 0.02791 | 0.1927204 |
| WAKASUGI_HAVE_ZNF143_BINDING_SITES | 13 | Up | 0.027974 | 0.1927204 |
| KEGG_NON_SMALL_CELL_LUNG_CANCER | 3 | Down | 0.027999 | 0.1927204 |
| NUYTTEN_NIPP1_TARGETS_UP | 100 | Down | 0.028025 | 0.1927204 |
| KIM_ALL_DISORDERS_DURATION_CORR_DN | 6 | Down | 0.028086 | 0.1928564 |
| CHEN_LVAD_SUPPORT_OF_FAILING_HEART_UP | 13 | Down | 0.028317 | 0.1941602 |
| SHETH_LIVER_CANCER_VS_TXNIP_LOSS_PAM2 | 35 | Up | 0.028378 | 0.194299 |
| NAKAMURA_TUMOR_ZONE_PERIPHERAL_VS_CENTRAL_UP | 62 | Up | 0.028907 | 0.197637 |
| FARMER_BREAST_CANCER_BASAL_VS_LULMINAL | 53 | Up | 0.029072 | 0.1983066 |

| | | | | |
|---|---|---|---|---|
| SA_G2_AND_M_PHASES | 2 | Up | 0.029089 | 0.1983066 |
| DELLA_RESPONSE_TO_TSA_AND_BUTYRATE | 2 | Up | 0.030495 | 0.2074821 |
| SCHUETZ_BREAST_CANCER_DUCTAL_INVASIVE_UP | 80 | Down | 0.030522 | 0.2074821 |
| REACTOME_PEPTIDE_LIGAND_BINDING_RECEPTORS | 16 | Down | 0.030656 | 0.2080917 |
| REACTOME_RESPONSE_TO_ELEVATED_PLATELET_CYTOSOLIC_CA2 | 15 | Down | 0.030765 | 0.2085338 |
| DACOSTA_UV_RESPONSE_VIA_ERCC3_UP | 23 | Up | 0.031235 | 0.2114157 |
| BIOCARTA_ERBB4_PATHWAY | 2 | Down | 0.031427 | 0.2124135 |
| APPIERTO_RESPONSE_TO_FENRETINIDE_UP | 5 | Down | 0.031551 | 0.2129442 |
| NIKOLSKY_BREAST_CANCER_16P13_AMPLICON | 6 | Up | 0.031992 | 0.2156171 |
| UEDA_PERIFERAL_CLOCK | 19 | Down | 0.032101 | 0.2160467 |
| REACTOME_EXTRA_NUCLEAR_ESTROGEN_SIGNALING | 8 | Down | 0.032159 | 0.216127 |
| REACTOME_METALLOPROTEASE_DUBS | 2 | Up | 0.032746 | 0.2197626 |
| LINDVALL_IMMORTALIZED_BY_TERT_UP | 14 | Down | 0.032793 | 0.2197636 |
| ABE_VEGFA_TARGETS_2HR | 2 | Up | 0.032883 | 0.2200567 |
| RADAEVA_RESPONSE_TO_IFNA1_UP | 4 | Up | 0.033157 | 0.2214103 |
| DIAZ_CHRONIC_MEYLOGENOUS_LEUKEMIA_DN | 13 | Down | 0.033179 | 0.2214103 |
| HOFFMANN_PRE_BI_TO_LARGE_PRE_BII_LYMPHOCYTE_UP | 1 | Up | 0.033301 | 0.2219106 |
| WESTON_VEGFA_TARGETS_6HR | 13 | Down | 0.033393 | 0.2222138 |
| MCCLUNG_DELTA_FOSB_TARGETS_8WK | 5 | Down | 0.033656 | 0.223647 |
| NUYTTEN_NIPP1_TARGETS_DN | 129 | Up | 0.034019 | 0.2254546 |
| JIANG_VHL_TARGETS | 9 | Up | 0.034023 | 0.2254546 |
| BROWNE_INTERFERON_RESPONSIVE_GENES | 6 | Up | 0.034273 | 0.2263763 |
| JOSEPH_RESPONSE_TO_SODIUM_BUTYRATE_UP | 3 | Down | 0.034299 | 0.2263763 |
| KEGG_PPAR_SIGNALING_PATHWAY | 12 | Down | 0.034345 | 0.2263763 |
| KORKOLA_CORRELATED_WITH_POU5F1 | 4 | Up | 0.034353 | 0.2263763 |
| REN_ALVEOLAR_RHABDOMYOSARCOMA_DN | 86 | Down | 0.034408 | 0.2264252 |
| REACTOME_CELLULAR_RESPONSE_TO_HEAT_STRESS | 9 | Down | 0.034571 | 0.2268917 |
| WEST_ADRENOCORTICAL_TUMOR_DN | 90 | Down | 0.034575 | 0.2268917 |
| REACTOME_DEPOLYMERISATION_OF_THE_NUCLEAR_LAMINA | 5 | Up | 0.034656 | 0.2271074 |
| WANG_CLASSIC_ADIPOGENIC_TARGETS_OF_PPARG | 6 | Down | 0.034711 | 0.2271579 |
| REACTOME_REGULATION_OF_LIPID_METABOLISM_BY_PPARALPHA | 8 | Down | 0.035374 | 0.2311765 |
| BIOCARTA_MET_PATHWAY | 1 | Up | 0.035501 | 0.2316868 |
| KENNY_CTNNB1_TARGETS_UP | 11 | Up | 0.035601 | 0.2320167 |
| KEGG_MISMATCH_REPAIR | 6 | Up | 0.035685 | 0.2322463 |
| BIOCARTA_G1_PATHWAY | 6 | Up | 0.035931 | 0.2335284 |
| FEVR_CTNNB1_TARGETS_UP | 72 | Down | 0.036074 | 0.2341339 |
| REACTOME_SIGNALING_BY_NUCLEAR_RECEPTORS | 24 | Down | 0.036274 | 0.2344413 |
| REACTOME_MEIOSIS | 14 | Up | 0.036309 | 0.2344413 |
| REACTOME_REPRODUCTION | 14 | Up | 0.036309 | 0.2344413 |
| DANG_MYC_TARGETS_UP | 9 | Up | 0.036319 | 0.2344413 |
| MATHEW_FANCONI_ANEMIA_GENES | 3 | Up | 0.03647 | 0.2349467 |
| BILD_CTNNB1_ONCOGENIC_SIGNATURE | 13 | Down | 0.036497 | 0.2349467 |
| KONG_E2F1_TARGETS | 1 | Up | 0.036856 | 0.2367382 |

| | | | | |
|---|---|---|---|---|
| VERRECCHIA_EARLY_RESPONSE_TO_TGFB1 | 16 | Down | 0.036909 | 0.2367382 |
| AIGNER_ZEB1_TARGETS | 5 | Down | 0.036925 | 0.2367382 |
| REACTOME_MISMATCH_REPAIR | 5 | Up | 0.037251 | 0.2385087 |
| SCHWAB_TARGETS_OF_BMYB_POLYMORPHIC_VARIANTS_UP | 2 | Up | 0.037338 | 0.2387393 |
| REACTOME_SIGNALING_BY_VEGF | 10 | Down | 0.037588 | 0.2400186 |
| CHICAS_RB1_TARGETS_SENESCENT | 115 | Up | 0.037764 | 0.2408155 |
| GAUSSMANN_MLL_AF4_FUSION_TARGETS_C_UP | 18 | Down | 0.037819 | 0.2408389 |
| BERTUCCI_INVASIVE_CARCINOMA_DUCTAL_VS_LOBULAR_DN | 9 | Down | 0.037973 | 0.241495 |
| TERAMOTO_OPN_TARGETS_CLUSTER_8 | 1 | Down | 0.038117 | 0.2417625 |
| REACTOME_NRCAM_INTERACTIONS | 1 | Down | 0.038117 | 0.2417625 |
| WANG_THOC1_TARGETS_UP | 2 | Up | 0.03824 | 0.2422181 |
| HERNANDEZ_MITOTIC_ARREST_BY_DOCETAXEL_1_DN | 16 | Up | 0.038451 | 0.2432294 |
| REACTOME_NON_INTEGRIN_MEMBRANE_ECM_INTERACTIONS | 27 | Down | 0.03864 | 0.2441043 |
| REACTOME_SYNAPTIC_ADHESION_LIKE_MOLECULES | 3 | Down | 0.039024 | 0.2461967 |
| MATSUDA_NATURAL_KILLER_DIFFERENTIATION | 65 | Up | 0.039636 | 0.2497281 |
| WEIGEL_OXIDATIVE_STRESS_BY_HNE_AND_H2O2 | 5 | Up | 0.039864 | 0.2508333 |
| WANG_TUMOR_INVASIVENESS_DN | 22 | Up | 0.039941 | 0.2509679 |
| REACTOME_GLYCEROPHOSPHOLIPID_BIOSYNTHESIS | 15 | Down | 0.039992 | 0.2509679 |
| SCHAEFFER_PROSTATE_DEVELOPMENT_12HR_UP | 27 | Down | 0.040082 | 0.2512043 |
| DORN_ADENOVIRUS_INFECTION_48HR_UP | 5 | Up | 0.040469 | 0.2530453 |
| ZHAN_VARIABLE_EARLY_DIFFERENTIATION_GENES_DN | 3 | Up | 0.040483 | 0.2530453 |
| MATZUK_MALE_REPRODUCTION_SERTOLI | 8 | Up | 0.040656 | 0.253794 |
| BIOCARTA_BCELLSURVIVAL_PATHWAY | 1 | Up | 0.040861 | 0.2543999 |
| REACTOME_TP53_REGULATES_TRANSCRIPTION_OF_SEVERAL_ADDITIONAL_CELL_DEATH_GENES_WHOSE_SPECIFIC_ROLES_IN_P53_DEPENDENT_APOPTOSIS_REMAIN_UNCERTAIN | 1 | Up | 0.040861 | 0.2543999 |
| RODRIGUES_THYROID_CARCINOMA_ANAPLASTIC_DN | 91 | Down | 0.04116 | 0.2547014 |
| BIOCARTA_ATM_PATHWAY | 7 | Up | 0.041361 | 0.2547014 |
| BENPORATH_PRC2_TARGETS | 110 | Down | 0.041649 | 0.2547014 |
| MARCHINI_TRABECTEDIN_RESISTANCE_DN | 10 | Down | 0.041771 | 0.2547014 |
| FOURNIER_ACINAR_DEVELOPMENT_LATE_UP | 3 | Down | 0.041864 | 0.2547014 |
| WHITFIELD_CELL_CYCLE_M_G1 | 13 | Up | 0.041922 | 0.2547014 |
| DALESSIO_TSA_RESPONSE | 3 | Down | 0.042299 | 0.2547014 |
| REACTOME_TP53_REGULATES_TRANSCRIPTION_OF_DNA_REPAIR_GENES | 10 | Up | 0.042571 | 0.2547014 |
| BOYLAN_MULTIPLE_MYELOMA_C_UP | 7 | Up | 0.042897 | 0.2547014 |
| KEGG_LEISHMANIA_INFECTION | 1 | Down | 0.043181 | 0.2547014 |
| BIOCARTA_AT1R_PATHWAY | 1 | Down | 0.043181 | 0.2547014 |
| BIOCARTA_AGPCR_PATHWAY | 1 | Down | 0.043181 | 0.2547014 |
| BIOCARTA_BCR_PATHWAY | 1 | Down | 0.043181 | 0.2547014 |
| BIOCARTA_CDMAC_PATHWAY | 1 | Down | 0.043181 | 0.2547014 |
| BIOCARTA_CCR3_PATHWAY | 1 | Down | 0.043181 | 0.2547014 |
| BIOCARTA_CXCR4_PATHWAY | 1 | Down | 0.043181 | 0.2547014 |
| BIOCARTA_CALCINEURIN_PATHWAY | 1 | Down | 0.043181 | 0.2547014 |
| BIOCARTA_EGF_PATHWAY | 1 | Down | 0.043181 | 0.2547014 |

| | | | | |
|---|---|---|---|---|
| BIOCARTA_FCER1_PATHWAY | 1 | Down | 0.043181 | 0.2547014 |
| BIOCARTA_PYK2_PATHWAY | 1 | Down | 0.043181 | 0.2547014 |
| BIOCARTA_NOS1_PATHWAY | 1 | Down | 0.043181 | 0.2547014 |
| BIOCARTA_ARENRF2_PATHWAY | 1 | Down | 0.043181 | 0.2547014 |
| BIOCARTA_CCR5_PATHWAY | 1 | Down | 0.043181 | 0.2547014 |
| BIOCARTA_MYOSIN_PATHWAY | 1 | Down | 0.043181 | 0.2547014 |
| BIOCARTA_MEF2D_PATHWAY | 1 | Down | 0.043181 | 0.2547014 |
| BIOCARTA_GPCR_PATHWAY | 1 | Down | 0.043181 | 0.2547014 |
| BIOCARTA_TPO_PATHWAY | 1 | Down | 0.043181 | 0.2547014 |
| BIOCARTA_CREB_PATHWAY | 1 | Down | 0.043181 | 0.2547014 |
| BIOCARTA_VEGF_PATHWAY | 1 | Down | 0.043181 | 0.2547014 |
| PID_TCR_RAS_PATHWAY | 1 | Down | 0.043181 | 0.2547014 |
| PID_TCR_JNK_PATHWAY | 1 | Down | 0.043181 | 0.2547014 |
| PID_IL8_CXCR2_PATHWAY | 1 | Down | 0.043181 | 0.2547014 |
| PID_IL8_CXCR1_PATHWAY | 1 | Down | 0.043181 | 0.2547014 |
| REACTOME_DOWNSTREAM_SIGNALING_EVENTS_OF_B_CELL_RECEPTOR_BCR | 1 | Down | 0.043181 | 0.2547014 |
| SCHAEFFER_PROSTATE_DEVELOPMENT_AND_CANCER_BOX4_UP | 1 | Down | 0.043181 | 0.2547014 |
| BIOCARTA_PLC_PATHWAY | 1 | Down | 0.043181 | 0.2547014 |
| BIOCARTA_PKC_PATHWAY | 1 | Down | 0.043181 | 0.2547014 |
| BIOCARTA_ION_PATHWAY | 1 | Down | 0.043181 | 0.2547014 |
| BIOCARTA_PLCD_PATHWAY | 1 | Down | 0.043181 | 0.2547014 |
| REACTOME_DISINHIBITION_OF_SNARE_FORMATION | 1 | Down | 0.043181 | 0.2547014 |
| REACTOME_WNT5A_DEPENDENT_INTERNALIZATION_OF_FZD4 | 1 | Down | 0.043181 | 0.2547014 |
| REACTOME_NR1H2_AND_NR1H3_MEDIATED_SIGNALING | 3 | Down | 0.043186 | 0.2547014 |
| KIM_WT1_TARGETS_UP | 36 | Down | 0.043221 | 0.2547014 |
| TURASHVILI_BREAST_DUCTAL_CARCINOMA_VS_DUCTAL_NORMAL_DN | 35 | Down | 0.043361 | 0.2552085 |
| REACTOME_ORGANELLE_BIOGENESIS_AND_MAINTENANCE | 36 | Up | 0.04347 | 0.2555059 |
| GRANDVAUX_IRF3_TARGETS_DN | 4 | Down | 0.043519 | 0.2555059 |
| SCHAEFFER_SOX9_TARGETS_IN_PROSTATE_DEVELOPMENT_UP | 4 | Down | 0.043678 | 0.2555609 |
| BASSO_B_LYMPHOCYTE_NETWORK | 19 | Up | 0.04374 | 0.2555609 |
| REACTOME_PHOSPHOLIPID_METABOLISM | 24 | Down | 0.043828 | 0.2555609 |
| REACTOME_DNA_METHYLATION | 1 | Up | 0.043862 | 0.2555609 |
| BIOCARTA_SET_PATHWAY | 1 | Up | 0.04396 | 0.2555609 |
| REACTOME_APOPTOSIS_INDUCED_DNA_FRAGMENTATION | 1 | Up | 0.04396 | 0.2555609 |
| WU_HBX_TARGETS_1_UP | 1 | Up | 0.04396 | 0.2555609 |
| SARTIPY_NORMAL_AT_INSULIN_RESISTANCE_DN | 1 | Up | 0.04396 | 0.2555609 |
| REACTOME_MEIOTIC_RECOMBINATION | 9 | Up | 0.044511 | 0.2584469 |
| ZHENG_IL22_SIGNALING_DN | 11 | Up | 0.045305 | 0.2627381 |
| WALLACE_PROSTATE_CANCER_DN | 5 | Down | 0.045928 | 0.2659456 |
| PID_A6B1_A6B4_INTEGRIN_PATHWAY | 8 | Down | 0.045971 | 0.2659456 |
| REN_ALVEOLAR_RHABDOMYOSARCOMA_UP | 19 | Down | 0.046062 | 0.2661508 |
| KEGG_BIOSYNTHESIS_OF_UNSATURATED_FATTY_ACIDS | 8 | Down | 0.046564 | 0.2687205 |
| CHO_NR4A1_TARGETS | 1 | Down | 0.046621 | 0.2687205 |

| | | | | |
|---|---|---|---|---|
| PROVENZANI_METASTASIS_UP | 29 | Up | 0.046994 | 0.2705454 |
| REACTOME_NEUROTRANSMITTER_RELEASE_CYCLE | 7 | Down | 0.047076 | 0.2706441 |
| SEKI_INFLAMMATORY_RESPONSE_LPS_DN | 4 | Down | 0.047126 | 0.2706441 |
| REACTOME_NEUROTRANSMITTER_RECEPTORS_AND_POSTSYNAPTIC_SIGNAL_TRANSMISSION | 23 | Down | 0.04742 | 0.2710117 |
| DITTMER_PTHLH_TARGETS_UP | 15 | Up | 0.047505 | 0.2710117 |
| LEE_EARLY_T_LYMPHOCYTE_DN | 5 | Down | 0.047529 | 0.2710117 |
| MOREAUX_MULTIPLE_MYELOMA_BY_TACI_DN | 25 | Up | 0.047537 | 0.2710117 |
| KOKKINAKIS_METHIONINE_DEPRIVATION_48HR_DN | 12 | Up | 0.047575 | 0.2710117 |
| REACTOME_SIGNALING_BY_GPCR | 85 | Down | 0.047588 | 0.2710117 |
| REACTOME_SYNTHESIS_OF_VERY_LONG_CHAIN_FATTY_ACYL_COAS | 8 | Down | 0.04759 | 0.2710117 |
| NIKOLSKY_BREAST_CANCER_5P15_AMPLICON | 4 | Up | 0.047889 | 0.2723875 |
| MIYAGAWA_TARGETS_OF_EWSR1_ETS_FUSIONS_DN | 53 | Down | 0.048249 | 0.2741027 |
| BOYLAN_MULTIPLE_MYELOMA_C_CLUSTER_UP | 7 | Up | 0.048314 | 0.274147 |
| DORN_ADENOVIRUS_INFECTION_32HR_UP | 4 | Up | 0.048403 | 0.274321 |
| KEGG_ALDOSTERONE_REGULATED_SODIUM_REABSORPTION | 4 | Down | 0.04859 | 0.2750539 |
| BIOCARTA_ACE2_PATHWAY | 5 | Down | 0.048812 | 0.2755318 |
| LIN_MELANOMA_COPY_NUMBER_UP | 8 | Down | 0.048865 | 0.2755318 |
| REACTOME_RUNX2_REGULATES_CHONDROCYTE_MATURATION | 1 | Down | 0.049004 | 0.2755318 |
| REACTOME_RUNX2_REGULATES_GENES_INVOLVED_IN_CELL_MIGRATION | 1 | Down | 0.049004 | 0.2755318 |
| BIOCARTA_SPPA_PATHWAY | 2 | Down | 0.049023 | 0.2755318 |
| BIOCARTA_PAR1_PATHWAY | 2 | Down | 0.049023 | 0.2755318 |
| REACTOME_RHO_GTPASES_ACTIVATE_CIT | 5 | Up | 0.049157 | 0.2756275 |
| STARK_PREFRONTAL_CORTEX_22Q11_DELETION_UP | 12 | Down | 0.049157 | 0.2756275 |
| REACTOME_HYALURONAN_UPTAKE_AND_DEGRADATION | 1 | Up | 0.050013 | 0.2794603 |
