## Supplementary material for "p38γMAPK delays myelination and remyelination and is abundant in multiple sclerosis lesions": Sup Table 4

lrt_Mapk12ko vs. wt_topGO

| | Term | Ont | N | Up | Down | P.Up | P.Down |
|---|---|---|---|---|---|---|---|
| GO:0007049 | cell cycle | BP | 276 | 53 | 5 | 6.77E-33 | 7.92E-01 |
| GO:1903047 | mitotic cell cycle process | BP | 166 | 43 | 2 | 7.15E-31 | 9.10E-01 |
| GO:0022402 | cell cycle process | BP | 215 | 46 | 3 | 1.97E-29 | 8.93E-01 |
| GO:0000278 | mitotic cell cycle | BP | 191 | 43 | 2 | 4.78E-28 | 9.47E-01 |
| GO:0000280 | nuclear division | BP | 101 | 30 | 0 | 2.04E-22 | 1.00E+00 |
| GO:0000070 | mitotic sister chromatid segregation | BP | 55 | 24 | 0 | 2.04E-22 | 1.00E+00 |
| GO:0000819 | sister chromatid segregation | BP | 62 | 25 | 0 | 2.52E-22 | 1.00E+00 |
| GO:0051276 | chromosome organization | BP | 145 | 34 | 2 | 6.34E-22 | 8.64E-01 |
| GO:0007059 | chromosome segregation | BP | 88 | 28 | 0 | 9.24E-22 | 1.00E+00 |
| GO:0048285 | organelle fission | BP | 106 | 30 | 1 | 1.01E-21 | 9.23E-01 |
| GO:0051301 | cell division | BP | 130 | 32 | 1 | 3.30E-21 | 9.58E-01 |
| GO:0051726 | regulation of cell cycle | BP | 168 | 35 | 3 | 8.77E-21 | 7.65E-01 |
| GO:0140014 | mitotic nuclear division | BP | 78 | 26 | 0 | 9.97E-21 | 1.00E+00 |
| GO:0098813 | nuclear chromosome segregation | BP | 74 | 25 | 0 | 4.55E-20 | 1.00E+00 |
| GO:0010564 | regulation of cell cycle process | BP | 131 | 31 | 1 | 6.33E-20 | 9.59E-01 |
| GO:0007051 | spindle organization | BP | 48 | 18 | 0 | 2.20E-15 | 1.00E+00 |
| GO:0090068 | positive regulation of cell cycle process | BP | 55 | 18 | 0 | 3.61E-14 | 1.00E+00 |
| GO:0000226 | microtubule cytoskeleton organization | BP | 102 | 23 | 2 | 4.43E-14 | 6.98E-01 |
| GO:0007017 | microtubule-based process | BP | 125 | 25 | 2 | 5.16E-14 | 8.01E-01 |
| GO:0006996 | organelle organization | BP | 477 | 46 | 9 | 6.34E-14 | 8.16E-01 |
| GO:0044772 | mitotic cell cycle phase transition | BP | 84 | 21 | 1 | 7.92E-14 | 8.68E-01 |
| GO:0045787 | positive regulation of cell cycle | BP | 63 | 18 | 0 | 5.28E-13 | 1.00E+00 |
| GO:1902850 | microtubule cytoskeleton organization involved in mitosis | BP | 41 | 15 | 0 | 9.96E-13 | 1.00E+00 |
| GO:0051321 | meiotic cell cycle | BP | 57 | 17 | 0 | 1.19E-12 | 1.00E+00 |
| GO:0044770 | cell cycle phase transition | BP | 96 | 21 | 1 | 1.40E-12 | 9.02E-01 |
| GO:0007052 | mitotic spindle organization | BP | 35 | 14 | 0 | 1.46E-12 | 1.00E+00 |
| GO:0050000 | chromosome localization | BP | 21 | 11 | 0 | 1.17E-11 | 1.00E+00 |
| GO:0051303 | establishment of chromosome localization | BP | 21 | 11 | 0 | 1.17E-11 | 1.00E+00 |
| GO:0051225 | spindle assembly | BP | 29 | 12 | 0 | 4.26E-11 | 1.00E+00 |
| GO:0051983 | regulation of chromosome segregation | BP | 36 | 13 | 0 | 4.74E-11 | 1.00E+00 |
| GO:0006259 | DNA metabolic process | BP | 154 | 24 | 2 | 6.02E-11 | 8.86E-01 |
| GO:0008608 | attachment of spindle microtubules to kinetochore | BP | 14 | 9 | 0 | 8.22E-11 | 1.00E+00 |
| GO:0051310 | metaphase plate congression | BP | 19 | 10 | 0 | 1.04E-10 | 1.00E+00 |
| GO:1903046 | meiotic cell cycle process | BP | 47 | 14 | 0 | 1.55E-10 | 1.00E+00 |
| GO:1901990 | regulation of mitotic cell cycle phase transition | BP | 65 | 16 | 1 | 1.56E-10 | 7.89E-01 |
| GO:0007346 | regulation of mitotic cell cycle | BP | 97 | 19 | 1 | 1.65E-10 | 9.04E-01 |
| GO:0007080 | mitotic metaphase plate congression | BP | 15 | 9 | 0 | 2.00E-10 | 1.00E+00 |
| GO:0033044 | regulation of chromosome organization | BP | 49 | 14 | 0 | 2.91E-10 | 1.00E+00 |
| GO:0051304 | chromosome separation | BP | 35 | 12 | 0 | 5.80E-10 | 1.00E+00 |
| GO:0140694 | non-membrane-bounded organelle assembly | BP | 63 | 15 | 0 | 1.06E-09 | 1.00E+00 |
| GO:0006974 | cellular response to DNA damage stimulus | BP | 124 | 20 | 1 | 1.96E-09 | 9.51E-01 |
| GO:1905818 | regulation of chromosome separation | BP | 31 | 11 | 0 | 2.14E-09 | 1.00E+00 |
| GO:1901987 | regulation of cell cycle phase transition | BP | 77 | 16 | 1 | 2.39E-09 | 8.43E-01 |
| GO:0140013 | meiotic nuclear division | BP | 40 | 12 | 0 | 3.38E-09 | 1.00E+00 |
| GO:0051656 | establishment of organelle localization | BP | 49 | 13 | 0 | 3.77E-09 | 1.00E+00 |
| GO:0030261 | chromosome condensation | BP | 11 | 7 | 0 | 1.41E-08 | 1.00E+00 |
| GO:0044839 | cell cycle G2/M phase transition | BP | 37 | 11 | 1 | 1.83E-08 | 5.85E-01 |
| GO:0051988 | regulation of attachment of spindle microtubules to kinetochore | BP | 8 | 6 | 0 | 3.94E-08 | 1.00E+00 |
| GO:0007093 | mitotic cell cycle checkpoint signaling | BP | 40 | 11 | 0 | 4.54E-08 | 1.00E+00 |
| GO:0007010 | cytoskeleton organization | BP | 217 | 24 | 5 | 8.48E-08 | 5.82E-01 |
| GO:0000075 | cell cycle checkpoint signaling | BP | 43 | 11 | 0 | 1.04E-07 | 1.00E+00 |
| GO:0033043 | regulation of organelle organization | BP | 174 | 21 | 0 | 1.56E-07 | 1.00E+00 |
| GO:0006281 | DNA repair | BP | 102 | 16 | 1 | 1.69E-07 | 9.15E-01 |
| GO:0051640 | organelle localization | BP | 67 | 13 | 3 | 2.22E-07 | 2.03E-01 |
| GO:0061640 | cytoskeleton-dependent cytokinesis | BP | 15 | 7 | 1 | 2.46E-07 | 2.99E-01 |
| GO:0016043 | cellular component organization | BP | 796 | 49 | 24 | 2.66E-07 | 7.04E-02 |
| GO:0051315 | attachment of mitotic spindle microtubules to kinetochore | BP | 6 | 5 | 0 | 2.66E-07 | 1.00E+00 |
| GO:0045786 | negative regulation of cell cycle | BP | 68 | 13 | 1 | 2.67E-07 | 8.04E-01 |
| GO:1901992 | positive regulation of mitotic cell cycle phase transition | BP | 22 | 8 | 0 | 3.27E-07 | 1.00E+00 |
| GO:0051783 | regulation of nuclear division | BP | 39 | 10 | 0 | 4.10E-07 | 1.00E+00 |
| GO:0071840 | cellular component organization or biogenesis | BP | 808 | 49 | 24 | 4.57E-07 | 8.26E-02 |
| GO:0045931 | positive regulation of mitotic cell cycle | BP | 23 | 8 | 0 | 4.87E-07 | 1.00E+00 |
| GO:1901991 | negative regulation of mitotic cell cycle phase transition | BP | 40 | 10 | 0 | 5.32E-07 | 1.00E+00 |
| GO:0090307 | mitotic spindle assembly | BP | 17 | 7 | 0 | 7.02E-07 | 1.00E+00 |
| GO:0044784 | metaphase/anaphase transition of cell cycle | BP | 24 | 8 | 0 | 7.10E-07 | 1.00E+00 |
| GO:0007091 | metaphase/anaphase transition of mitotic cell cycle | BP | 24 | 8 | 0 | 7.10E-07 | 1.00E+00 |
| GO:0010948 | negative regulation of cell cycle process | BP | 62 | 12 | 0 | 7.11E-07 | 1.00E+00 |
| GO:0051306 | mitotic sister chromatid separation | BP | 25 | 8 | 0 | 1.02E-06 | 1.00E+00 |
| GO:1901989 | positive regulation of cell cycle phase transition | BP | 25 | 8 | 0 | 1.02E-06 | 1.00E+00 |
| GO:0010965 | regulation of mitotic sister chromatid separation | BP | 25 | 8 | 0 | 1.02E-06 | 1.00E+00 |
| GO:1901988 | negative regulation of cell cycle phase transition | BP | 53 | 11 | 0 | 1.04E-06 | 1.00E+00 |
| GO:0044771 | meiotic cell cycle phase transition | BP | 4 | 4 | 0 | 1.38E-06 | 1.00E+00 |
| GO:0033045 | regulation of sister chromatid segregation | BP | 26 | 8 | 0 | 1.43E-06 | 1.00E+00 |
| GO:0045930 | negative regulation of mitotic cell cycle | BP | 55 | 11 | 0 | 1.55E-06 | 1.00E+00 |
| GO:0000086 | G2/M transition of mitotic cell cycle | BP | 35 | 9 | 1 | 1.61E-06 | 5.65E-01 |
| GO:0061982 | meiosis I cell cycle process | BP | 27 | 8 | 0 | 1.97E-06 | 1.00E+00 |
| GO:0071174 | mitotic spindle checkpoint signaling | BP | 20 | 7 | 0 | 2.57E-06 | 1.00E+00 |
| GO:2001251 | negative regulation of chromosome organization | BP | 28 | 8 | 0 | 2.68E-06 | 1.00E+00 |
| GO:0000281 | mitotic cytokinesis | BP | 14 | 6 | 1 | 3.57E-06 | 2.82E-01 |
| GO:0031577 | spindle checkpoint signaling | BP | 21 | 7 | 0 | 3.75E-06 | 1.00E+00 |
| GO:1902099 | regulation of metaphase/anaphase transition of cell cycle | BP | 22 | 7 | 0 | 5.35E-06 | 1.00E+00 |
| GO:0030071 | regulation of mitotic metaphase/anaphase transition | BP | 22 | 7 | 0 | 5.35E-06 | 1.00E+00 |
| GO:0022414 | reproductive process | BP | 180 | 19 | 0 | 5.83E-06 | 1.00E+00 |
| GO:0000003 | reproduction | BP | 181 | 19 | 0 | 6.34E-06 | 1.00E+00 |
| GO:0051984 | positive regulation of chromosome segregation | BP | 10 | 5 | 0 | 1.00E-05 | 1.00E+00 |
| GO:0007088 | regulation of mitotic nuclear division | BP | 35 | 8 | 0 | 1.67E-05 | 1.00E+00 |
| GO:0051383 | kinetochore organization | BP | 11 | 5 | 0 | 1.79E-05 | 1.00E+00 |
| GO:0071459 | protein localization to chromosome, centromeric region | BP | 11 | 5 | 0 | 1.79E-05 | 1.00E+00 |
| GO:0007127 | meiosis I | BP | 26 | 7 | 0 | 1.84E-05 | 1.00E+00 |
| GO:0007094 | mitotic spindle assembly checkpoint signaling | BP | 18 | 6 | 0 | 1.97E-05 | 1.00E+00 |
| GO:0071173 | spindle assembly checkpoint signaling | BP | 18 | 6 | 0 | 1.97E-05 | 1.00E+00 |
| GO:0044260 | cellular macromolecule metabolic process | BP | 557 | 36 | 12 | 2.05E-05 | 6.80E-01 |
| GO:0007018 | microtubule-based movement | BP | 37 | 8 | 1 | 2.58E-05 | 5.85E-01 |
| GO:0006260 | DNA replication | BP | 60 | 10 | 1 | 2.70E-05 | 7.62E-01 |
| GO:0033554 | cellular response to stress | BP | 237 | 21 | 5 | 2.79E-05 | 6.64E-01 |
| GO:1902100 | negative regulation of metaphase/anaphase transition of cell cycl | BP | 19 | 6 | 0 | 2.81E-05 | 1.00E+00 |
| GO:0045841 | negative regulation of mitotic metaphase/anaphase transition | BP | 19 | 6 | 0 | 2.81E-05 | 1.00E+00 |
| GO:0070925 | organelle assembly | BP | 132 | 15 | 1 | 2.86E-05 | 9.60E-01 |
| GO:0007143 | female meiotic nuclear division | BP | 12 | 5 | 0 | 2.98E-05 | 1.00E+00 |
| GO:0000910 | cytokinesis | BP | 28 | 7 | 1 | 3.14E-05 | 4.86E-01 |
| GO:1905819 | negative regulation of chromosome separation | BP | 20 | 6 | 0 | 3.90E-05 | 1.00E+00 |
| GO:0033048 | negative regulation of mitotic sister chromatid segregation | BP | 20 | 6 | 0 | 3.90E-05 | 1.00E+00 |
| GO:2000816 | negative regulation of mitotic sister chromatid separation | BP | 20 | 6 | 0 | 3.90E-05 | 1.00E+00 |
| GO:0033046 | negative regulation of sister chromatid segregation | BP | 20 | 6 | 0 | 3.90E-05 | 1.00E+00 |
| GO:0034502 | protein localization to chromosome | BP | 20 | 6 | 0 | 3.90E-05 | 1.00E+00 |
| GO:0033047 | regulation of mitotic sister chromatid segregation | BP | 20 | 6 | 0 | 3.90E-05 | 1.00E+00 |
| GO:0008315 | G2/MI transition of meiotic cell cycle | BP | 3 | 3 | 0 | 4.11E-05 | 1.00E+00 |
| GO:0031536 | positive regulation of exit from mitosis | BP | 3 | 3 | 0 | 4.11E-05 | 1.00E+00 |
| GO:0007076 | mitotic chromosome condensation | BP | 7 | 4 | 0 | 4.45E-05 | 1.00E+00 |
| GO:0048812 | neuron projection morphogenesis | BP | 97 | 0 | 10 | 1.00E+00 | 4.98E-05 |
| GO:0051985 | negative regulation of chromosome segregation | BP | 21 | 6 | 0 | 5.31E-05 | 1.00E+00 |
| GO:0120039 | plasma membrane bounded cell projection morphogenesis | BP | 102 | 0 | 10 | 1.00E+00 | 7.73E-05 |
| GO:1905820 | positive regulation of chromosome separation | BP | 8 | 4 | 0 | 8.66E-05 | 1.00E+00 |
| GO:0048858 | cell projection morphogenesis | BP | 104 | 0 | 10 | 1.00E+00 | 9.15E-05 |
| GO:0006270 | DNA replication initiation | BP | 15 | 5 | 0 | 1.04E-04 | 1.00E+00 |
| GO:0045839 | negative regulation of mitotic nuclear division | BP | 24 | 6 | 0 | 1.21E-04 | 1.00E+00 |
| GO:0051784 | negative regulation of nuclear division | BP | 24 | 6 | 0 | 1.21E-04 | 1.00E+00 |
| GO:0032990 | cell part morphogenesis | BP | 108 | 0 | 10 | 1.00E+00 | 1.27E-04 |
| GO:0006310 | DNA recombination | BP | 58 | 9 | 1 | 1.27E-04 | 7.50E-01 |
| GO:0032506 | cytokinetic process | BP | 9 | 4 | 0 | 1.52E-04 | 1.00E+00 |
| GO:0048667 | cell morphogenesis involved in neuron differentiation | BP | 90 | 0 | 9 | 1.00E+00 | 1.60E-04 |
| GO:0000022 | mitotic spindle elongation | BP | 4 | 3 | 0 | 1.60E-04 | 1.00E+00 |
| GO:0051256 | mitotic spindle midzone assembly | BP | 4 | 3 | 0 | 1.60E-04 | 1.00E+00 |
| GO:0051231 | spindle elongation | BP | 4 | 3 | 0 | 1.60E-04 | 1.00E+00 |
| GO:0051255 | spindle midzone assembly | BP | 4 | 3 | 0 | 1.60E-04 | 1.00E+00 |
| GO:0010638 | positive regulation of organelle organization | BP | 63 | 9 | 0 | 2.45E-04 | 1.00E+00 |
| GO:1902749 | regulation of cell cycle G2/M phase transition | BP | 27 | 6 | 1 | 2.45E-04 | 4.73E-01 |
| GO:0010389 | regulation of G2/M transition of mitotic cell cycle | BP | 27 | 6 | 1 | 2.45E-04 | 4.73E-01 |
| GO:1903083 | protein localization to condensed chromosome | BP | 10 | 4 | 0 | 2.46E-04 | 1.00E+00 |
| GO:0034501 | protein localization to kinetochore | BP | 10 | 4 | 0 | 2.46E-04 | 1.00E+00 |
| GO:0051445 | regulation of meiotic cell cycle | BP | 10 | 4 | 0 | 2.46E-04 | 1.00E+00 |
| GO:0010639 | negative regulation of organelle organization | BP | 65 | 9 | 0 | 3.12E-04 | 1.00E+00 |
| GO:0007095 | mitotic G2 DNA damage checkpoint signaling | BP | 11 | 4 | 0 | 3.77E-04 | 1.00E+00 |
| GO:0032989 | cellular component morphogenesis | BP | 123 | 1 | 10 | 9.89E-01 | 3.77E-04 |
| GO:0007056 | spindle assembly involved in female meiosis | BP | 5 | 3 | 0 | 3.90E-04 | 1.00E+00 |
| GO:0007057 | spindle assembly involved in female meiosis I | BP | 5 | 3 | 0 | 3.90E-04 | 1.00E+00 |
| GO:0034329 | cell junction assembly | BP | 82 | 0 | 8 | 1.00E+00 | 4.62E-04 |
| GO:0045132 | meiotic chromosome segregation | BP | 20 | 5 | 0 | 4.68E-04 | 1.00E+00 |
| GO:0048666 | neuron development | BP | 179 | 1 | 12 | 9.99E-01 | 5.52E-04 |
| GO:0006261 | DNA-templated DNA replication | BP | 43 | 7 | 1 | 5.61E-04 | 6.41E-01 |
| GO:0034330 | cell junction organization | BP | 130 | 1 | 10 | 9.92E-01 | 5.92E-04 |
| GO:0051965 | positive regulation of synapse assembly | BP | 31 | 0 | 5 | 1.00E+00 | 6.00E-04 |
| GO:0098660 | inorganic ion transmembrane transport | BP | 86 | 0 | 8 | 1.00E+00 | 6.41E-04 |
| GO:0007144 | female meiosis I | BP | 6 | 3 | 0 | 7.61E-04 | 1.00E+00 |
| GO:0090306 | meiotic spindle assembly | BP | 6 | 3 | 0 | 7.61E-04 | 1.00E+00 |
| GO:0007096 | regulation of exit from mitosis | BP | 6 | 3 | 0 | 7.61E-04 | 1.00E+00 |
| GO:1904029 | regulation of cyclin-dependent protein kinase activity | BP | 13 | 4 | 0 | 7.74E-04 | 1.00E+00 |
| GO:0000079 | regulation of cyclin-dependent protein serine/threonine kinase ac | BP | 13 | 4 | 0 | 7.74E-04 | 1.00E+00 |
| GO:0007062 | sister chromatid cohesion | BP | 13 | 4 | 0 | 7.74E-04 | 1.00E+00 |
| GO:0007409 | axonogenesis | BP | 68 | 0 | 7 | 1.00E+00 | 7.87E-04 |
| GO:0042391 | regulation of membrane potential | BP | 50 | 0 | 6 | 1.00E+00 | 8.52E-04 |
| GO:0031175 | neuron projection development | BP | 162 | 1 | 11 | 9.98E-01 | 8.86E-04 |
| GO:0051338 | regulation of transferase activity | BP | 108 | 11 | 4 | 1.00E-03 | 2.42E-01 |
| GO:0090304 | nucleic acid metabolic process | BP | 487 | 29 | 8 | 1.04E-03 | 9.12E-01 |
| GO:0044818 | mitotic G2/M transition checkpoint | BP | 14 | 4 | 0 | 1.06E-03 | 1.00E+00 |
| GO:1902750 | negative regulation of cell cycle G2/M phase transition | BP | 14 | 4 | 0 | 1.06E-03 | 1.00E+00 |
| GO:0010972 | negative regulation of G2/M transition of mitotic cell cycle | BP | 14 | 4 | 0 | 1.06E-03 | 1.00E+00 |
| GO:2001252 | positive regulation of chromosome organization | BP | 14 | 4 | 0 | 1.06E-03 | 1.00E+00 |
| GO:0000902 | cell morphogenesis | BP | 168 | 2 | 11 | 9.85E-01 | 1.21E-03 |
| GO:0099607 | lateral attachment of mitotic spindle microtubules to kinetochore | BP | 2 | 2 | 0 | 1.21E-03 | 1.00E+00 |
| GO:1901993 | regulation of meiotic cell cycle phase transition | BP | 2 | 2 | 0 | 1.21E-03 | 1.00E+00 |
| GO:0090235 | regulation of metaphase plate congression | BP | 2 | 2 | 0 | 1.21E-03 | 1.00E+00 |
| GO:0000727 | double-strand break repair via break-induced replication | BP | 7 | 3 | 0 | 1.30E-03 | 1.00E+00 |
| GO:0010458 | exit from mitosis | BP | 7 | 3 | 0 | 1.30E-03 | 1.00E+00 |
| GO:0000212 | meiotic spindle organization | BP | 7 | 3 | 0 | 1.30E-03 | 1.00E+00 |
| GO:0090224 | regulation of spindle organization | BP | 7 | 3 | 0 | 1.30E-03 | 1.00E+00 |
| GO:0061564 | axon development | BP | 74 | 0 | 7 | 1.00E+00 | 1.31E-03 |
| GO:0016310 | phosphorylation | BP | 205 | 16 | 5 | 1.35E-03 | 5.29E-01 |
| GO:0042770 | signal transduction in response to DNA damage | BP | 25 | 5 | 0 | 1.40E-03 | 1.00E+00 |
| GO:0000904 | cell morphogenesis involved in differentiation | BP | 121 | 1 | 9 | 9.88E-01 | 1.48E-03 |
| GO:1901890 | positive regulation of cell junction assembly | BP | 38 | 0 | 5 | 1.00E+00 | 1.57E-03 |
| GO:0046548 | retinal rod cell development | BP | 3 | 0 | 2 | 1.00E+00 | 1.57E-03 |
| GO:0006941 | striated muscle contraction | BP | 23 | 0 | 4 | 1.00E+00 | 1.66E-03 |
| GO:0018105 | peptidyl-serine phosphorylation | BP | 26 | 5 | 2 | 1.69E-03 | 1.21E-01 |
| GO:0051963 | regulation of synapse assembly | BP | 39 | 0 | 5 | 1.00E+00 | 1.77E-03 |
| GO:0098662 | inorganic cation transmembrane transport | BP | 78 | 0 | 7 | 1.00E+00 | 1.80E-03 |
| GO:0006468 | protein phosphorylation | BP | 173 | 14 | 5 | 2.01E-03 | 3.78E-01 |
| GO:0007098 | centrosome cycle | BP | 27 | 5 | 1 | 2.01E-03 | 4.73E-01 |
| GO:0051128 | regulation of cellular component organization | BP | 343 | 22 | 11 | 2.13E-03 | 1.62E-01 |
| GO:0051054 | positive regulation of DNA metabolic process | BP | 40 | 6 | 0 | 2.21E-03 | 1.00E+00 |
| GO:2001022 | positive regulation of response to DNA damage stimulus | BP | 17 | 4 | 0 | 2.32E-03 | 1.00E+00 |
| GO:0051649 | establishment of localization in cell | BP | 219 | 16 | 10 | 2.73E-03 | 2.60E-02 |
| GO:0051052 | regulation of DNA metabolic process | BP | 71 | 8 | 1 | 2.73E-03 | 8.18E-01 |
| GO:0031023 | microtubule organizing center organization | BP | 29 | 5 | 1 | 2.80E-03 | 4.98E-01 |
| GO:0006950 | response to stress | BP | 397 | 24 | 5 | 2.88E-03 | 9.69E-01 |
| GO:0044773 | mitotic DNA damage checkpoint signaling | BP | 18 | 4 | 0 | 2.90E-03 | 1.00E+00 |
| GO:0016358 | dendrite development | BP | 44 | 1 | 5 | 7.95E-01 | 3.06E-03 |
| GO:0098870 | action potential propagation | BP | 4 | 0 | 2 | 1.00E+00 | 3.10E-03 |
| GO:0019227 | neuronal action potential propagation | BP | 4 | 0 | 2 | 1.00E+00 | 3.10E-03 |
| GO:0030182 | neuron differentiation | BP | 218 | 2 | 12 | 9.97E-01 | 3.19E-03 |
| GO:0034220 | ion transmembrane transport | BP | 110 | 0 | 8 | 1.00E+00 | 3.23E-03 |
| GO:0007416 | synapse assembly | BP | 45 | 0 | 5 | 1.00E+00 | 3.38E-03 |
| GO:0022403 | cell cycle phase | BP | 3 | 2 | 0 | 3.54E-03 | 1.00E+00 |
| GO:0016321 | female meiosis chromosome segregation | BP | 3 | 2 | 0 | 3.54E-03 | 1.00E+00 |
| GO:0061952 | midbody abscission | BP | 3 | 2 | 0 | 3.54E-03 | 1.00E+00 |
| GO:1905463 | negative regulation of DNA duplex unwinding | BP | 3 | 2 | 0 | 3.54E-03 | 1.00E+00 |
| GO:0051987 | positive regulation of attachment of spindle microtubules to kinet | BP | 3 | 2 | 0 | 3.54E-03 | 1.00E+00 |
| GO:1904355 | positive regulation of telomere capping | BP | 3 | 2 | 0 | 3.54E-03 | 1.00E+00 |
| GO:0071103 | DNA conformation change | BP | 19 | 4 | 0 | 3.58E-03 | 1.00E+00 |
| GO:0000077 | DNA damage checkpoint signaling | BP | 19 | 4 | 0 | 3.58E-03 | 1.00E+00 |
| GO:0032508 | DNA duplex unwinding | BP | 19 | 4 | 0 | 3.58E-03 | 1.00E+00 |
| GO:0032392 | DNA geometric change | BP | 19 | 4 | 0 | 3.58E-03 | 1.00E+00 |
| GO:0007292 | female gamete generation | BP | 31 | 5 | 0 | 3.80E-03 | 1.00E+00 |
| GO:0050808 | synapse organization | BP | 89 | 1 | 7 | 9.61E-01 | 3.85E-03 |
| GO:0006268 | DNA unwinding involved in DNA replication | BP | 10 | 3 | 0 | 4.12E-03 | 1.00E+00 |
| GO:1902751 | positive regulation of cell cycle G2/M phase transition | BP | 10 | 3 | 0 | 4.12E-03 | 1.00E+00 |
| GO:0032467 | positive regulation of cytokinesis | BP | 10 | 3 | 0 | 4.12E-03 | 1.00E+00 |
| GO:0010971 | positive regulation of G2/M transition of mitotic cell cycle | BP | 10 | 3 | 0 | 4.12E-03 | 1.00E+00 |
| GO:0065008 | regulation of biological quality | BP | 455 | 9 | 19 | 9.88E-01 | 4.30E-03 |
| GO:0032465 | regulation of cytokinesis | BP | 20 | 4 | 0 | 4.36E-03 | 1.00E+00 |
| GO:0051283 | negative regulation of sequestering of calcium ion | BP | 15 | 0 | 3 | 1.00E+00 | 4.44E-03 |
| GO:0051282 | regulation of sequestering of calcium ion | BP | 15 | 0 | 3 | 1.00E+00 | 4.44E-03 |
| GO:0051209 | release of sequestered calcium ion into cytosol | BP | 15 | 0 | 3 | 1.00E+00 | 4.44E-03 |
| GO:0098655 | cation transmembrane transport | BP | 92 | 0 | 7 | 1.00E+00 | 4.64E-03 |
| GO:0098916 | anterograde trans-synaptic signaling | BP | 93 | 1 | 7 | 9.66E-01 | 4.93E-03 |
| GO:0007268 | chemical synaptic transmission | BP | 93 | 1 | 7 | 9.66E-01 | 4.93E-03 |
| GO:0006302 | double-strand break repair | BP | 62 | 7 | 1 | 5.03E-03 | 7.73E-01 |
| GO:0042462 | eye photoreceptor cell development | BP | 5 | 0 | 2 | 1.00E+00 | 5.09E-03 |
| GO:0060221 | retinal rod cell differentiation | BP | 5 | 0 | 2 | 1.00E+00 | 5.09E-03 |
| GO:0044774 | mitotic DNA integrity checkpoint signaling | BP | 21 | 4 | 0 | 5.24E-03 | 1.00E+00 |
| GO:0048699 | generation of neurons | BP | 232 | 3 | 12 | 9.91E-01 | 5.37E-03 |
| GO:0051208 | sequestering of calcium ion | BP | 16 | 0 | 3 | 1.00E+00 | 5.38E-03 |
| GO:0051293 | establishment of spindle localization | BP | 11 | 3 | 0 | 5.53E-03 | 1.00E+00 |
| GO:0051785 | positive regulation of nuclear division | BP | 11 | 3 | 0 | 5.53E-03 | 1.00E+00 |
| GO:0044843 | cell cycle G1/S phase transition | BP | 34 | 5 | 0 | 5.73E-03 | 1.00E+00 |
| GO:0018209 | peptidyl-serine modification | BP | 34 | 5 | 2 | 5.73E-03 | 1.87E-01 |
| GO:0051962 | positive regulation of nervous system development | BP | 73 | 0 | 6 | 1.00E+00 | 6.07E-03 |
| GO:0006811 | ion transport | BP | 177 | 1 | 10 | 9.99E-01 | 6.22E-03 |
| GO:0099537 | trans-synaptic signaling | BP | 97 | 1 | 7 | 9.71E-01 | 6.23E-03 |
| GO:0009987 | cellular process | BP | ### | 69 | 44 | 6.26E-03 | 1.82E-01 |
| GO:0006139 | nucleobase-containing compound metabolic process | BP | 545 | 29 | 8 | 6.53E-03 | 9.63E-01 |
| GO:0099536 | synaptic signaling | BP | 98 | 1 | 7 | 9.72E-01 | 6.59E-03 |
| GO:0050790 | regulation of catalytic activity | BP | 197 | 14 | 6 | 6.73E-03 | 3.08E-01 |
| GO:0031145 | anaphase-promoting complex-dependent catabolic process | BP | 4 | 2 | 0 | 6.91E-03 | 1.00E+00 |
| GO:0044848 | biological phase | BP | 4 | 2 | 0 | 6.91E-03 | 1.00E+00 |
| GO:0010032 | meiotic chromosome condensation | BP | 4 | 2 | 0 | 6.91E-03 | 1.00E+00 |
| GO:1902410 | mitotic cytokinetic process | BP | 4 | 2 | 0 | 6.91E-03 | 1.00E+00 |
| GO:1902042 | negative regulation of extrinsic apoptotic signaling pathway via d | BP | 4 | 2 | 0 | 6.91E-03 | 1.00E+00 |
| GO:0051444 | negative regulation of ubiquitin-protein transferase activity | BP | 4 | 2 | 0 | 6.91E-03 | 1.00E+00 |
| GO:1905821 | positive regulation of chromosome condensation | BP | 4 | 2 | 0 | 6.91E-03 | 1.00E+00 |
| GO:1905462 | regulation of DNA duplex unwinding | BP | 4 | 2 | 0 | 6.91E-03 | 1.00E+00 |
| GO:0033365 | protein localization to organelle | BP | 100 | 9 | 0 | 6.92E-03 | 1.00E+00 |
| GO:0045739 | positive regulation of DNA repair | BP | 12 | 3 | 0 | 7.19E-03 | 1.00E+00 |
| GO:0044380 | protein localization to cytoskeleton | BP | 12 | 3 | 0 | 7.19E-03 | 1.00E+00 |
| GO:0072698 | protein localization to microtubule cytoskeleton | BP | 12 | 3 | 0 | 7.19E-03 | 1.00E+00 |
| GO:0051653 | spindle localization | BP | 12 | 3 | 0 | 7.19E-03 | 1.00E+00 |
| GO:0071478 | cellular response to radiation | BP | 23 | 4 | 2 | 7.35E-03 | 9.88E-02 |
| GO:0031570 | DNA integrity checkpoint signaling | BP | 23 | 4 | 0 | 7.35E-03 | 1.00E+00 |
| GO:1901888 | regulation of cell junction assembly | BP | 54 | 0 | 5 | 1.00E+00 | 7.47E-03 |
| GO:0060219 | camera-type eye photoreceptor cell differentiation | BP | 6 | 0 | 2 | 1.00E+00 | 7.52E-03 |
| GO:0003009 | skeletal muscle contraction | BP | 6 | 0 | 2 | 1.00E+00 | 7.52E-03 |
| GO:0097553 | calcium ion transmembrane import into cytosol | BP | 18 | 0 | 3 | 1.00E+00 | 7.59E-03 |
| GO:0050773 | regulation of dendrite development | BP | 18 | 1 | 3 | 4.75E-01 | 7.59E-03 |
| GO:0120036 | plasma membrane bounded cell projection organization | BP | 245 | 2 | 12 | 9.99E-01 | 8.35E-03 |
| GO:0022412 | cellular process involved in reproduction in multicellular organism | BP | 52 | 6 | 0 | 8.43E-03 | 1.00E+00 |
| GO:0044089 | positive regulation of cellular component biogenesis | BP | 79 | 2 | 6 | 7.74E-01 | 8.92E-03 |
| GO:0051641 | cellular localization | BP | 315 | 19 | 12 | 9.31E-03 | 5.22E-02 |
| GO:0050807 | regulation of synapse organization | BP | 57 | 1 | 5 | 8.72E-01 | 9.39E-03 |
| GO:0050803 | regulation of synapse structure or activity | BP | 57 | 1 | 5 | 8.72E-01 | 9.39E-03 |
| GO:0044092 | negative regulation of molecular function | BP | 87 | 8 | 1 | 9.57E-03 | 8.77E-01 |
| GO:0051094 | positive regulation of developmental process | BP | 219 | 5 | 11 | 8.97E-01 | 9.81E-03 |
| GO:0065004 | protein-DNA complex assembly | BP | 25 | 4 | 0 | 9.97E-03 | 1.00E+00 |
| GO:2000241 | regulation of reproductive process | BP | 25 | 4 | 0 | 9.97E-03 | 1.00E+00 |
| GO:0000724 | double-strand break repair via homologous recombination | BP | 39 | 5 | 1 | 1.04E-02 | 6.05E-01 |
| GO:0000725 | recombinational repair | BP | 39 | 5 | 1 | 1.04E-02 | 6.05E-01 |
| GO:0001754 | eye photoreceptor cell differentiation | BP | 7 | 0 | 2 | 1.00E+00 | 1.04E-02 |
| GO:0007187 | G protein-coupled receptor signaling pathway, coupled to cyclic | BP | 7 | 0 | 2 | 1.00E+00 | 1.04E-02 |
| GO:0042461 | photoreceptor cell development | BP | 7 | 0 | 2 | 1.00E+00 | 1.04E-02 |
| GO:0045859 | regulation of protein kinase activity | BP | 71 | 7 | 3 | 1.06E-02 | 2.28E-01 |
| GO:0051130 | positive regulation of cellular component organization | BP | 162 | 10 | 9 | 5.18E-02 | 1.07E-02 |
| GO:0030030 | cell projection organization | BP | 254 | 2 | 12 | 9.99E-01 | 1.11E-02 |
| GO:0030001 | metal ion transport | BP | 108 | 0 | 7 | 1.00E+00 | 1.11E-02 |
| GO:0007064 | mitotic sister chromatid cohesion | BP | 5 | 2 | 0 | 1.13E-02 | 1.00E+00 |
| GO:0051446 | positive regulation of meiotic cell cycle | BP | 5 | 2 | 0 | 1.13E-02 | 1.00E+00 |
| GO:0051973 | positive regulation of telomerase activity | BP | 5 | 2 | 0 | 1.13E-02 | 1.00E+00 |
| GO:0060623 | regulation of chromosome condensation | BP | 5 | 2 | 0 | 1.13E-02 | 1.00E+00 |
| GO:0035825 | homologous recombination | BP | 14 | 3 | 0 | 1.13E-02 | 1.00E+00 |
| GO:0018107 | peptidyl-threonine phosphorylation | BP | 14 | 3 | 1 | 1.13E-02 | 2.82E-01 |
| GO:0140527 | reciprocal homologous recombination | BP | 14 | 3 | 0 | 1.13E-02 | 1.00E+00 |
| GO:0007131 | reciprocal meiotic recombination | BP | 14 | 3 | 0 | 1.13E-02 | 1.00E+00 |
| GO:0046483 | heterocycle metabolic process | BP | 566 | 29 | 8 | 1.15E-02 | 9.74E-01 |
| GO:0043170 | macromolecule metabolic process | BP | 917 | 42 | 16 | 1.16E-02 | 9.59E-01 |
| GO:0010976 | positive regulation of neuron projection development | BP | 39 | 0 | 4 | 1.00E+00 | 1.18E-02 |
| GO:0001508 | action potential | BP | 21 | 0 | 3 | 1.00E+00 | 1.18E-02 |
| GO:0060402 | calcium ion transport into cytosol | BP | 21 | 0 | 3 | 1.00E+00 | 1.18E-02 |
| GO:0006796 | phosphate-containing compound metabolic process | BP | 300 | 18 | 6 | 1.22E-02 | 7.22E-01 |
| GO:0006464 | cellular protein modification process | BP | 324 | 19 | 8 | 1.26E-02 | 4.91E-01 |
| GO:0036211 | protein modification process | BP | 324 | 19 | 8 | 1.26E-02 | 4.91E-01 |
| GO:0010975 | regulation of neuron projection development | BP | 85 | 1 | 6 | 9.55E-01 | 1.26E-02 |
| GO:0071824 | protein-DNA complex subunit organization | BP | 27 | 4 | 0 | 1.31E-02 | 1.00E+00 |
| GO:0086001 | cardiac muscle cell action potential | BP | 8 | 0 | 2 | 1.00E+00 | 1.36E-02 |
| GO:0018198 | peptidyl-cysteine modification | BP | 8 | 0 | 2 | 1.00E+00 | 1.36E-02 |
| GO:1900006 | positive regulation of dendrite development | BP | 8 | 0 | 2 | 1.00E+00 | 1.36E-02 |
| GO:0033135 | regulation of peptidyl-serine phosphorylation | BP | 8 | 0 | 2 | 1.00E+00 | 1.36E-02 |
| GO:0051298 | centrosome duplication | BP | 15 | 3 | 0 | 1.38E-02 | 1.00E+00 |
| GO:0070192 | chromosome organization involved in meiotic cell cycle | BP | 15 | 3 | 0 | 1.38E-02 | 1.00E+00 |
| GO:0051129 | negative regulation of cellular component organization | BP | 112 | 9 | 2 | 1.42E-02 | 7.47E-01 |
| GO:0006793 | phosphorus metabolic process | BP | 306 | 18 | 6 | 1.49E-02 | 7.40E-01 |
| GO:0031346 | positive regulation of cell projection organization | BP | 64 | 0 | 5 | 1.00E+00 | 1.51E-02 |
| GO:0060401 | cytosolic calcium ion transport | BP | 23 | 0 | 3 | 1.00E+00 | 1.52E-02 |
| GO:0006936 | muscle contraction | BP | 42 | 0 | 4 | 1.00E+00 | 1.52E-02 |
| GO:0032501 | multicellular organismal process | BP | 864 | 22 | 28 | 9.84E-01 | 1.53E-02 |
| GO:0006725 | cellular aromatic compound metabolic process | BP | 578 | 29 | 8 | 1.55E-02 | 9.79E-01 |
| GO:0051307 | meiotic chromosome separation | BP | 6 | 2 | 0 | 1.65E-02 | 1.00E+00 |
| GO:0033314 | mitotic DNA replication checkpoint signaling | BP | 6 | 2 | 0 | 1.65E-02 | 1.00E+00 |
| GO:0032873 | negative regulation of stress-activated MAPK cascade | BP | 6 | 2 | 0 | 1.65E-02 | 1.00E+00 |
| GO:0070303 | negative regulation of stress-activated protein kinase signaling ca | BP | 6 | 2 | 0 | 1.65E-02 | 1.00E+00 |
| GO:0032212 | positive regulation of telomere maintenance via telomerase | BP | 6 | 2 | 0 | 1.65E-02 | 1.00E+00 |
| GO:0051865 | protein autoubiquitination | BP | 6 | 2 | 0 | 1.65E-02 | 1.00E+00 |
| GO:0040020 | regulation of meiotic nuclear division | BP | 6 | 2 | 0 | 1.65E-02 | 1.00E+00 |
| GO:0060236 | regulation of mitotic spindle organization | BP | 6 | 2 | 0 | 1.65E-02 | 1.00E+00 |
| GO:0051972 | regulation of telomerase activity | BP | 6 | 2 | 0 | 1.65E-02 | 1.00E+00 |
| GO:1904353 | regulation of telomere capping | BP | 6 | 2 | 0 | 1.65E-02 | 1.00E+00 |
| GO:0048468 | cell development | BP | 335 | 5 | 14 | 9.95E-01 | 1.68E-02 |
| GO:0000082 | G1/S transition of mitotic cell cycle | BP | 29 | 4 | 0 | 1.68E-02 | 1.00E+00 |
| GO:0008104 | protein localization | BP | 289 | 17 | 8 | 1.81E-02 | 3.58E-01 |
| GO:0031400 | negative regulation of protein modification process | BP | 45 | 5 | 1 | 1.87E-02 | 6.58E-01 |
| GO:0000302 | response to reactive oxygen species | BP | 25 | 0 | 3 | 1.00E+00 | 1.91E-02 |
| GO:0022607 | cellular component assembly | BP | 362 | 20 | 14 | 1.94E-02 | 3.15E-02 |
| GO:0051348 | negative regulation of transferase activity | BP | 17 | 3 | 0 | 1.96E-02 | 1.00E+00 |
| GO:0022008 | neurogenesis | BP | 274 | 5 | 12 | 9.73E-01 | 1.97E-02 |

| | | | | | | | |
|---|---|---|---|---|---|---|---|
| GO:0043412 | macromolecule modification | BP | 339 | 19 | 8 | 1.99E-02 | 5.46E-01 |
| GO:0035556 | intracellular signal transduction | BP | 293 | 17 | 7 | 2.06E-02 | 5.36E-01 |
| GO:0098742 | cell-cell adhesion via plasma-membrane adhesion molecules | BP | 46 | 0 | 4 | 1.00E+00 | 2.07E-02 |
| GO:0044085 | cellular component biogenesis | BP | 379 | 20 | 15 | 3.09E-02 | 2.10E-02 |
| GO:0048813 | dendrite morphogenesis | BP | 26 | 0 | 3 | 1.00E+00 | 2.12E-02 |
| GO:0007156 | homophilic cell adhesion via plasma membrane adhesion molecu | BP | 26 | 0 | 3 | 1.00E+00 | 2.12E-02 |
| GO:0016339 | calcium-dependent cell-cell adhesion via plasma membrane cell a | BP | 10 | 0 | 2 | 1.00E+00 | 2.13E-02 |
| GO:0021954 | central nervous system neuron development | BP | 10 | 0 | 2 | 1.00E+00 | 2.13E-02 |
| GO:0050879 | multicellular organismal movement | BP | 10 | 0 | 2 | 1.00E+00 | 2.13E-02 |
| GO:0050881 | musculoskeletal movement | BP | 10 | 0 | 2 | 1.00E+00 | 2.13E-02 |
| GO:0003407 | neural retina development | BP | 10 | 0 | 2 | 1.00E+00 | 2.13E-02 |
| GO:0046530 | photoreceptor cell differentiation | BP | 10 | 0 | 2 | 1.00E+00 | 2.13E-02 |
| GO:0006325 | chromatin organization | BP | 47 | 5 | 1 | 2.22E-02 | 6.74E-01 |
| GO:0007276 | gamete generation | BP | 82 | 7 | 0 | 2.23E-02 | 1.00E+00 |
| GO:0007155 | cell adhesion | BP | 246 | 2 | 11 | 9.99E-01 | 2.24E-02 |
| GO:0034644 | cellular response to UV | BP | 7 | 2 | 1 | 2.26E-02 | 1.52E-01 |
| GO:0016572 | histone phosphorylation | BP | 7 | 2 | 0 | 2.26E-02 | 1.00E+00 |
| GO:0051382 | kinetochore assembly | BP | 7 | 2 | 0 | 2.26E-02 | 1.00E+00 |
| GO:0045840 | positive regulation of mitotic nuclear division | BP | 7 | 2 | 0 | 2.26E-02 | 1.00E+00 |
| GO:1904358 | positive regulation of telomere maintenance via telomere lengther | BP | 7 | 2 | 0 | 2.26E-02 | 1.00E+00 |
| GO:1902041 | regulation of extrinsic apoptotic signaling pathway via death dom | BP | 7 | 2 | 0 | 2.26E-02 | 1.00E+00 |
| GO:0032210 | regulation of telomere maintenance via telomerase | BP | 7 | 2 | 0 | 2.26E-02 | 1.00E+00 |
| GO:0051438 | regulation of ubiquitin-protein transferase activity | BP | 7 | 2 | 0 | 2.26E-02 | 1.00E+00 |
| GO:0018210 | peptidyl-threonine modification | BP | 18 | 3 | 1 | 2.29E-02 | 3.47E-01 |
| GO:0007197 | adenylate cyclase-inhibiting G protein-coupled acetylcholine rece | BP | 1 | 0 | 1 | 1.00E+00 | 2.33E-02 |
| GO:0035881 | amacrine cell differentiation | BP | 1 | 0 | 1 | 1.00E+00 | 2.33E-02 |
| GO:0031547 | brain-derived neurotrophic factor receptor signaling pathway | BP | 1 | 0 | 1 | 1.00E+00 | 2.33E-02 |
| GO:0048791 | calcium ion-regulated exocytosis of neurotransmitter | BP | 1 | 0 | 1 | 1.00E+00 | 2.33E-02 |
| GO:1990416 | cellular response to brain-derived neurotrophic factor stimulus | BP | 1 | 0 | 1 | 1.00E+00 | 2.33E-02 |
| GO:0048669 | collateral sprouting in absence of injury | BP | 1 | 0 | 1 | 1.00E+00 | 2.33E-02 |
| GO:0003335 | corneocyte development | BP | 1 | 0 | 1 | 1.00E+00 | 2.33E-02 |
| GO:1990504 | dense core granule exocytosis | BP | 1 | 0 | 1 | 1.00E+00 | 2.33E-02 |
| GO:0061789 | dense core granule priming | BP | 1 | 0 | 1 | 1.00E+00 | 2.33E-02 |
| GO:0030205 | dermatan sulfate metabolic process | BP | 1 | 0 | 1 | 1.00E+00 | 2.33E-02 |
| GO:0050655 | dermatan sulfate proteoglycan metabolic process | BP | 1 | 0 | 1 | 1.00E+00 | 2.33E-02 |
| GO:0008065 | establishment of blood-nerve barrier | BP | 1 | 0 | 1 | 1.00E+00 | 2.33E-02 |
| GO:0060857 | establishment of glial blood-brain barrier | BP | 1 | 0 | 1 | 1.00E+00 | 2.33E-02 |
| GO:0031443 | fast-twitch skeletal muscle fiber contraction | BP | 1 | 0 | 1 | 1.00E+00 | 2.33E-02 |
| GO:0007213 | G protein-coupled acetylcholine receptor signaling pathway | BP | 1 | 0 | 1 | 1.00E+00 | 2.33E-02 |
| GO:0034436 | glycoprotein transport | BP | 1 | 0 | 1 | 1.00E+00 | 2.33E-02 |
| GO:0036077 | intratendonous ossification | BP | 1 | 0 | 1 | 1.00E+00 | 2.33E-02 |
| GO:1902083 | negative regulation of peptidyl-cysteine S-nitrosylation | BP | 1 | 0 | 1 | 1.00E+00 | 2.33E-02 |
| GO:0099011 | neuronal dense core vesicle exocytosis | BP | 1 | 0 | 1 | 1.00E+00 | 2.33E-02 |
| GO:0045213 | neurotransmitter receptor metabolic process | BP | 1 | 0 | 1 | 1.00E+00 | 2.33E-02 |
| GO:0021627 | olfactory nerve morphogenesis | BP | 1 | 0 | 1 | 1.00E+00 | 2.33E-02 |
| GO:0021629 | olfactory nerve structural organization | BP | 1 | 0 | 1 | 1.00E+00 | 2.33E-02 |
| GO:0018119 | peptidyl-cysteine S-nitrosylation | BP | 1 | 0 | 1 | 1.00E+00 | 2.33E-02 |
| GO:1904028 | positive regulation of collagen fibril organization | BP | 1 | 0 | 1 | 1.00E+00 | 2.33E-02 |
| GO:1900020 | positive regulation of protein kinase C activity | BP | 1 | 0 | 1 | 1.00E+00 | 2.33E-02 |
| GO:2000651 | positive regulation of sodium ion transmembrane transporter acti | BP | 1 | 0 | 1 | 1.00E+00 | 2.33E-02 |
| GO:0099525 | presynaptic dense core vesicle exocytosis | BP | 1 | 0 | 1 | 1.00E+00 | 2.33E-02 |
| GO:0038026 | reelin-mediated signaling pathway | BP | 1 | 0 | 1 | 1.00E+00 | 2.33E-02 |
| GO:0010882 | regulation of cardiac muscle contraction by calcium ion signaling | BP | 1 | 0 | 1 | 1.00E+00 | 2.33E-02 |
| GO:0010881 | regulation of cardiac muscle contraction by regulation of the relea | BP | 1 | 0 | 1 | 1.00E+00 | 2.33E-02 |
| GO:0110026 | regulation of DNA strand resection involved in replication fork pro | BP | 1 | 0 | 1 | 1.00E+00 | 2.33E-02 |
| GO:2000169 | regulation of peptidyl-cysteine S-nitrosylation | BP | 1 | 0 | 1 | 1.00E+00 | 2.33E-02 |
| GO:0044375 | regulation of peroxisome size | BP | 1 | 0 | 1 | 1.00E+00 | 2.33E-02 |
| GO:1900019 | regulation of protein kinase C activity | BP | 1 | 0 | 1 | 1.00E+00 | 2.33E-02 |
| GO:0014722 | regulation of skeletal muscle contraction by calcium ion signaling | BP | 1 | 0 | 1 | 1.00E+00 | 2.33E-02 |
| GO:0014809 | regulation of skeletal muscle contraction by regulation of release | BP | 1 | 0 | 1 | 1.00E+00 | 2.33E-02 |
| GO:0099538 | synaptic signaling via neuropeptide | BP | 1 | 0 | 1 | 1.00E+00 | 2.33E-02 |
| GO:0016082 | synaptic vesicle priming | BP | 1 | 0 | 1 | 1.00E+00 | 2.33E-02 |
| GO:0002932 | tendon sheath development | BP | 1 | 0 | 1 | 1.00E+00 | 2.33E-02 |
| GO:0099540 | trans-synaptic signaling by neuropeptide | BP | 1 | 0 | 1 | 1.00E+00 | 2.33E-02 |
| GO:0099551 | trans-synaptic signaling by neuropeptide, modulating synaptic tr | BP | 1 | 0 | 1 | 1.00E+00 | 2.33E-02 |
| GO:0007021 | tubulin complex assembly | BP | 1 | 0 | 1 | 1.00E+00 | 2.33E-02 |
| GO:0014721 | twitch skeletal muscle contraction | BP | 1 | 0 | 1 | 1.00E+00 | 2.33E-02 |
| GO:0034447 | very-low-density lipoprotein particle clearance | BP | 1 | 0 | 1 | 1.00E+00 | 2.33E-02 |
| GO:0003010 | voluntary skeletal muscle contraction | BP | 1 | 0 | 1 | 1.00E+00 | 2.33E-02 |
| GO:0030534 | adult behavior | BP | 27 | 0 | 3 | 1.00E+00 | 2.35E-02 |
| GO:0051651 | maintenance of location in cell | BP | 27 | 0 | 3 | 1.00E+00 | 2.35E-02 |
| GO:0071805 | potassium ion transmembrane transport | BP | 27 | 0 | 3 | 1.00E+00 | 2.35E-02 |
| GO:0034641 | cellular nitrogen compound metabolic process | BP | 622 | 30 | 9 | 2.36E-02 | 9.76E-01 |
| GO:0016567 | protein ubiquitination | BP | 48 | 5 | 0 | 2.42E-02 | 1.00E+00 |
| GO:1901360 | organic cyclic compound metabolic process | BP | 597 | 29 | 9 | 2.42E-02 | 9.64E-01 |
| GO:0055085 | transmembrane transport | BP | 155 | 1 | 8 | 9.97E-01 | 2.43E-02 |
| GO:0031399 | regulation of protein modification process | BP | 164 | 11 | 5 | 2.45E-02 | 3.34E-01 |
| GO:0007193 | adenylate cyclase-inhibiting G protein-coupled receptor signaling | BP | 11 | 0 | 2 | 1.00E+00 | 2.56E-02 |
| GO:0034332 | adherens junction organization | BP | 11 | 0 | 2 | 1.00E+00 | 2.56E-02 |
| GO:0045661 | regulation of myoblast differentiation | BP | 11 | 0 | 2 | 1.00E+00 | 2.56E-02 |
| GO:0051279 | regulation of release of sequestered calcium ion into cytosol | BP | 11 | 0 | 2 | 1.00E+00 | 2.56E-02 |
| GO:0032963 | collagen metabolic process | BP | 28 | 0 | 3 | 1.00E+00 | 2.59E-02 |
| GO:0001819 | positive regulation of cytokine production | BP | 33 | 4 | 0 | 2.61E-02 | 1.00E+00 |
| GO:0043086 | negative regulation of catalytic activity | BP | 49 | 5 | 1 | 2.62E-02 | 6.90E-01 |
| GO:0051347 | positive regulation of transferase activity | BP | 85 | 7 | 3 | 2.66E-02 | 3.18E-01 |
| GO:0050877 | nervous system process | BP | 159 | 0 | 8 | 1.00E+00 | 2.79E-02 |
| GO:0048534 | hematopoietic or lymphoid organ development | BP | 86 | 7 | 1 | 2.81E-02 | 8.74E-01 |
| GO:0043549 | regulation of kinase activity | BP | 86 | 7 | 4 | 2.81E-02 | 1.38E-01 |
| GO:0032446 | protein modification by small protein conjugation | BP | 50 | 5 | 0 | 2.83E-02 | 1.00E+00 |
| GO:0009612 | response to mechanical stimulus | BP | 29 | 0 | 3 | 1.00E+00 | 2.84E-02 |
| GO:0048522 | positive regulation of cellular process | BP | 684 | 32 | 17 | 2.86E-02 | 4.26E-01 |
| GO:0006284 | base-excision repair | BP | 8 | 2 | 0 | 2.95E-02 | 1.00E+00 |
| GO:0000076 | DNA replication checkpoint signaling | BP | 8 | 2 | 0 | 2.95E-02 | 1.00E+00 |
| GO:0006271 | DNA strand elongation involved in DNA replication | BP | 8 | 2 | 0 | 2.95E-02 | 1.00E+00 |
| GO:0007019 | microtubule depolymerization | BP | 8 | 2 | 0 | 2.95E-02 | 1.00E+00 |
| GO:2000781 | positive regulation of double-strand break repair | BP | 8 | 2 | 0 | 2.95E-02 | 1.00E+00 |
| GO:1900087 | positive regulation of G1/S transition of mitotic cell cycle | BP | 8 | 2 | 0 | 2.95E-02 | 1.00E+00 |
| GO:0071539 | protein localization to centrosome | BP | 8 | 2 | 0 | 2.95E-02 | 1.00E+00 |
| GO:1905508 | protein localization to microtubule organizing center | BP | 8 | 2 | 0 | 2.95E-02 | 1.00E+00 |
| GO:0006278 | RNA-templated DNA biosynthetic process | BP | 8 | 2 | 0 | 2.95E-02 | 1.00E+00 |
| GO:0007004 | telomere maintenance via telomerase | BP | 8 | 2 | 0 | 2.95E-02 | 1.00E+00 |
| GO:0007631 | feeding behavior | BP | 12 | 0 | 2 | 1.00E+00 | 3.03E-02 |
| GO:0002244 | hematopoietic progenitor cell differentiation | BP | 20 | 3 | 0 | 3.05E-02 | 1.00E+00 |
| GO:0006812 | cation transport | BP | 132 | 0 | 7 | 1.00E+00 | 3.07E-02 |
| GO:0003012 | muscle system process | BP | 52 | 0 | 4 | 1.00E+00 | 3.10E-02 |
| GO:0006813 | potassium ion transport | BP | 30 | 0 | 3 | 1.00E+00 | 3.11E-02 |
| GO:0019953 | sexual reproduction | BP | 88 | 7 | 0 | 3.15E-02 | 1.00E+00 |
| GO:0032886 | regulation of microtubule-based process | BP | 35 | 4 | 0 | 3.17E-02 | 1.00E+00 |
| GO:2001020 | regulation of response to DNA damage stimulus | BP | 35 | 4 | 1 | 3.17E-02 | 5.65E-01 |
| GO:0050794 | regulation of cellular process | BP | ### | 50 | 33 | 3.18E-02 | 8.87E-02 |
| GO:0002520 | immune system development | BP | 89 | 7 | 1 | 3.32E-02 | 8.83E-01 |
| GO:0043269 | regulation of ion transport | BP | 79 | 0 | 5 | 1.00E+00 | 3.45E-02 |
| GO:2000045 | regulation of G1/S transition of mitotic cell cycle | BP | 21 | 3 | 0 | 3.47E-02 | 1.00E+00 |
| GO:0022411 | cellular component disassembly | BP | 36 | 4 | 0 | 3.48E-02 | 1.00E+00 |
| GO:0030263 | apoptotic chromosome condensation | BP | 1 | 1 | 0 | 3.50E-02 | 1.00E+00 |
| GO:0001783 | B cell apoptotic process | BP | 1 | 1 | 0 | 3.50E-02 | 1.00E+00 |
| GO:0006287 | base-excision repair, gap-filling | BP | 1 | 1 | 0 | 3.50E-02 | 1.00E+00 |
| GO:0071681 | cellular response to indole-3-methanol | BP | 1 | 1 | 0 | 3.50E-02 | 1.00E+00 |
| GO:1990705 | cholangiocyte proliferation | BP | 1 | 1 | 0 | 3.50E-02 | 1.00E+00 |
| GO:0051305 | chromosome movement towards spindle pole | BP | 1 | 1 | 0 | 3.50E-02 | 1.00E+00 |
| GO:0036089 | cleavage furrow formation | BP | 1 | 1 | 0 | 3.50E-02 | 1.00E+00 |
| GO:0045004 | DNA replication proofreading | BP | 1 | 1 | 0 | 3.50E-02 | 1.00E+00 |
| GO:0006265 | DNA topological change | BP | 1 | 1 | 0 | 3.50E-02 | 1.00E+00 |
| GO:0040016 | embryonic cleavage | BP | 1 | 1 | 0 | 3.50E-02 | 1.00E+00 |
| GO:0051309 | female meiosis chromosome separation | BP | 1 | 1 | 0 | 3.50E-02 | 1.00E+00 |
| GO:0010994 | free ubiquitin chain polymerization | BP | 1 | 1 | 0 | 3.50E-02 | 1.00E+00 |
| GO:0043988 | histone H3-S28 phosphorylation | BP | 1 | 1 | 0 | 3.50E-02 | 1.00E+00 |
| GO:0034770 | histone H4-K20 methylation | BP | 1 | 1 | 0 | 3.50E-02 | 1.00E+00 |
| GO:0035404 | histone-serine phosphorylation | BP | 1 | 1 | 0 | 3.50E-02 | 1.00E+00 |
| GO:0002418 | immune response to tumor cell | BP | 1 | 1 | 0 | 3.50E-02 | 1.00E+00 |
| GO:0032608 | interferon-beta production | BP | 1 | 1 | 0 | 3.50E-02 | 1.00E+00 |
| GO:0032610 | interleukin-1 alpha production | BP | 1 | 1 | 0 | 3.50E-02 | 1.00E+00 |
| GO:0006272 | leading strand elongation | BP | 1 | 1 | 0 | 3.50E-02 | 1.00E+00 |
| GO:0098765 | meiotic prometaphase I | BP | 1 | 1 | 0 | 3.50E-02 | 1.00E+00 |
| GO:0033316 | meiotic spindle assembly checkpoint signaling | BP | 1 | 1 | 0 | 3.50E-02 | 1.00E+00 |
| GO:0044779 | meiotic spindle checkpoint signaling | BP | 1 | 1 | 0 | 3.50E-02 | 1.00E+00 |
| GO:0051323 | metaphase | BP | 1 | 1 | 0 | 3.50E-02 | 1.00E+00 |
| GO:0044785 | metaphase/anaphase transition of meiotic cell cycle | BP | 1 | 1 | 0 | 3.50E-02 | 1.00E+00 |
| GO:0099606 | microtubule plus-end directed mitotic chromosome migration | BP | 1 | 1 | 0 | 3.50E-02 | 1.00E+00 |
| GO:0098763 | mitotic cell cycle phase | BP | 1 | 1 | 0 | 3.50E-02 | 1.00E+00 |
| GO:0007079 | mitotic chromosome movement towards spindle pole | BP | 1 | 1 | 0 | 3.50E-02 | 1.00E+00 |
| GO:0000087 | mitotic M phase | BP | 1 | 1 | 0 | 3.50E-02 | 1.00E+00 |
| GO:0000089 | mitotic metaphase | BP | 1 | 1 | 0 | 3.50E-02 | 1.00E+00 |
| GO:0002903 | negative regulation of B cell apoptotic process | BP | 1 | 1 | 0 | 3.50E-02 | 1.00E+00 |
| GO:1903126 | negative regulation of centriole-centriole cohesion | BP | 1 | 1 | 0 | 3.50E-02 | 1.00E+00 |
| GO:1901994 | negative regulation of meiotic cell cycle phase transition | BP | 1 | 1 | 0 | 3.50E-02 | 1.00E+00 |
| GO:1905133 | negative regulation of meiotic chromosome separation | BP | 1 | 1 | 0 | 3.50E-02 | 1.00E+00 |
| GO:1902103 | negative regulation of metaphase/anaphase transition of meiotic c | BP | 1 | 1 | 0 | 3.50E-02 | 1.00E+00 |
| GO:0045950 | negative regulation of mitotic recombination | BP | 1 | 1 | 0 | 3.50E-02 | 1.00E+00 |
| GO:0006297 | nucleotide-excision repair, DNA gap filling | BP | 1 | 1 | 0 | 3.50E-02 | 1.00E+00 |
| GO:1905786 | positive regulation of anaphase-promoting complex-dependent c | BP | 1 | 1 | 0 | 3.50E-02 | 1.00E+00 |
| GO:1902425 | positive regulation of attachment of mitotic spindle microtubules | BP | 1 | 1 | 0 | 3.50E-02 | 1.00E+00 |
| GO:1901978 | positive regulation of cell cycle checkpoint | BP | 1 | 1 | 0 | 3.50E-02 | 1.00E+00 |
| GO:0110032 | positive regulation of G2/MI transition of meiotic cell cycle | BP | 1 | 1 | 0 | 3.50E-02 | 1.00E+00 |
| GO:2000617 | positive regulation of histone H3-K9 acetylation | BP | 1 | 1 | 0 | 3.50E-02 | 1.00E+00 |
| GO:0090240 | positive regulation of histone H4 acetylation | BP | 1 | 1 | 0 | 3.50E-02 | 1.00E+00 |
| GO:2000620 | positive regulation of histone H4-K16 acetylation | BP | 1 | 1 | 0 | 3.50E-02 | 1.00E+00 |
| GO:0070512 | positive regulation of histone H4-K20 methylation | BP | 1 | 1 | 0 | 3.50E-02 | 1.00E+00 |
| GO:0002839 | positive regulation of immune response to tumor cell | BP | 1 | 1 | 0 | 3.50E-02 | 1.00E+00 |
| GO:0032728 | positive regulation of interferon-beta production | BP | 1 | 1 | 0 | 3.50E-02 | 1.00E+00 |
| GO:0032730 | positive regulation of interleukin-1 alpha production | BP | 1 | 1 | 0 | 3.50E-02 | 1.00E+00 |
| GO:0032732 | positive regulation of interleukin-1 production | BP | 1 | 1 | 0 | 3.50E-02 | 1.00E+00 |
| GO:1905116 | positive regulation of lateral attachment of mitotic spindle microtu | BP | 1 | 1 | 0 | 3.50E-02 | 1.00E+00 |
| GO:0045654 | positive regulation of megakaryocyte differentiation | BP | 1 | 1 | 0 | 3.50E-02 | 1.00E+00 |
| GO:1901995 | positive regulation of meiotic cell cycle phase transition | BP | 1 | 1 | 0 | 3.50E-02 | 1.00E+00 |
| GO:0090267 | positive regulation of mitotic cell cycle spindle assembly checkpo | BP | 1 | 1 | 0 | 3.50E-02 | 1.00E+00 |
| GO:0032075 | positive regulation of nuclease activity | BP | 1 | 1 | 0 | 3.50E-02 | 1.00E+00 |
| GO:0010845 | positive regulation of reciprocal meiotic recombination | BP | 1 | 1 | 0 | 3.50E-02 | 1.00E+00 |
| GO:0002836 | positive regulation of response to tumor cell | BP | 1 | 1 | 0 | 3.50E-02 | 1.00E+00 |
| GO:0045870 | positive regulation of single stranded viral RNA replication via dou | BP | 1 | 1 | 0 | 3.50E-02 | 1.00E+00 |
| GO:0090232 | positive regulation of spindle checkpoint | BP | 1 | 1 | 0 | 3.50E-02 | 1.00E+00 |
| GO:0090129 | positive regulation of synapse maturation | BP | 1 | 1 | 0 | 3.50E-02 | 1.00E+00 |
| GO:0031915 | positive regulation of synaptic plasticity | BP | 1 | 1 | 0 | 3.50E-02 | 1.00E+00 |
| GO:1905359 | protein localization to meiotic spindle | BP | 1 | 1 | 0 | 3.50E-02 | 1.00E+00 |
| GO:1903096 | protein localization to meiotic spindle midzone | BP | 1 | 1 | 0 | 3.50E-02 | 1.00E+00 |
| GO:0072272 | proximal/distal pattern formation involved in metanephric nephro | BP | 1 | 1 | 0 | 3.50E-02 | 1.00E+00 |
| GO:1905784 | regulation of anaphase-promoting complex-dependent catabolic | BP | 1 | 1 | 0 | 3.50E-02 | 1.00E+00 |
| GO:1902423 | regulation of attachment of mitotic spindle microtubules to kineto | BP | 1 | 1 | 0 | 3.50E-02 | 1.00E+00 |
| GO:0002902 | regulation of B cell apoptotic process | BP | 1 | 1 | 0 | 3.50E-02 | 1.00E+00 |
